# Type III interferons may suppress viral infections by triggering cell death

**DOI:** 10.1101/2024.09.09.612051

**Authors:** Wiktor Prus, Frederic Grabowski, Paulina Koza, Zbigniew Korwek, Maciej Czerkies, Paulina Kaczyńska, Nazanin Amirinejad, Marek Kochańczyk, Tomasz Lipniacki

## Abstract

Type III interferons (IFN-λ1–λ4) are known to limit influenza virus infections in vivo and are non-redundant to type I interferons (IFN-α and IFN-β). Here, we demonstrate that IFN-λ acts through mechanisms that are beyond its ability to induce JAK/STAT signaling and promotes fast cell death in epithelial cells stimulated with poly(I:C). Studying influenza A virus (IAV) and respiratory syncytial virus (RSV) infections in vitro we notice that type I interferons, when provided prior to infection, induce higher STAT1/2 activation and a stronger accumulation of proteins coded by interferon-stimulated genes, and correspondingly suppress both IAV and RSV spread more effectively than type III interferons. The knockout of the IFN-λ receptor (subunit IFNLR1), compared to the knockout of the IFN-β receptor (subunit IFNAR1), only slightly influences levels of STAT1/2 phosphorylation during infection with any of the viruses; However, IFNLR1 knockout results in a greater proportion of IAV-infected cells and higher viral RNA and protein levels. We showed that the ratio of dying to infected cells is lower in IFNLR1-deficient cells compared to wild-type cells, suggesting that IFN-λ promotes the rapid death of IAV-infected cells and thereby limits viral spread. This effect was not observed in RSV-infected cultures, possibly due to the RSV’s ability to suppress host cell death through its nonstructural proteins. Overall, our results reveal a distinct role for IFN-λ in restricting viral infections by triggering rapid death of infected epithelial cells, in contrast to type I interferons, which are the main inducers of JAK/STAT-mediated expression of ISGs, that may be used by IFN-λ to trigger cell death.

## 1 Introduction

The discovery of type III interferons significantly expanded our awareness of the complexity of antiviral mechanisms in innate immunity (Kotenko et al, 2003; Sheppard et al, 2003). This relatively novel cytokine family, consisting of interferons λ1, λ2, λ3, and λ4, was initially believed to be mostly supplementary or even redundant to a much better described and understood family of type I interferons (represented importantly by interferons α and β, but also including several others). This was due to similarities between both cytokine families in the mechanisms of their production, recognition, activation of signaling pathways and downstream expression of target genes. The production of cytokines from both families is triggered by recognition of pathogen particles by the same pathogen recognition receptors and is dependent on activation of transcription factors NF-κB and IRF3/7 (Osterlund et al, 2007). Type I and type III interferons bind to their cognate receptors, IFNAR1/2 and IFNLR1/IL10R2, respectively, to activate the JAK/STAT pathway, which mediates phosphorylation of STAT1 and STAT2 (collectively referred to as STAT1/2) (Mesev et al, 2019). These two transcription factors form a complex with IRF9 and initiate expression of the so-called interferon-stimulated genes (ISGs), which allow the cell to enter a state of increased viral resistance and help activate adaptive immune responses (Schoggins, 2019). These common characteristics of type I and III interferons are generally not shared by type II interferon, IFN-γ, which exhibits a distinct immunomodulatory mode of action and is secreted mostly by specialized immune cells (Schoenborn & Wilson, 2007).

Despite their similarities, accumulating in vivo evidence suggests that the roles of type I and III interferons in response to viral infections are partially divergent, with IFN-λ playing a significant role in containing influenza A virus (IAV) infections. In animal studies, Galani et al, 2017 demonstrated that IFN-λ suppresses the initial viral spread, ensuring optimal viral clearance, while, in the case of high-dose infections, type I interferons are activated later to enhance antiviral and pro-inflammatory responses. Accordingly, IFNLR1-deficient mice display enhanced type I IFN production and lethality. Klinkhammer et al, 2018 showed that in the upper airways IFN-λ, but not type I interferons, confer long-lasting antiviral protection against influenza virus transmission. Hermann et al, 2019 showed, in a murine IAV infection model, that IFN-λ signaling in dendritic cells increases activity of CD8+ T cells. The same effect was also observed for IFN-λ (but not for IFN-β) during SARS-CoV-2 infection, despite IFN-λ signaling minimally affecting the accumulation of ISGs (Solstad et al, 2024). A more significant role of type III interferons compared to type I interferons was also observed in mouse models for other viruses, including rotavirus (Pott et al, 2011) and norovirus (Nice et al, 2015). Studies using in vitro cell line models, such as A549 alveolar epithelium cells, confirmed that type III interferons are produced in response to respiratory viral infections. Secretion of IFN-λ1 in A549 cells was observed by Okabayashi et al, 2011 in response to RSV, while Wei et al, 2014 reported the same during infection with IAV (interestingly, both viruses elicited similar level of IFN-λ1 secretion at MOI=1, about 8 ng/ml for RSV and 6 ng/ml for IAV). Also Wu et al, 2015 showed that secretion of IFN-λ1 and IFN-λ2/3 in A549 cells after IAV (MOI=1, 24h) is several hundred times higher with respect to mock (while for IFN-β the difference with respect to mock is much lower).

In the context of the innate immune system, the non-redundant and significant role of IFN-λ is puzzling, as transcriptome profiling studies have established that the type I and type III signatures are overlapping. Moreover, the ISGs induced by type III IFNs appear to be a subset of those induced by type I IFNs (Lazear et al, 2019 and references therein), though there are findings which report type III interferons uniquely inducing certain genes in specific cell lines: Mmp7, Serpinb1a, and Csprs in mouse IEC cells by IFN-λ1 (Selvakumar et al, 2017); or CXCL10, CXCL11, IFIT3, IFI30, TDRD7 in HECV cells also by IFN-λ1 (Caine et al, 2019). IFN-β was found to be a much more potent inducer of IRF1 expression than IFN-λ, in both mouse IEC cells (Forero et al, 2019) and human BEAS-2B respiratory cells (Novatt et al, 2016,), which may explain the difference in their proinflammatory potential (Panda et al, 2019). However, the major difference between the action of type I and type III interferons seems to lie in the way they affect kinetics and magnitude of target genes’ expression of genes stimulated by them – strong and rapid in the first group, weaker but more sustained in the second group. Wilder et al, 2023 showed (in murine MLE-12 lung epithelial cells) that after 4 h, IFN-β induces over 300 genes, 2.3 times as many as induced by IFN-λ3; however, at 24 h many genes showed a slightly higher induction after IFN-λ3. In their review, Lazear et al, 2019 characterize type I versus type III interferons as having respectively high/low potency, rapid/slower kinetics, being inflammatory/less inflammatory, systemic/restricted by anatomic barriers. Observations indicate that the expression of IFNLR is generally limited to epithelial surfaces of the airways, gut, liver, urinary tract and the blood–brain barrier, being especially high in the airway epithelium, which contrasts with a much more ubiquitous expression of IFNAR (Lazear et al, 2015; Wack et al, 2015). Findings of Forero et al, 2019 support the notion of spatiotemporal differences in action of type I and III interferons, linking low IFNLR abundance on epithelial cells to a deficient STAT activation and inability to promote IRF1 expression. However, such a localized expression of IFNLR predisposes type III interferons to play a major role in combating infections with respiratory viruses, while limiting systemic damage through an excessive inflammatory response. Thus, to some extent, the distinct role of type III interferons observed in animal models can be attributed to differences in their expression and the expression of their cognate receptor.

In this in vitro study, we show that IFN-λ plays a significant and distinct role — compared to IFN-β — in resisting viral infections by promoting faster cell death in infected cells. We have observed that, upon stimulation with poly(I:C), an analog of viral RNA, the resulting cell death largely depends on the presence of the IFNLR receptor, despite concurrent IFN-β–induced overexpression of ISGs. This suggests the existence of a general IFN-λ–centered, antiviral, death-promoting mechanism. To confirm the relevance of our findings to actual respiratory virus infections, we demonstrate that IFN-λ plays a distinct and significant role in limiting infection with IAV — but not with respiratory syncytial virus (RSV) — in respiratory epithelial cells in vitro.

As expected, we observed that prestimulation with IFN-β leads to stronger accumulation of ISGs than prestimulation with IFN-λ1, resulting in a stronger inhibition of both IAV and RSV proliferation. Upon viral infection, phosphorylation of STAT1/2 is dramatically reduced in IFNAR1-deficient but not in IFNLR1-deficient cells, whereas, surprisingly, proliferation of IAV (but not of RSV) is better facilitated by the deficiency of IFNLR1 compared to the deficiency of IFNAR1. Since IFNLR1 has no ligands other than type III interferons (Peterson et al, 2019), this indicates that IFN-λ confers protection against IAV through mechanisms that are beyond its ability to induce JAK/STAT signaling. We observed that the ratio of infected to dead cells is higher in populations of IFNLR1-deficient cells compared to wild-type (WT) cells, indicating that during IAV infection, IFN-λ expedites the death of infected cells, thereby limiting secondary infections and virus spread. Induction of cell death by IFN-λ may rely on accumulation of ISGs due to IFN-β-JAK/STAT signaling; in our earlier study we showed that cell death in response to poly(I:C) is significantly enhanced by IFN-β prestimulation. In the case of infection with RSV, IFNLR1-deficiency has no significant effect, which may be due to inhibition of cell death by RSV nonstructural proteins (Bitko et al, 2007).

## 2 Materials and Methods

### 2.1 Cell lines and cultures

The A549, HEK-293T, HeLa and MDCK cell lines were purchased from the ATCC. A549 cells were cultured in F-12K basal medium supplemented with 10% fetal bovine serum (FBS) and a penicillin-streptomycin antibiotic solution. MDCK, HeLa cells and HEK-293T cells were maintained in Dulbecco’s modified Eagle’s medium (DMEM) supplemented with 10% FBS and a penicillin-streptomycin antibiotic solution. All cell lines were cultured under standard conditions (37°C with 5% CO2) and kept in monolayers up to 90% confluence. The IFNAR1, IFNLR1, and IFNAR1-IFNLR1 knockouts based on the A549 cell line were developed using the CRISPR lentivirus system (Abm). The detailed procedure for obtaining these cell lines have been described in (Czerkies et al, 2022, see Methods section and Supplementary Figure S3 therein).

### 2.2 Virus amplification, isolation and quantification

Respiratory syncytial virus strain A2 and influenza A virus H1N1 strain A/PR/8/34 were purchased from the ATCC and amplified in HeLa and MDCK cells, respectively. Cells were seeded into 225-cm^2^ tissue culture flasks (Falcon) and cultured as described above for 2 to 3 days until they reached 90% confluence. On the day of infection, the following virus growth medium was prepared: DMEM with 2% FBS for RSV or basal minimal essential medium (MEM) with 0.3% bovine serum albumin (BSA) and 1 μg/mL of L-1-tosylamido-2-phenylethyl chloromethyl ketone (TPCK-trypsin) for IAV. The dilutions of virus were prepared in an appropriate medium, with target MOIs of around 0.01 for RSV and 0.05 for IAV. Culture media were removed, and cells were washed once with phosphate-buffered saline (PBS) and overlaid with 10 mL of the inoculum. Virus was allowed to adsorb to cells for 2 h at 37°C, with occasional stirring. Next, additional virus growth medium was added to a total volume of 40 mL per flask. Infected cells were cultured at 37°C until the development of cytopathic effects could be observed in at least 80% of the cells (typically at around 3 days for IAV and 5 days for RSV). Virus-containing culture fluid was then collected and clarified by centrifugation at 3,000 × g at 4°C for 20 min. Next, virus particles were precipitated by adding 50% (wt/vol) polyethylene glycol 6000 (PEG 6000) (Sigma-Aldrich) in NT buffer (150 mM NaCl, 50 mM Tris-HCl [pH 7.5]) to a final concentration of 10% and stirring the mixture gently at 4°C for 90 min. Virus was centrifuged at 3,250 × g at 4°C for 20 min and recentrifuged after the removal of the supernatant to remove the remaining fluid. The pellet was suspended in 1 mL of NT buffer or in 20% sucrose in NT buffer in the case of RSV, aliquoted, and stored at −80°C.

The virus concentration in the collected samples was quantified using an immunofluorescence protocol. HeLa or MDCK cells were seeded onto microscopic coverslips and cultured upon reaching 90 to 100% confluence. Serial dilutions of virus samples were made in virus growth medium in a range of 10−3 to 10−6. After washing with PBS, cells were overlaid in duplicates with diluted virus, which was allowed to adhere for 2 h, with occasional stirring. Afterward, the virus-containing medium was removed, and cells were overlaid with fresh virus growth medium and cultured for 16 h (for IAV) or 24 h (for RSV). Next, cells were washed with PBS and fixed with 4% formaldehyde for 20 min at room temperature. Cells were stained using a standard immunofluorescence protocol with anti-RSV fusion glycoprotein antibody (catalog number ab43812; Abcam) or anti-influenza A virus nucleoprotein antibody (clone C43) (catalog number ab128193; Abcam). Cells containing stained viral proteins were counted using a Leica SP5 confocal microscope. The virus concentration was calculated using the following formula: (average number of infected cells)/(dilution factor × volume containing virus added) = infectious particles/mL. For a given MOI, we observe fewer cells expressing viral proteins in the A549 than in the HeLa and MDCK lines, which are known to be permissive for RSV and IAV infections, respectively.

### 2.3 Compounds and stimulation protocols

Human IFN-β1A and IFN-λ1 were purchased from Thermo Fisher Scientific (catalog numbers PHC4244 and 34-8298-64, respectively) and prepared according to the manufacturer’s instructions. For cell stimulation, interferons were further diluted to the desired concentrations in F-12K medium supplemented with 2% FBS, to prevent the inhibition of viral attachment and entry by serum proteins.

Interferon–virus experiments have been performed as described previously in Czerkies et al, 2022: the cell culture media were exchanged for interferon-containing or control media at time zero and were not removed afterward until the end of the experiment. Appropriately diluted virus was added in small volumes (less than 10 μL) directly into the wells. The even distribution of virus across the cell population was aided by the intermittent rocking of the plate for 2 h.

For poly(I:C) (Sigma-Aldrich) experiments, the medium was changed to antibiotic-free F12K with 2% FBS at least 4 h after seeding, before overnight incubation. Poly(I:C) was delivered to cells by means of lipid-based transfection, using Lipofectamine LTX with Plus reagent (Thermo Fisher Scientific). Modified manufacturer’s protocol optimized for transfection was used; poly(I:C) was mixed with Plus reagent diluted in serum-free F12K and then mixed with F12K-diluted lipofectamine. Liposome-poly(I:C) complexes were allowed to form for 20 minutes before adding them to cells at room temperature. Per 18,000 cells in 270 µl of medium in one well of 96-well plate, 0.1 µl or 0.3 µl poly(I:C) stock solution (1 µg/ml) was added together with 1.8 µl of lipofectamine and 1.8 µl of Plus reagent, diluted in 30 µl of serum-free F12K.

### 2.4 Antibodies

#### 2.4.1 Antibodies for Western blotting

Primary antibodies for Western blotting were anti-phospho-STAT1 (Tyr701) (clone 58D6) (catalog number 9167; Cell Signaling Technologies) (1:1,000), anti-phospho-STAT2 (Tyr690) (clone D3P2P) (catalog number 88410; Cell Signaling Technologies) (1:1,000), anti-RIG-I (clone D14G6) (catalog number 3743; Cell Signaling Technologies) (1:1,000), anti-STAT1 (catalog number 610116; BD Biosciences) (1:1,000), anti-STAT2 (catalog number PAF-ST2; R&D Systems) (1:1,000), anti-PKR (clone B-10) (catalog number sc-6282; Santa Cruz Biotechnology) (1:1,000), anti-OAS1 (clone F-3) (catalog number sc-374656; Santa Cruz Biotechnology) (1:1,000), anti-influenza A virus nucleoprotein (clone C43) (catalog number ab128193; Abcam) (1:1,000), anti-respiratory syncytial virus (clone 2F7) (catalog number ab43812; Abcam) (1:1,000), and hFAB rhodamine anti-glyceraldehyde-3-phosphate dehydrogenase (GAPDH) (catalog number 12004168; Bio-Rad) (1:10,000).

Secondary antibodies for Western blotting were DyLight 800 4× PEG-conjugated goat anti-rabbit IgG(H+L) (catalog number SA5-35571; Thermo Fisher Scientific) (1:10,000), DyLight 800 4× PEG-conjugated goat anti-mouse IgG(H+L) (catalog number SA5-35521; Thermo Fisher Scientific) (1:10,000), StarBright Blue 700-conjugated goat anti-mouse IgG (catalog number 12004159; Bio-Rad) (1:10,000), and horseradish peroxidase (HRP)-conjugated rabbit anti-goat immunoglobulins (catalog number P0449; Agilent) (1:10,000).

#### 2.4.2 Antibodies for immunostaining

Primary antibodies for immunostaining were anti-phospho-STAT1 (Tyr701) (clone 58D6) (catalog number 9167; Cell Signaling Technologies) (1:1,000), anti-respiratory syncytial virus (catalog number ab20745; Abcam) (1:1,000), anti-influenza A virus (catalog number ab20841; Abcam) (1:1,000), anti-influenza A virus nucleoprotein (clone C43) (catalog number ab128193; Abcam) (1:1,000), and anti-human interferon beta (catalog number MAB8142; R&D Systems).

Secondary antibodies for immunostaining were Alexa Fluor 488-conjugated donkey anti-rabbit IgG(H+L) (catalog number A-21206; Thermo Fisher Scientific) (1:1,000), Alexa Fluor 555-conjugated donkey anti-mouse IgG(H+L) (catalog number A-31570; Thermo Fisher Scientific) (1:1,000), and Alexa Fluor 633-conjugated donkey anti-goat IgG(H+L) (catalog number A-21082; Thermo Fisher Scientific) (1:1,000).

### 2.5 RNA isolation

For gene expression analysis experiments, cells were seeded on 12- or 24-well plates (Falcon) at a density of, respectively, 1.0×10^5^ or 1.5×10^5^ cells/well. Upon completed stimulation according to chosen protocol, cells were washed once with PBS and submitted to isolation of total RNA using PureLink RNA Mini Kit (Thermo Fisher Scientific), following manufacturer’s instructions: cells were harvested and vortexed in Lysis Buffer with 2-mercaptoethanol and then vortexed again with one volume of 70% ethanol. Upon transferring to the spin cartridge, cellular RNA content was bound to the column, washed with appropriate buffers and eluted, all by centrifugation at 12,000×g. Eluted RNA in RNase-free water was used immediately for reverse transcription or stored for later use at −80 °C.

### 2.6 Reverse transcription

RNA concentration and quality were determined by measuring UV absorbance of samples diluted 1:100 in 10 mM Tris-HCl (pH 7.5) at 260 and 280 nm, using the Multiskan GO spectrophotometer (Thermo Fisher Scientific). Independently, RIN index for isolated RNA was checked using Agilent 2100 Bioanalyzer. Around 1 µg of RNA was used as a template for reverse transcription, performed using High-Capacity cDNA Reverse Transcription Kit (Thermo Fisher Scientific). Following the manufacturer’s protocol, diluted RNA samples were mixed 1:1 with freshly prepared Master Mix containing RT Buffer, RT Random Primers, dNTP Mix, MultiScribe Reverse Transcriptase, and RNase Inhibitor. Reaction was performed in Mastercycle Gradient thermal cycler (Eppendorf) under the following conditions: 10 min at 25 °C, 120 min at 37 °C, and 5 min at 85°C.

### 2.7 Real-Time Quantitative PCR

Reaction was performed on a QuantStudio 12K Flex Real-Time PCR system with an Array Card block (Life Technologies). In most cases, 750 ng of cDNA from reverse transcription was mixed with reaction Master Mix and loaded onto TaqMan Array Card containing probes and primers for 24 genes (in 2 replicates), including endogenous reference controls (see Source Data S2). Reaction was conducted using QuantStudio “Standard” protocol, with FAM/ROX chemistry. Upon completion, expression of target genes was analyzed based on absolute Ct quantification or classical ΔΔCt method with QuantStudio 12K Flex software, normalized against GAPDH gene expression.

### 2.8 Digital PCR

For viral RNA copy number measurements, cells were seeded onto 24-well plates at a density of 100,000 cells per well. RNA from virus-infected cell populations was isolated and reverse-transcribed as described above. Then, digitalPCR using QuantStudio3D system (Thermo Fisher Scientific) with TaqMan Vi99990011_po and Vi99990014_po gene expression assays have been performed as described previously in Czerkies et al, 2022.

### 2.9 Western blotting

Performed as previously described in Czerkies et al, 2022. Specifically, cells were washed twice with PBS, lysed in Laemmli sample buffer containing dithiothreitol (DTT), and boiled at 95°C for 10 min. Proteins were separated on 4 to 16% Mini-Protean TGX stain-free precast gels using the Mini-Protean Tetra cell electrophoresis system (Bio-Rad). Upon completion, proteins were transferred to a nitrocellulose membrane using Mini-Protean apparatus (400 mA for 50 min). The membrane was rinsed with TBST (Tris-buffered saline [TBS] containing 0.1% Tween 20; Sigma-Aldrich) and blocked for 1 h with 5% BSA–TBS or 5% nonfat dry milk. Subsequently, the membranes were incubated with one of the primary antibodies diluted in 5% BSA–TBS buffer at 4°C overnight. After thorough washing with TBST, the membranes were incubated with secondary antibodies conjugated to a specific fluorochrome (DyLight 800; Thermo Fisher Scientific) or horseradish peroxidase (HRP-conjugated polyclonal anti-mouse/anti-rabbit immunoglobulins; Dako) diluted 1:5,000 in 5% nonfat dry milk–TBST for 1 h at room temperature. The chemiluminescence reaction was developed with the Clarity Western ECL system (Bio-Rad). For GAPDH detection, hFAB rhodamine anti-GAPDH primary antibody (Bio-Rad) was used. Specific protein bands were detected using the ChemiDoc MP imaging system (Bio-Rad).

### 2.10 Immunostaining

For staining of intracellular proteins, cells were seeded onto 12-mm round glass coverslips, which were previously washed in 60% ethanol–40% HCl, thoroughly rinsed with water, and sterilized. After stimulation with interferon or viral infection at the desired time points, cells on the coverslips were washed with PBS and immediately fixed with 4% formaldehyde (20 min at room temperature). Cells were then washed three times with PBS and incubated with 100% cold methanol for 20 min at −20°C. After washing with PBS, coverslips with cells were blocked and permeabilized for 1.5 h with 5% BSA (Sigma-Aldrich) with 0.3% Triton X-100 (Sigma-Aldrich) in PBS at room temperature. After the removal of the blocking solution, coverslips with cells were incubated with primary antibodies diluted in PBS containing 1% BSA and 0.3% Triton X-100 overnight at 4°C. After the cells were washed five times with PBS, the appropriate secondary antibodies conjugated with fluorescent dyes were added, and the cells were incubated for 1 h at room temperature. Subsequently, the cells were washed, and their nuclei were stained with 200 ng/mL 4′,6-diamidino-2-phenylindole (DAPI) (Sigma-Aldrich) for 10 min. After a final wash in MilliQ water, coverslips with stained cells were mounted onto microscope slides with a drop of Vectashield Antifade Mounting Medium (Vector Labs). The cellular sublocalization of stained proteins was observed using a Leica SP5 confocal microscope and Leica Application Suite AF software.

### 2.11 FACS/Cell death assay

Cell death was measured using the annexin V and propidium iodide staining method, according to the protocol described previously in Czerkies et al, 2018. Cells which are both annexin V- and propidium iodide (PI)-negative were considered viable, annexin V-positive and PI-negative cells were considered early apoptotic or pyroptotic; PI-positive cells, irrespective of the outcome of annexin V staining, were considered dying or dead.

### 2.12 Live imaging propidium iodide and CellEvent™ assay

To estimate cell death rates during infection with IAV, RSV and poly(I:C), live imaging of PI intracellular accumulation was performed. One day before the experiment, cell nuclei were stained with Hoechst 33342 (Thermo) diluted 1 µg/ml in PBS for 10 minutes at 37°C. After staining, the cells were washed and seeded onto a µClear 96-well microplate at a density of 18,000 cells per well. The following day, the cells were infected with IAV, RSV or poly(I:C) treated and PI was added. In some experiments, CellEvent™ Caspase-3/7 Detection Reagent (Invitrogen) was used, according to manufacturers’ protocol, by adding a concentrated stock solution to the culture medium. Cells were then observed for up to 48 h, and within these periods a fraction of PI-positive cells was determined. After 12, 24 or 48 h p.i. cell fixation and immunostaining were performed to estimate the number of infected cells. Imaging was carried out on an Operetta CLS (PerkinElmer) using Harmony software.

### 2.13 Image analysis

Confocal images obtained from immunostaining were analyzed using our in-house software (https://pmbm.ippt.pan.pl/software/shuttletracker). Nuclear outlines were detected based on DAPI staining. Outlines of nuclei that were partially out of frame or mitotic were excluded from the analysis; outlines of overlapping nuclei were split based on geometric convexity defects when possible. Cells were classified as either virus-negative or virus-positive based on the sum of the intensities of the pixels inside nuclear outlines. Cells were classified as dead/dying based on the intensities of pixels inside nuclear outlines in the PI channel.

### 2.14 Statistical analysis

Western blot quantifications were first normalized by the corresponding reference GAPDH, then by the maximum value (for each protein and replicate separately). Finally, 0.03 was added to each value, as justified in Grabowski et al, 2023. For statistical analysis, the resulting values were additionally log-transformed. To account for independent normalization of each experimental replicate, in ANOVA tests, we treat replicate number as an additional factor. This approach ensures that rescaling an individual replicate by a constant factor does not change the result of the statistical test. Further details of statistical analysis are provided in Supplementary Methods.

## 3 Results

### 3.1 IFN-λ signaling promotes more rapid cell death after poly(I:C)

We have used viral RNA analog, poly(I:C), to induce cell death in A549 alveolar epithelial cells and observed that cytotoxic effect is highly dependent on the presence of IFNLR receptor. In Fig. 1, we show that in response to poly(I:C) stimulation, WT cells are dying at a markedly faster rate than IFNLR1 KO cells. Furthermore, the effect of IFNLR1 knockout is significant both in experiments without and with IFN-β costimulation. This indicates that IFN-λ signaling (here, triggered by poly(I:C), Salka et al, 2020) contributes to more rapid cell death in a mechanism that is in part independent of its ability to activate STAT1/2, which is relatively weak in comparison to IFN-β. As shown in Figs. S1A and 1B, stimulation with IFN-λ1 (at its saturation concentration of 50 ng/ml) results in lower levels of phosphorylated STAT1/2 (p-STAT1/2) after 1 h, after intermediate times (2 and 6 h), and after longer times (24 and 48 h), compared to stimulation with IFN-β (also at its saturation concentration of 1000 U/ml; saturation concentrations for IFN-λ1 and IFN-β were determined in Fig. S2; all replicates of Western blots used in this study are shown in supplementary Source Data S1).

**Figure 1.**
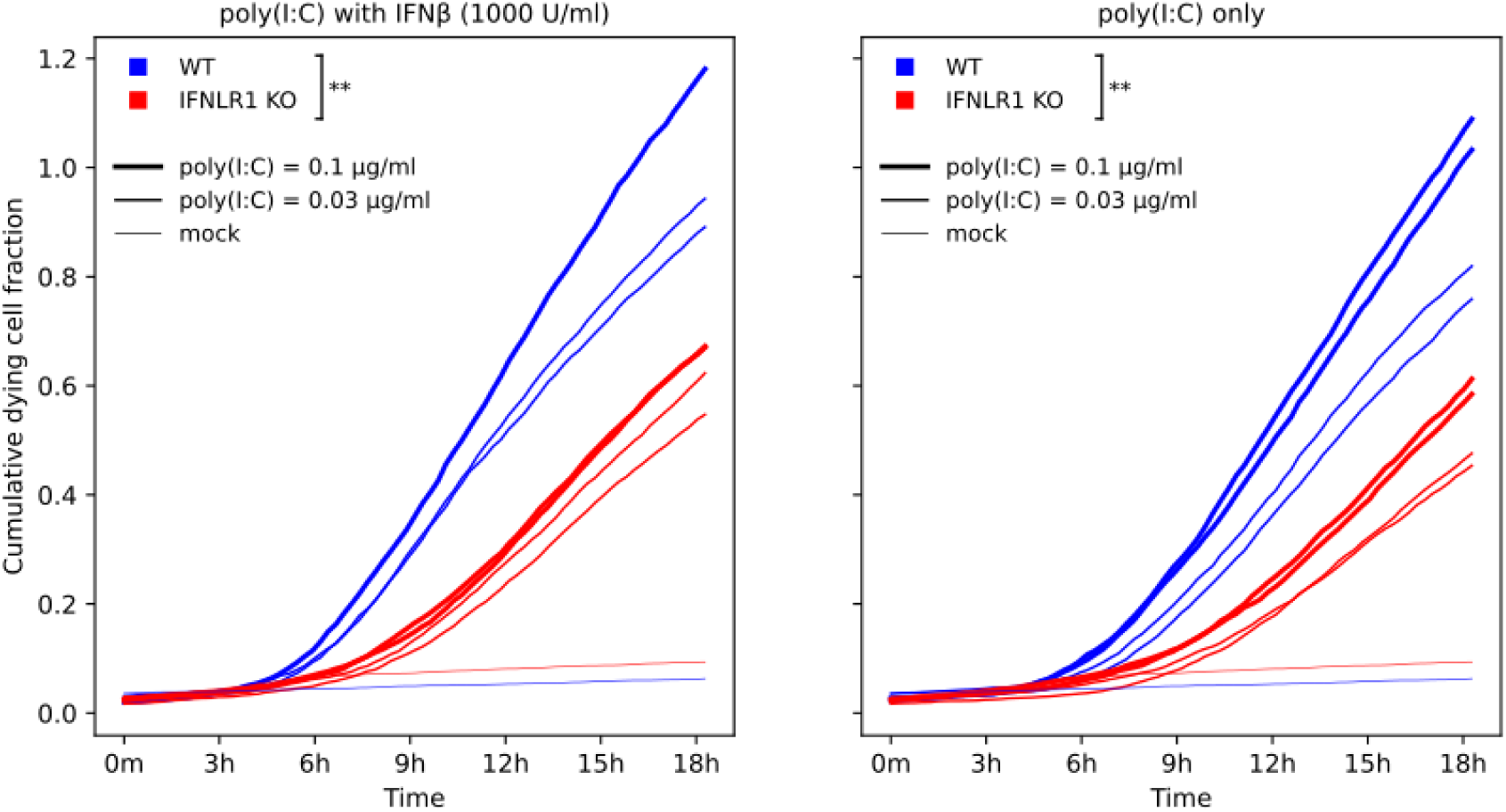
Influence of IFN-λ signaling on poly(I:C) induced cell death. WT and IFNLR1 KO A549 cells were stimulated with the indicated concentrations of poly(I:C) for 18 h, either with or without IFN-β costimulation. One of three experimental replicates is shown; the remaining two are presented in Source Data S1. The dying cells were counted based on the propidium iodide (PI) signal during live microscopy experiments lasting 18 h after adding poly(I:C). The cumulative dying cell fraction is the sum of the fractions of cells that start showing an intracellular PI signal in subsequent hours. The cumulative fraction can increase above one due to cell divisions. After 18 h of poly(I:C), WT cells exhibited significantly higher cumulative dying cell fraction than IFNLR1 KO cells, both with and without IFN-β stimulation, with p < 0.01 (**) determined using a multiple ANOVA test based on three experimental replicates (see Methods for details). The effect is significant at any time point in the last six hours of the experiment, with p < 0.01 in the case of IFN-β costimulation and with p < 0.05 without IFN-β costimulation.

However, it is worth noting that the accumulation of ISG-encoded proteins after IFN-β stimulation seems to positively contribute to poly(I:C)-induced mortality of WT A549 cells, in accordance with earlier observations (Czerkies et al, 2018). This may mean that even if the putative cell death-inducing IFN-λ mechanism does not depend on STAT1/2 signaling, it still makes use of accumulated effector ISG-encoded proteins. As fast cell death is a potent antiviral defense mechanism, preventing secondary infections, we will analyze how IFN-λ signaling (in comparison to IFN-β) regulates the spread of IAV and RSV.

### 3.2 Prestimulation with IFN-β inhibits virus proliferation more markedly than prestimulation with IFN-λ1

As a consequence of generally lower levels of phosphorylated STAT1/2 after stimulation with IFN-λ1 compared to IFN-β, the former leads to a lower accumulation of STAT1, STAT2, and three important ISG-encoded proteins: RIG-I, PKR, and OAS1 (Figs. S1A and S1B). Similarly, mRNA levels are significantly lower after stimulation with IFN-λ1 compared to IFN-β for 15 out of 16 analyzed ISGs important in the regulation of the innate immune response (Fig. S1C). These observations can be tied to the differing efficiency of both interferons in inhibiting infections with respiratory viruses.

We have previously demonstrated that prestimulation with IFN-β strongly reduces IAV replication in host cells (lowering viral RNA count and fraction of infected cells), while the effect on RSV is more moderate (Czerkies et al, 2022, Figs. 2A and 2B therein). The effect of IFN-λ1 prestimulation is much weaker: moderate in case of IAV and negligible in the case of RSV. This is consistent with the weaker activation of ISGs by IFN-λ1 with respect to IFN-β.

**Figure 2.**
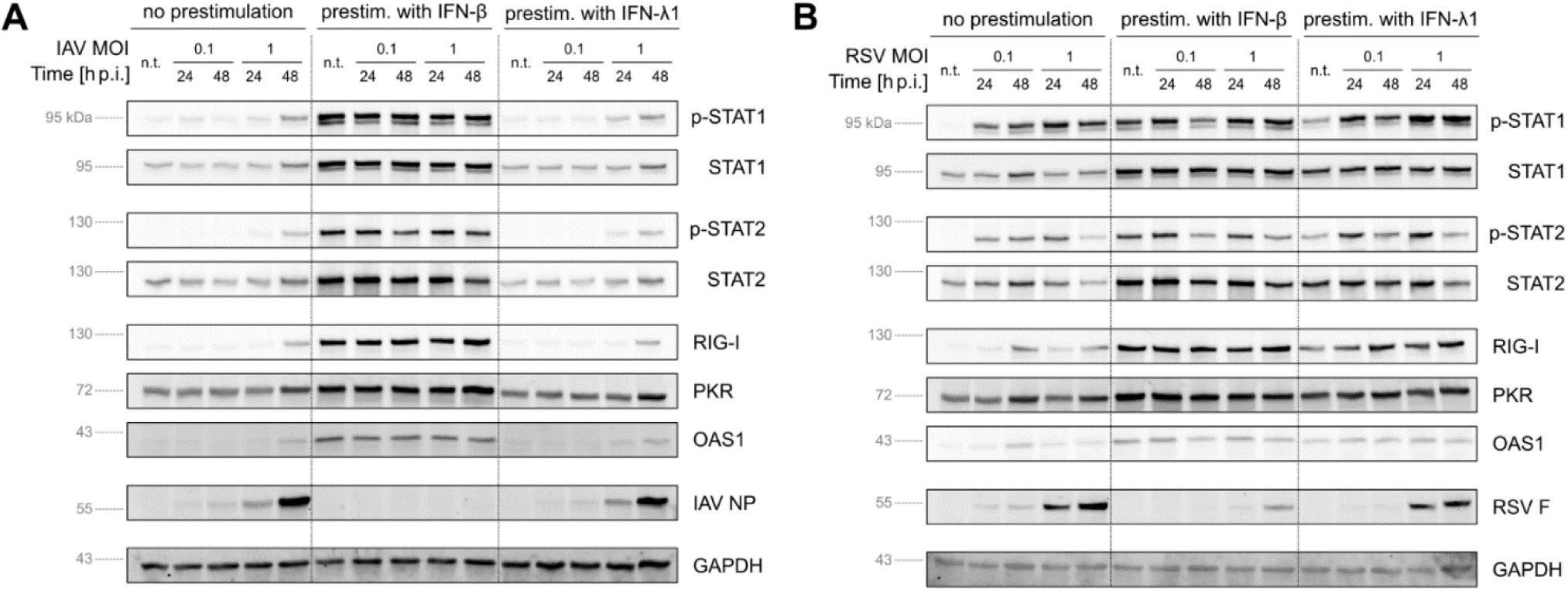
Effect of the prestimulation with IFN-β and IFN-λ1 on STAT1/2 activation, expression of ISGs, and virus proliferation. A A549 cells were prestimulated with either IFN-β (1000 U/ml) or IFN-λ1 (50 ng/ml) (or mock) 24 h before infection with IAV at MOIs of 0.1 or 1 and lysed 24 or 48 h post-infection (p.i.). Label ‘n.t.’ denotes cells not treated with a virus. B A549 cells were prestimulated with IFN-β or IFN-λ1 (or mock) 24 h before infection with RSV at MOIs of 0.1 or 1 and lysed 24 or 48 h p.i. Label ‘n.t.’ denotes cells not treated with a virus. In panels A and B, due to all available 15 lanes being used for samples, the marker lane was omitted. Locations of molecular weight markers were determined based on previous experiments.

Here, we confirm the more substantial effect of prestimulation with IFN-β compared to IFN-λ1 (for infections with either IAV or RSV) by Western blotting. In the case of an infection with IAV (Figs. 2A and S3A), a 24-hour prestimulation with IFN-β (when compared to both mock and IFN-λ1) leads to significantly higher levels of p-STAT1/2, and, consequently, to higher accumulation of STAT1/2 and ISGs observed at 24 and 48 h post infection (p.i.). This results in drastically lower IAV nucleoprotein (NP) levels, below the detection threshold for all combinations of the MOI and time p.i. In the case of an infection with RSV (Figs. 2B and S3B), p-STAT1/2 levels (at 24 and 48 h p.i.) are similar for non-prestimulated and IFN-λ1- or IFN-β-prestimulated cells, indicating that a sufficient amount of IFN-β is produced during infection to trigger STAT1/2 phosphorylation. However, the early accumulation of ISGs, observed in Figs. S1A and S1B after IFN-β prestimulation, resulted in an about 10-fold reduction of RSV fusion (F) protein levels (compared to non-prestimulated cells), not observed in the case of IFN-λ1 prestimulation (Figs. 2B and S3B).

In summary, we have shown that stimulation with IFN-β, compared to IFN-λ1, leads to a higher level of p-STAT1/2, higher expression of ISGs, and, consequently, when applied before viral infection, more effectively reduces virus spread.

### 3.3 IFNLR1 knockout markedly promotes IAV spread while modestly influencing STAT1/2 signaling

In previous experimental protocols, cells responded to a specific interferon provided during prestimulation and, at later stages of the infection, to type I and type III interferons secreted by infected cells. To disentangle the effect of a single type of interferon during viral infection, we employed A549 knock-out cell lines: IFNAR1 KO, IFNLR1 KO, and IFNAR1 & IFNLR1 double KO (further referred to as DKO). We found that despite the fact that in IFNLR1 KO cells the levels of p-STAT1/2, STAT1/2 and ISG are significantly higher (p < 0.01) than in IFNAR1 KO cells at 24 and 48 h p.i., accumulation of IAV NP in IFNLR1 KO cells is also significantly stronger compared to IFNAR1 KO and WT cells (Figs. 3A and 3B). IFNAR1 deficiency does not lead to a significant change in IAV NP accumulation when compared to that in WT cells.

**Figure 3.**
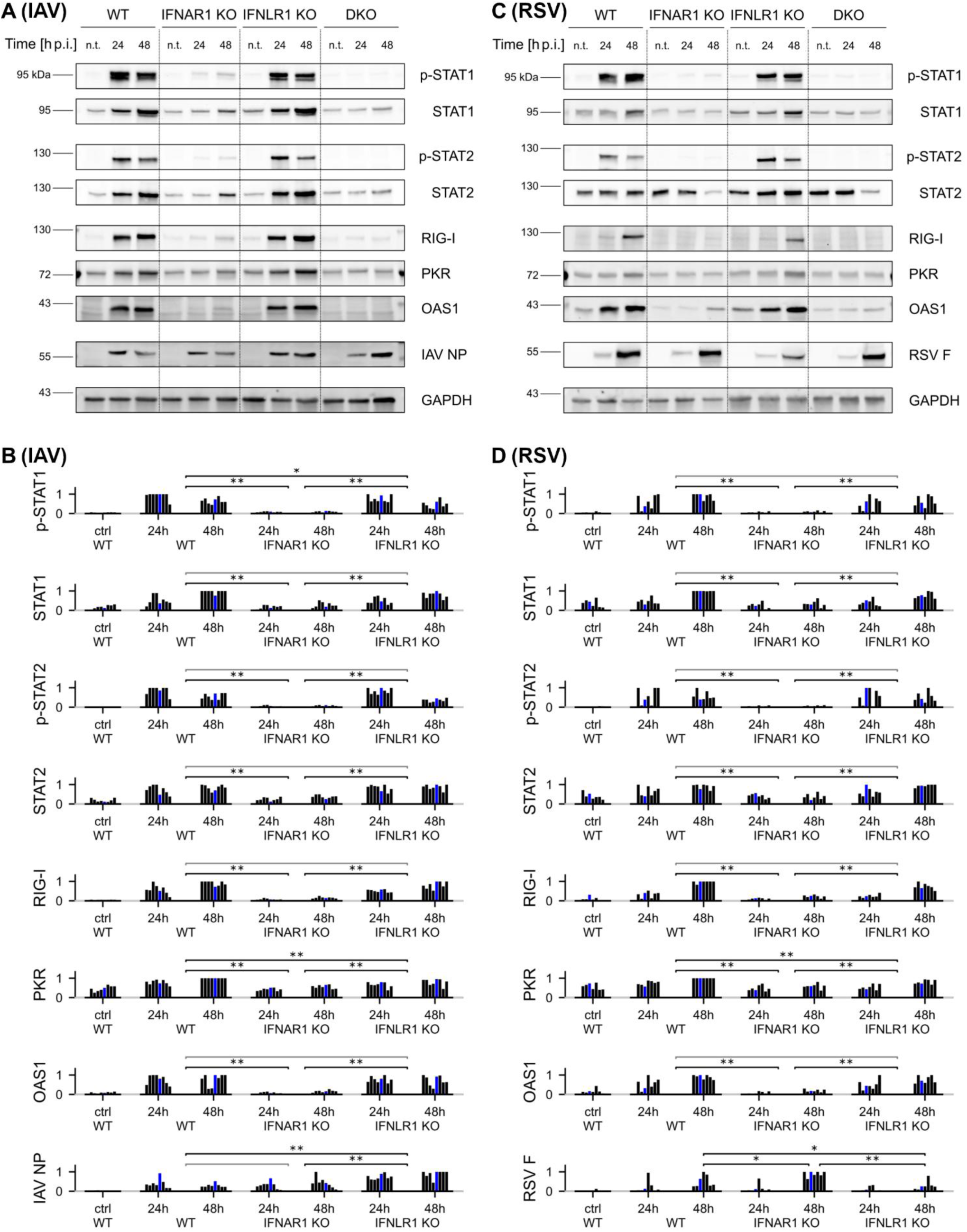
Effect of IFNLR1 and IFNAR1 knockout on IAV and RSV proliferation and activation of immune response. A A549 WT, IFNAR1 KO, IFNLR1 KO, and IFNAR1 & IFNLR1 double KO (DKO) cells were infected with IAV at an MOI of 0.1 and lysed 24 or 48 h p.i. B Quantification of selected bands from blots in panel A and its seven replicates at 24 and 48 h Quantification of the shown blot is marked in blue. C A549 WT, IFNAR1 KO, IFNLR1 KO and DKO cells were infected with RSV at an MOI of 0.1 and lysed 24 or 48 h p.i. D Quantification of selected bands from blots in panel C and its six replicates at 24 and 48 h. Quantification of the shown blot is marked in blue. In panels B and D, differences between cell lines were determined using multiple ANOVA tests on log intensities, corrected with 5% Benjamini-Hochberg false discovery rate. For proteins with significant differences, Tukey post-hoc tests were performed to obtain p values: p < 0.05 (*) and p < 0.01 (**) (see Methods for details). Each blot quantification is normalized to the maximum. The quantification of all cell lines (including DKO) from blots in A and C is shown in Figs. S4 and S5.

This suggests that IFN-λ slows IAV propagation independently of its ability to induce STAT1/2 signaling. This effect is, however, not observed in the case of infection with RSV, where a significantly weaker accumulation of RSV F protein is observed at 48 h p.i. in IFNLR1 KO cells compared to IFNAR1 KO cells (Figs. 3C and 3D). Given the substantial variability in viral protein levels between experimental replicates, Figs. 3B and 3D present the quantification of eight replicates for IAV and seven replicates for RSV (two from our earlier study, Czerkies et al, 2022, and the remaining performed for the current study). Full quantifications, including those for mock infections and DKO cells, can be found in Figs. S4 and S5. The increased spread of IAV in IFNLR1 KO cells, compared to WT cells, is also reflected at the IAV RNA level. Digital PCR measurements (Fig. S6) indicate approximately 3-fold higher accumulation of IAV RNA in IFNLR1 KO cells compared to WT cells (for MOIs of 0.01 and 0.1; 24 to 96 h p.i.), in good agreement with the Western blot results. Using the new IAV batch, we confirmed elevated expression of IAV proteins in IFNLR1 KO cells with respect to WT and IFNAR1 KO cells in the supplementary experiment, in which we measured the level of NS1 (nonstructural) IAV protein in addition to NP (Fig. S7).

Finally, higher levels of IAV mRNA and NP protein observed in IFNLR1 KO cells (but not in IFNAR1 KO cells) with respect to WT cells correlate with a higher fraction of infected cells observed in immunostaining images, shown in Fig. 4A and 4C. We note that p-STAT1 in IFNAR1 KO is almost entirely absent, while p-STAT1 in IFNLR1 KO is not distinguishable from that in WT. Overall, we showed that IFNLR1 deficiency increases IAV NP, IAV RNA, and the fraction of infected cells (with respect to WT), without substantially changing p-STAT1/2 levels and accumulation of ISG-encoded proteins.

**Figure 4.**
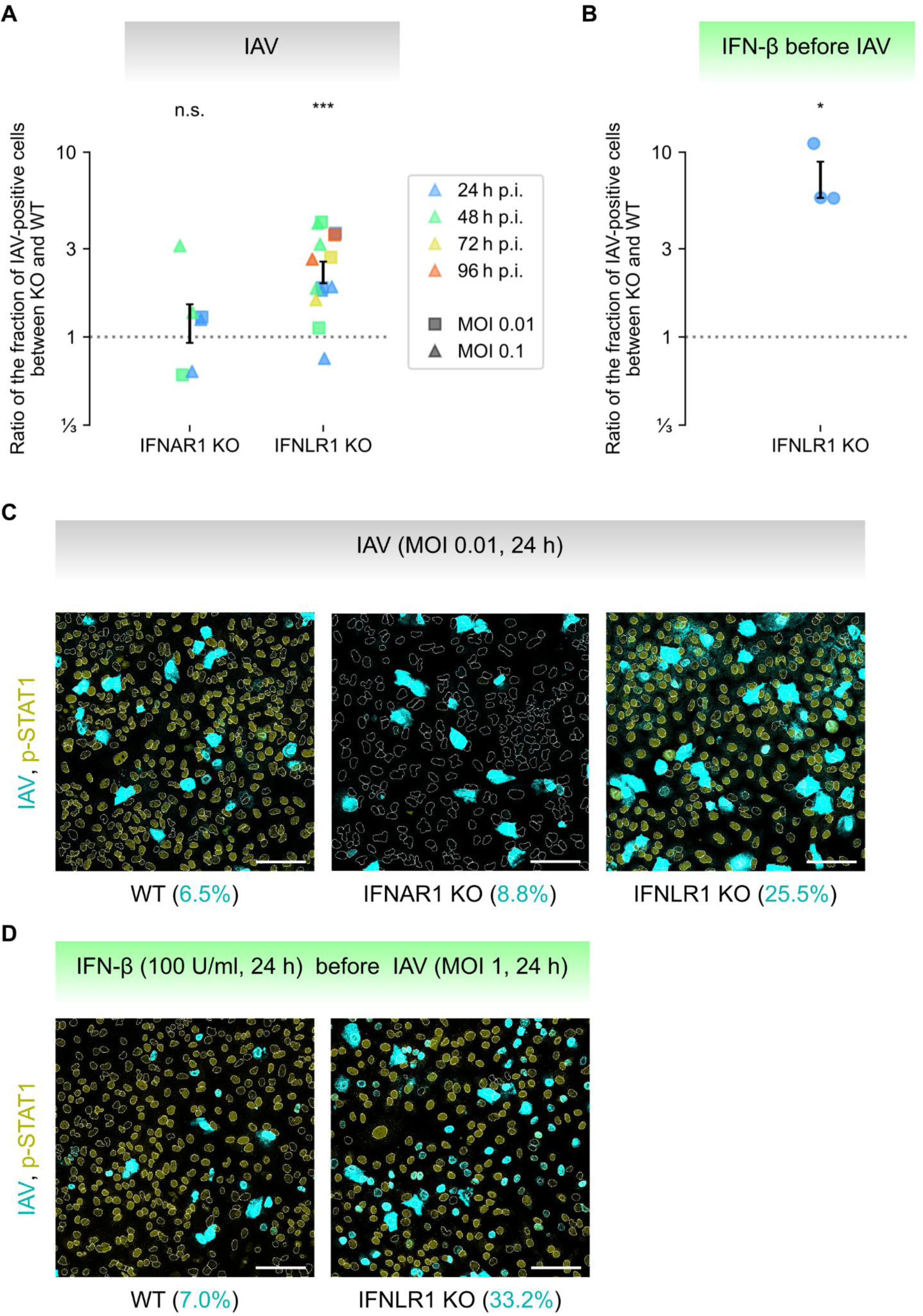
Effect of IFNLR1 and IFNAR1 knockout on IAV propagation and activation of immune response. A Fraction of IAV-positive A549 IFNAR1 KO and IFNLR1 KO cells with respect to the fraction observed for WT. Quantification of three experimental replicates, p < 0.001 (***) according to the one-sample two-tailed Student t-test on log ratios (between KO and WT cell lines) for 24 and 48 h p.i. and MOIs of 0.01 and 0.1 considered jointly. B Fraction of IAV positive A549 IFNAR1 KO cells with respect to the fraction observed for WT. Cells were prestimulated with IFN-β at a concentration of 100 U/ml 24 h before infection with IAV at an MOI of 1. Cells were fixed 24 h p.i. Quantification of three experimental replicates, p < 0.05 (*),one-sample two-tailed Student t-test on log ratios (between IFNLR1 KO and WT cell lines). C Representative fields of view (FOV) showing immunostaining for IAV and p-STAT1 in A549 WT, IFNAR1 KO and IFNLR1 KO cells fixed 24 h p.i. with IAV at an MOI of 0.01. Percentages of infected cells in shown FOVs are given in cyan. Scale bar, 100 μm. All immunostaining images are available at Zenodo (see Data availability). Quantifications shown in Panel A are based on averaging over multiple FOVs per experimental replicate. D Representative fields of view (FOV) showing immunostaining for IAV and p-STAT1 in A549 WT and IFNAR1 KO cells prestimulated with IFN-β at a concentration of 100 U/ml 24 h before infection with IAV at an MOI of 1, and fixed 24 h p.i. Percentages of infected cells in shown FOVs are given in cyan. Scale bar, 100 μm. All immunostaining images are available at Zenodo (see Data availability). Quantifications shown in Panel B are based on averaging over multiple FOVs per experimental replicate.

### 3.4 Functions of IFN-λ extend beyond induction of ISGs

As shown in the previous section, during IAV infection, lack of IFNLR1 does not significantly hinder the production of the considered ISG-encoded proteins but leads to significantly higher levels of viral RNA, viral proteins, and a greater fraction of infected cells. To exclude the effect of ISG production resulting from IFNLR1 activation, we performed an immunostaining experiment comparing the fraction of IAV-positive cells in WT and IFNLR1 KO cell populations prestimulated with 100 U/ml IFN-β for 24 hours before infection with IAV (MOI = 1) and lysed 24 h p.i. (Fig. 4B and 4D). We used a high MOI and a lower IFN-β concentration, because IFN-β prestimulation substantially inhibits IAV spread. We observed that the fraction of infected cells was about 6-fold higher in IFNLR1 KO cells compared to WT cells. This implies that IFN-λ signaling suppresses IAV spread even more effectively when expression of ISGs is already ensured by prior IFN-β prestimulation. This may suggest that IFN-λ suppresses IAV spread using ISGs which accumulate due to IFN-β stimulation.

In summary, we demonstrated that IFN-λ signaling hinders viral spread during an infection with IAV (but not with RSV), and this effect is independent of its regulation of expression of ISGs. In the following sections, we argue that IFN-λ signaling is linked to cell death (either apoptosis or pyroptosis), which is crucial for eliminating infected cells.

### 3.5 Measuring an increase in the cell death rate

We postulate that IAV-infected IFNLR1 KO cells are more resistant to cell death compared to WT cells, allowing for faster virus proliferation. The natural inclination is to compare the number of cell deaths in IFNLR1 KO and WT cell populations infected with IAV. However, a change in the cell death rate affects the total number of infected cells, rendering a direct comparison impossible. Instead, the ratio of cell death events to the number of infected cells is a much more reliable indicator of an increased cell death rate. We clarify this by adapting the susceptible, infected, removed (SIR) model (Eqs. 1–3) to represent viral infection of the cell population, interpreting the removed cells (R) as dead. The kinetics are given by the following equations:

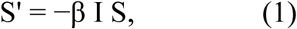

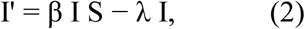

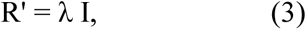

where ’ denotes differentiation with respect to time, β denotes the infection rate, and λ denotes the death rate. In Fig. 5, we show time trajectories of observables R, I, and their ratios for varied values of parameters β and λ for an MOI of 0.1 (see Fig. S8 for MOIs of 0.01 and 0.3). The fraction of dead cells R can increase with λ (at early times) or decrease with λ (at later times), and is generally not monotonic with respect to this parameter (Fig. 5A). This occurs because, at higher λ, the infection progresses more slowly, leading to an overall lower fraction of infected cells (Fig. 5B). However, since the fraction of infected cells (I) is also strongly affected by the infection rate β (Fig. 5C), it cannot be used alone to distinguish between a decrease in mortality and an increase in infectivity. In contrast, the ratio R / I changes strongly and monotonically with λ (Fig. 5D), but weakly and not monotonically with β (Fig. 5E).

**Figure 5.**
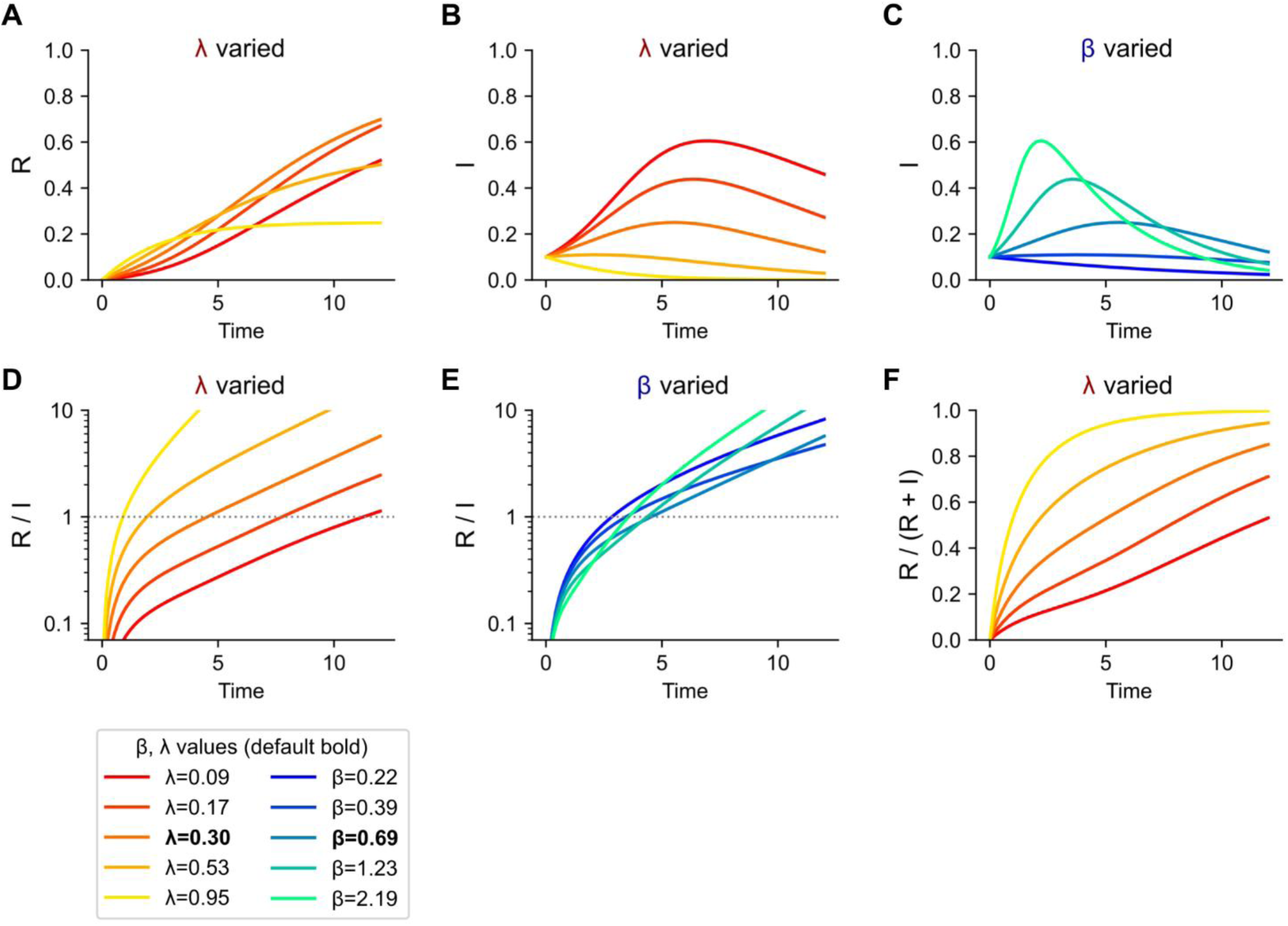
SIR model simulation. Influence of death and infection rates on virus propagation and cell death. A, B Fraction of dead (R, shown on panel A) or infected (I, shown on B) cells as a function of time and the death rate λ. C Fraction of infected cells (I) as a function of time and the infection rate β. D, E Ratio of dead (R) to infected (I) fractions as a function of time and the death rate λ (D) or the infection rate β (E). F Ratio of dead (R) to joint dead and infected (R + I) fractions as a function of time and the death rate λ. In all panels, I(0) = 0.1, corresponding to MOI = 0.1. Unless specified otherwise, β = 0.69 and λ = 0.3. The default value of β was selected so that the initial doubling time is 1.

In summary, a simultaneous increase in the virus-infected cell fraction (I) and a decrease in the ratio of dead to virus-infected cells (R / I) indicates a decrease in the death rate λ. An important detail is that when experimentally measuring the proportion of infected cells (by immunostaining) or viral proteins or RNA produced by infected cells (by digital PCR or Western blot), we account for live infected cells but also some already dead infected cells. Fortunately, the monotonicity of the ratio R / I with respect to λ implies monotonicity of the ratio R/(I + α R) for any α ≥ 0. Furthermore, even when α = 1, the ratio R/(I + R) still grows substantially with λ (Fig. 5F).

### 3.6 IFN-λ signaling suppresses viral spread by promoting cell death

In Fig. 6A we show that the ratio of cell deaths to accumulated IAV NP is higher in WT cells than in IFNLR1 KO cells, which is equivalent to showing that the ratio of dying to infected cells is higher in the case of WT than IFNLR1 KO cells. The left column of Fig. 6A presents results from Western blots and shows the ratio of IAV NP between IFNLR1 KO and WT cells at 24 h p.i. and 48 h p.i. The amount of IAV NP increases significantly in IFNLR1 KO cells, consistent with previous experiments (compare with Fig. 3B), despite the high variability observed between experiments. The right column of Fig. 6A shows the ratio of the fraction of cell deaths (measured by FACS) between IFNLR1 KO and WT cells for the same experimental protocol. On average, the ratios are only slightly higher than 1, indicating that the fraction of dead cells in IFNLR1 KO cell populations is comparable to that in WT cell populations. Since the ratio of viral proteins is higher than the ratio of cell deaths, we may conclude that the overall death rate is higher in WT than in IFNLR1 KO cells.

**Figure 6.**
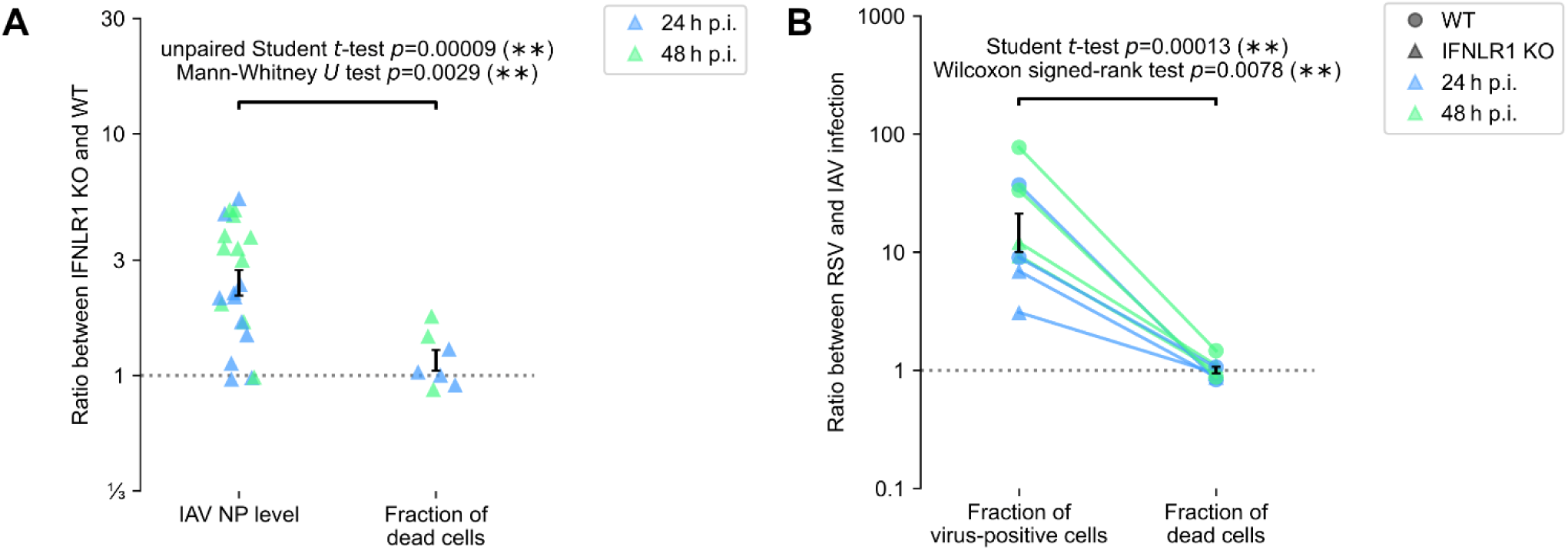
Influence of IFNLR1 deficiency and virus type on infected and dead cell fractions. A Left: ratio of the IAV NP level between IFNLR1 KO and WT cells – calculated based on Western blots, using cells lysed 24 and 48 h p.i. at an MOI of 0.1. Right: ratio of the fraction of dying/dead cells 24 and 48 h p.i. between IFNLR1 KO and WT cells – calculated based on cell death assay using annexin V and propidium iodide staining (see Methods). Statistical significance was determined by performing an unpaired two-tailed Student t-test and a Mann-Whitney U test on log ratios (left versus right). B Left: ratio of RSV-positive to IAV-positive cell fraction at 24 or 48 h p.i. at an MOI of 0.1 – calculated based on immunostaining images (with staining against RSV and IAV, respectively). Right: ratio of the fraction of dead/dying cells after infection with RSV to an analogous fraction after infection with IAV. The dead/dying cells were counted based on the propidium iodide signal during live microscopy experiments lasting 24 or 48 h p.i. The same cells were fixed and immunostained to determine the fraction of virus-infected cells used in the left part of the panel. Statistical significance was determined by performing a paired two-tailed Student t-test and a two-tailed Wilcoxon signed-rank test on log ratio-of-ratios (left / right, see Methods).

Prompted by an anonymous reviewer, we the considered hypothesis that higher level of IAV NP observed in IFNLR1 KO cells (with respect to WT) are due to these cells being more prone to infection. To address this hypothesis, we investigate the progression of infection in both cells lines. In Fig. S9B, we observe that at 12 h p.i. the fraction of cells expressing IAV proteins does not exceed 1% in both cells lines, while at 24 and 48 h p.i the fraction is markedly higher for IFNLR1 KO cells, Fig. S9A. In Western blots in Fig. S9B we see that IAV NP starts accumulating at 12 h p.i., with the maximum at 24 or 48 h p.i. Based on data in Figs. S9A and S9B alone it is hard to confirm or reject the hypothesis. When we combine data from all 14 Western blotting experiments in which expression of IAV NP was determined in WT and IFNLR1 KO cells at 24 and 48 h p.i., we see that, on average, infection progresses faster in INFLR1 KO cells, Fig. S9C. While this conclusion is statistically sound, the variability between experimental replicates is high. Faster infection progression in IFNLR1 KO cells is in line with the lower death rate in these cells, as they have more time to multiply the virus.

RSV provides a relevant comparison, as it is known that its NS1 protein effectively blocks cell death, promoting viral spread (Bitko et al, 2007). To compare cell death rates during infection with IAV and RSV, we measured fractions of cells displaying propidium iodide signal during 24 and 48-hour-long infections. Next, the cells were fixed, and fractions of virus-positive cells at 24 and 48 hours were measured based on immunostaining images. In Fig. 6B, we show that cell death fractions for RSV and IAV are similar, but the infected cell fraction is much higher for infection with RSV than for infection with IAV. Consequently, this indicates a higher death rate for IAV infections compared to RSV infections. The effect is observed for both WT and IFNLR1 KO cells. This supports our reasoning that an inhibition of cell death can be manifested by an increase of the fraction of infected cells rather than a decrease in the fraction of dead cells, suggesting that IFN-λ signaling suppresses viral spread by promoting cell death.

### 3.7 IAV – infected cells show caspase 3/7 activation and either apoptotic or pyroptotic morphology

During infection with IAV, A549 cells detected as dying by PI signal also show CellEvent^TM^ (CE) signal, which indicates caspase 3/7 activation (Fig. 7). Activation of caspase 3 may indicate both apoptosis and gasdermin E-mediated pyroptosis (Jiang et al, 2020). In the context of infection with IAV, it was demonstrated that caspase 3 cleaves Gasdermin D, to yield its inactive N-terminal p20 fragment (Wang et al, 2021), and cleaves gasdermin E, producing 35 kDa N-terminal fragments, which then mediate pyroptosis by forming pores in the cell membrane (Rogers et al, 2017). Moreover, activated gasdermin E permeabilizes the mitochondrial membrane, releasing cytochrome c, further augmenting caspase 3 activation (Rogers et al, 2017). This is in line with our observation that cells with outstanding caspase 3/7 activation exhibit pyroptotic morphology (Fig. 7A and 7B). Specifically, in Fig. 7B, we highlighted the formation of the characteristic pyroptotic bubbles, clearly visible in brightfield. Regarding the intensity of caspase 3/7 activation in dying cells, we have not observed any significant differences between WT and IFNLR1 KO cells (Fig. 7C and two other experimental replicates in Source Data S1), but the matter needs further study.

**Figure 7.**
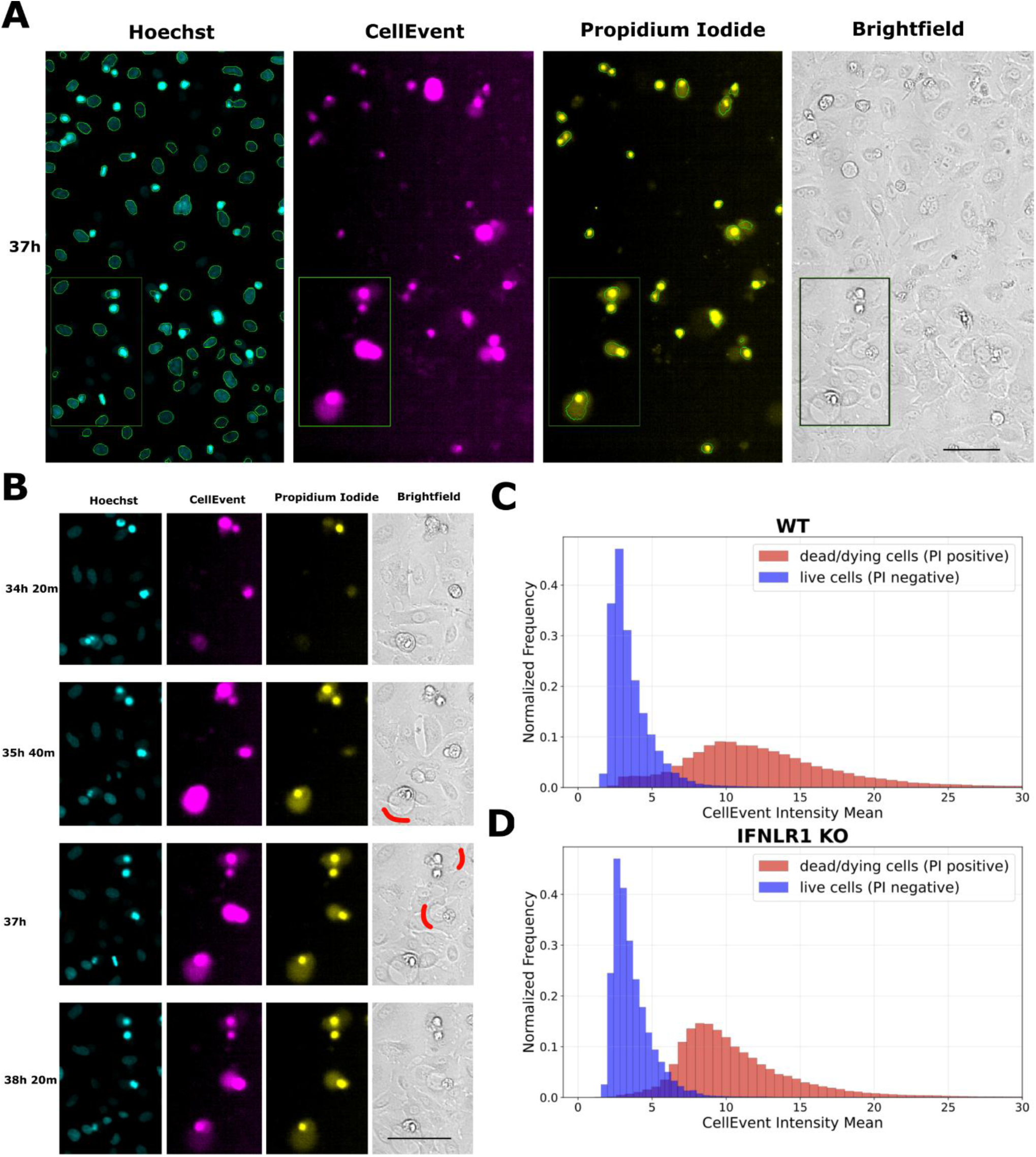
Activation of caspase 3/7 in dying cells with apoptotic and pyroptotic morphology. A Snapshot (at 37 h p.i.) from a live confocal experiment with A549 WT cells infected with IAV at an MOI of 0.1. Cell nuclei were detected based on Hoechst, and dying cells were detected based on propidium iodide (PI), as shown by respective contours. Nearly all dying cells show a CellEvent^TM^ (CE) signal indicating caspase 3/7 activation, and a few cells show a CE signal before PI. A fraction of cells with a strong CE signal show pyroptotic morphology visible on a bright field image. Scale bar 100 μm. B Series of snapshots of a rectangular area shown in Panel A, visualizing pyroptotic morphology of dying cells. Pyroptotic bubbles are highlighted by a red bold line on the bright field image. Scale bar 100 μm. C & D Histograms showing caspase 3/7 activation (CE signal) in dead/dying and live WT (panel C) and IFNLR1 KO (panel D) cells. Histograms were obtained based on snapshots from a 48-hour-long experiment. Corresponding histograms, obtained from two other experimental replicates, are provided in Source Data S1.

## 4 Discussion

Previous in vivo studies demonstrated pronounced role of type III interferons in limiting IAV and some other viral infections (Galani et al, 2017; Klinkhammer et al, 2018; Solstad et al, 2024; Pott et al, 2011; Nice et al, 2015), and current study confirms this role in epithelial cell populations in vitro. The strong effect of type III interferons is puzzling as they are generally weaker inducers of STAT signaling than type I interferons (Lazear et al, 2019 and references therein). Here, we showed that in response to viral RNA analog, poly(I:C), WT A549 cells exhibit faster death than IFNLR1-deficient cells, indicating that IFN-λ may have pro-death role in the context of viral infections. Shortening of time span between cell infection and its apoptosis (or pyroptosis) is critical, as it reduces probability of infecting neighboring cells. The prodeath effect of IFN-λ after poly(I:C) was observed both in presence and absence of IFN-β costimulation, ruling out possibility that IFN-λ promotes cell death by activation of STAT1/2.

Next, using WT, IFNLR1 KO, IFNAR1 KO, and IFNLR1–IFNAR1 DKO A549 cells, we demonstrated divergent roles of type I and type III interferons in control of IAV and RSV proliferation. We showed that (pre)stimulation with IFN-β leads to a strong increase of p-STAT1/2 and STAT1/2 levels, high accumulation of ISGs, and, through those mechanisms, a pronounced reduction of proliferation of IAV as well as RSV. Prestimulation with IFN-λ (compared to IFN-β) results in lower levels of p-STAT1/2 and STAT1/2, lower accumulation of ISGs, and only moderately reduces proliferation of IAV, and does not reduce proliferation of RSV. Next, we showed that in IAV- or RSV-infected IFNAR1 KO cells, p-STAT1/2, STAT1/2 and ISG levels are pronouncedly reduced with respect to those in WT and IFNLR1 KO cells. However, IAV proliferates significantly faster in IFNLR1 KO cells than in WT or IFNAR1 KO cells. This indicates that IFN-λ suppresses IAV proliferation independently of inducing JAK/STAT signaling. Inhibition of IAV spread through IFN-λ may, however, rely on ISGs, which are strongly upregulated by secreted IFN-β.

We note the possibility that IFNLR1 inhibits IAV proliferation independently of type III interferons; however, mouse studies indicate that type III interferon deficiency has the same consequences as IFNLR1 deficiency during infections with norovirus, reovirus, and influenza virus (Peterson et al, 2019). The faster viral propagation in IFNLR1 KO cells is not observed for RSV. (3) Finally, we showed that the absence of IFN-λ signaling significantly increases the level of IAV protein and the fraction of infected cells, without an increase in the fraction of dead cells. With the help of a simple mathematical model, we interpret this observation as the ability of IFN-λ to promote early death in cells infected with IAV. Additionally, we showed that for infection with IAV, the ratio of dead to infected cells is higher than for infection with RSV. This is in line with earlier observations that NS proteins of RSV very effectively suppress cell death in infected cells, promoting virus proliferation (Bitko et al, 2007). The viruses developed different evolutionary strategies: IAV primarily impairs interferon production, whereas RSV rely on evading interferon effects. This difference has been previously observed during our research on the cross-protection effect between RSV and IAV, as described in Czerkies et al. 2022. This could explain why the effect of IFN-λ on RSV spread appears negligible.

Although divergent roles of type I and type III interferons have been highlighted in numerous studies, reports of a non-transcriptional role, especially proapoptotic (or pyroptotic), of IFN-λ are scarce. In two studies not related to infections, it was shown that in intestinal cells type III interferons (compared to type I interferons) are a more potent inducers of apoptosis (Li et al, 2008) and pyroptosis (preprint: Sposito et al, 2022 and Jena et al, 2024). Broggi et al (Broggi et al, 2017), in a study on intestinal inflammation, showed that IFN-λ regulates functions of neutrophils (known to have high expression of IFNLR) independently of transcription, and that treatment with IFN-λ induced in these cells a reduction in the phosphorylation of AKT at Ser473 and Thr308, which could make these cells more prone to cell death. Earlier, Denisova et al (Denisova et al, 2014) showed that inhibiting AKT suppresses proliferation of IAV in A549 cells. Major et al (Major et al, 2020) found that IFN signaling (especially IFN-λ) hampers lung repair by inducing p53, which activates antiproliferative and cell death pathways. Transcriptional regulation of p53 by interferons was also demonstrated in an earlier study by Takaoka et al (Takaoka et al, 2003). Finally, activation of antiviral state by IFN-λ, but not IFN-β, was found to be dependent on the MAPK pathway in human IEC cells, providing another putative link from IFNLR activation to cellular death mechanisms (Pervolaraki et al., 2017). In the context of viral infections, a straightforward demonstration that IFN-λ increases the cell death rate is challenging because a higher cell death rate does not necessarily result in a greater fraction of dead cells, as it also suppresses viral propagation. We suspect that this may partially explain why this phenomenon was not reported earlier.

Our findings confirm that the role of IFN-β is to promote an antiviral state in not yet infected bystander cells, which increases their resistance to infection and/or slows down virus replication once they are infected. In terms of the SIR model, the effect of IFN-β can be seen as a reduction of the infection rate coefficient β, see Eq. (2). The role of IFN-λ (apart from STAT signaling) would be to promote cell death, and thus the removal of infected cells. In the SIR model, this corresponds to an increase of the death rate coefficient λ, see also Eq. (2). In this picture, type III interferons are more universal than type I interferons, being able to both induce some STAT signaling and trigger cell death. Finally, it should be stressed that our data suggest the possibility that IFN-λ triggers cell death using ISGs that are accumulated (mostly) due to IFN-β signaling. This is in line with the observation that the effect of IFNLR1 deficiency is strongest when cells are already prestimulated with IFN-β, and our earlier study (Czerkies et al, 2018) which showed that the death rate after poly(I:C) stimulation is much higher in IFN-β-prestimulated cells.

## 6 Author Contributions

WP: Conceptualization, Methodology, Investigation, Writing – original draft, Writing – review & editing

FG: Conceptualization, Formal analysis, Data curation, Writing – original draft, Writing – review & editing, Visualization

P Koza: Conceptualization, Methodology, Investigation, Writing – review & editing

ZK: Conceptualization, Methodology, Investigation, Writing – original draft, Writing – review & editing

MC: Conceptualization, Methodology, Investigation, Writing – original draft, Writing – review & editing

P Kaczyńska: Conceptualization, Visualization, Data curation, Writing – review & editing

NA: Conceptualization, Investigation, Writing – review & editing

MK: Conceptualization, Data curation, Writing – original draft, Writing – review & editing, Software, Supervision

TL: Conceptualization, Writing – original draft, Writing – review & editing, Supervision

## 8 Data Availability Statement

Images of all Western blots performed for this study are available in the supplementary file Source Data S1. Quantifications of Western blots, RT-PCR, digital PCR, FACS, imaging data, and codes to generate figures are available in the supplementary file Source Data S2. The raw imaging dataset (except live confocal images for Fig. 1 and Fig. 7) and quantification for Fig. 7C and Fig. 7D are available at Zenodo (https://doi.org/10.5281/zenodo.17122542). The set of confocal images for Fig. 1 and Fig. 7 (too large to be deposited on Zenodo) is available upon request.

