## Supplementary Materials for "Type III interferons may suppress viral infections by triggering cell death"

### *Supplementary Material*

#### 1 Supplementary Figures and Tables

##### 1.1 Supplementary Figures

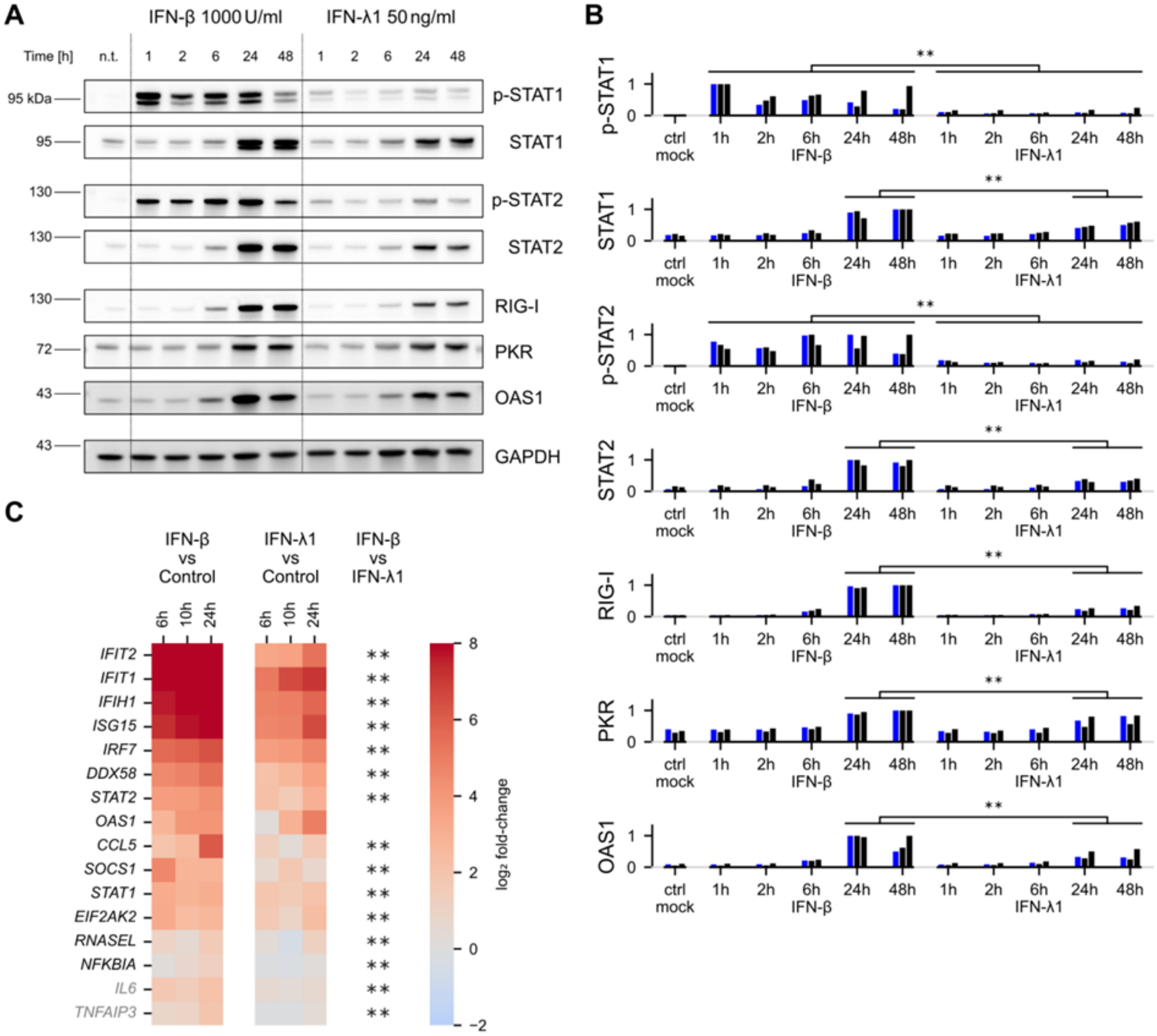


**Figure S1. STAT1/2 activation and accumulation of ISGs after stimulation with IFN-β and IFN-λ1.**

A A549 cells were stimulated with IFN-β (1000 U/ml) or IFN-λ1 (50 ng/ml) for 1, 2, 6, 24, or 48 h before lysis; p-STAT1, STAT1 phosphorylated at Tyr701; p-STAT2, STAT2 phosphorylated at Tyr690.

B Quantification of three experimental replicates. Blue bars indicate the experimental replicate shown in panel A. Each blot quantification is normalized to the maximum. The significances $p < 0.01$

(**) were determined using multiple ANOVA on log intensities corrected using 5% Benjamini–Hochberg false discovery rate (see Methods for details).

C Gene expression of chosen ISGs as well as IL6 and TNFAIP3 after 6, 10, and 24 h stimulation with either IFN-β (1000 U/ml) or IFN-λ1 (50 ng/ml), analyzed by RT-PCR. The averaged log2 difference (based on 3 experimental replicates) with respect to control is shown. Asterisks indicate that expression of a given gene after stimulation with IFN-β is significantly different (higher) than after IFN-λ1 at $p < 0.01$ (**); multiple ANOVA on log2 differences corrected using 5% Benjamini–Hochberg false discovery rate (see Methods for details).

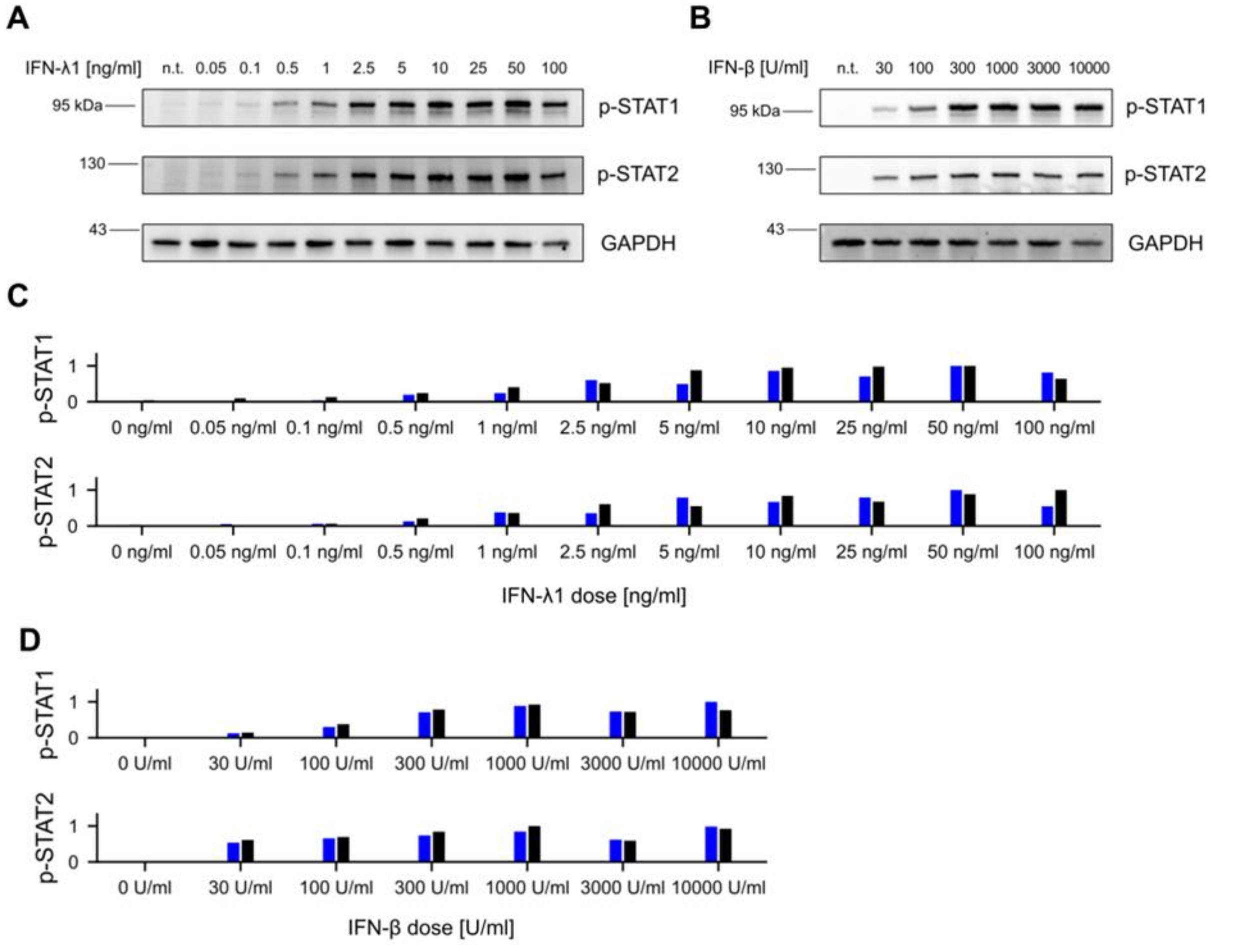


**Figure S2. IFN-β and IFN-λ dose–response analysis, determination of saturating concentrations.**

A Cells were stimulated with an array of concentrations of IFN-λ1 for 30 min before lysis.

B Cells were stimulated with an array of concentrations of IFN-β for 30 min before lysis.

C Quantification of 2 experimental replicates, blot shown in panel A is marked in blue.

D Quantification of 2 experimental replicates, blot shown in panel B is marked in blue.

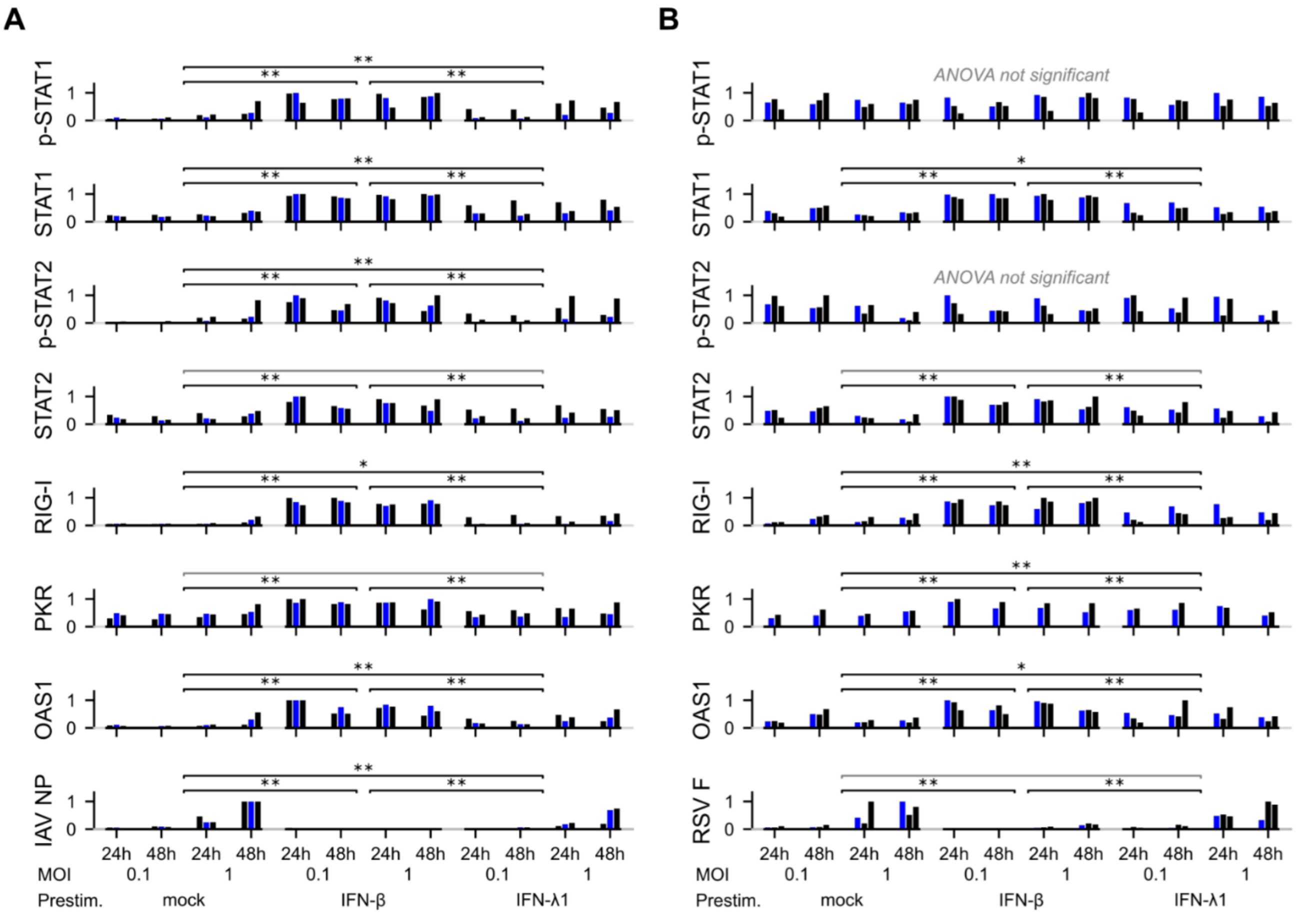


**Figure S3.**

A Quantification of three replicates of the experiment shown in Fig. 2A. Blue bars indicate the experimental replicate shown in Fig. 2A.

B Quantification of three replicates of the experiment shown in Fig. 2B. Blue bars indicate the experimental replicate shown in panel Fig. 2B.

In panels A and B, quantifications of the bands without infection with IAV or RSV (n.t.) are omitted. Each blot quantification is normalized to the maximum. Differences between prestimulations were determined using multiple ANOVA tests on log intensities, corrected with 5% Benjamini-Hochberg false discovery rate. For proteins with significant differences, Tukey post-hoc tests were performed to obtain p values: $p < 0.05$ (*) and $p < 0.01$ (**) (see Methods for details).

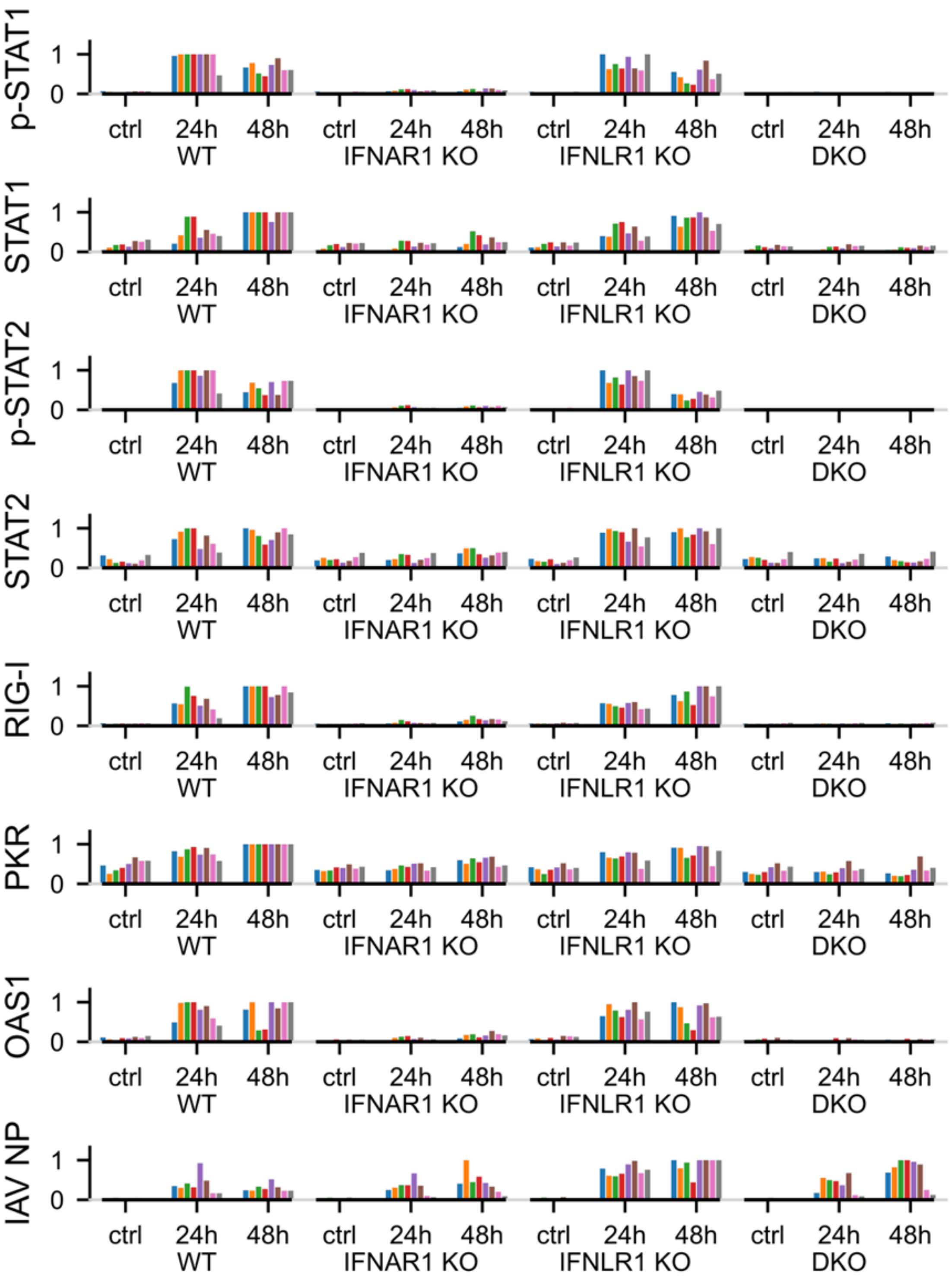


**Figure S4. Effect of IFNLR1 and IFNAR1 knockout on IAV proliferation and activation of the innate immune response** – full quantification of experiments shown in Fig. 3A. Colors correspond to independent experimental replicates of experiments shown in Fig. 3A

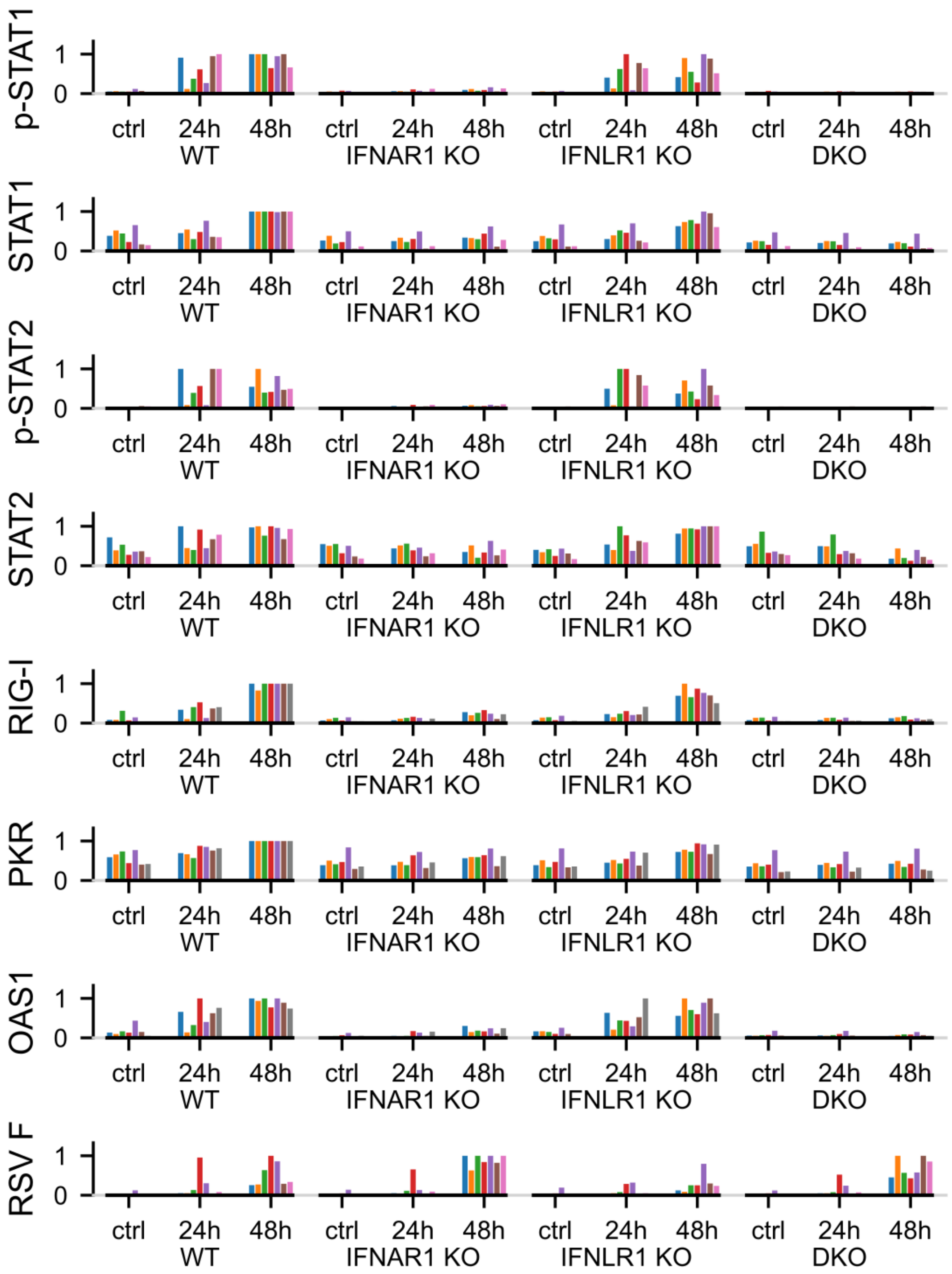


**Figure S5. Effect of IFNLR1 and IFNAR1 knockout on RSV proliferation and activation of the innate immune response** – full quantification of experiments shown in Fig. 3B. Colors correspond to independent experimental replicates of experiments shown in Fig. 3B.

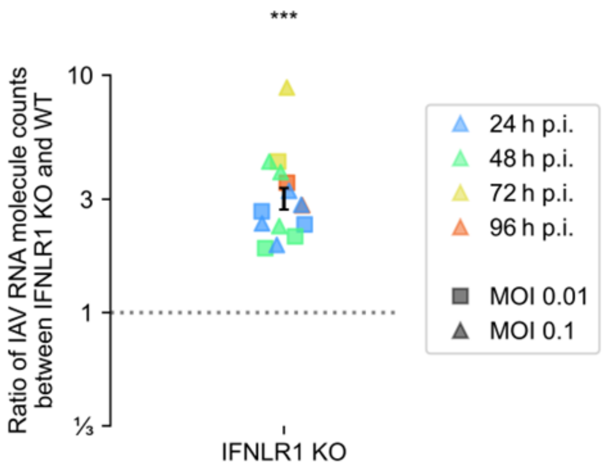


**Figure S6. Effect of IFNLR1 deficiency on IAV RNA expression**.

Levels of IAV RNA were measured by dPCR for A549 WT and IFNLR1 KO cells at 24, 48, 72 and 96 h p.i. at MOIs of 0.01 and 0.1 (as indicated by the legend). For each experimental replicate the ratio between IAV RNA expression in IFNLR1 KO and WT cells was calculated, and each square/triangle corresponds to an experimental replicate. Triple asterisk (***) indicates that IAV RNA expression is significantly higher in IFNLR1 KO cells than in WT cells ($p < 0.001$, one sample two-tailed Student t-test on log ratios between IFNLR1 KO and WT cells).

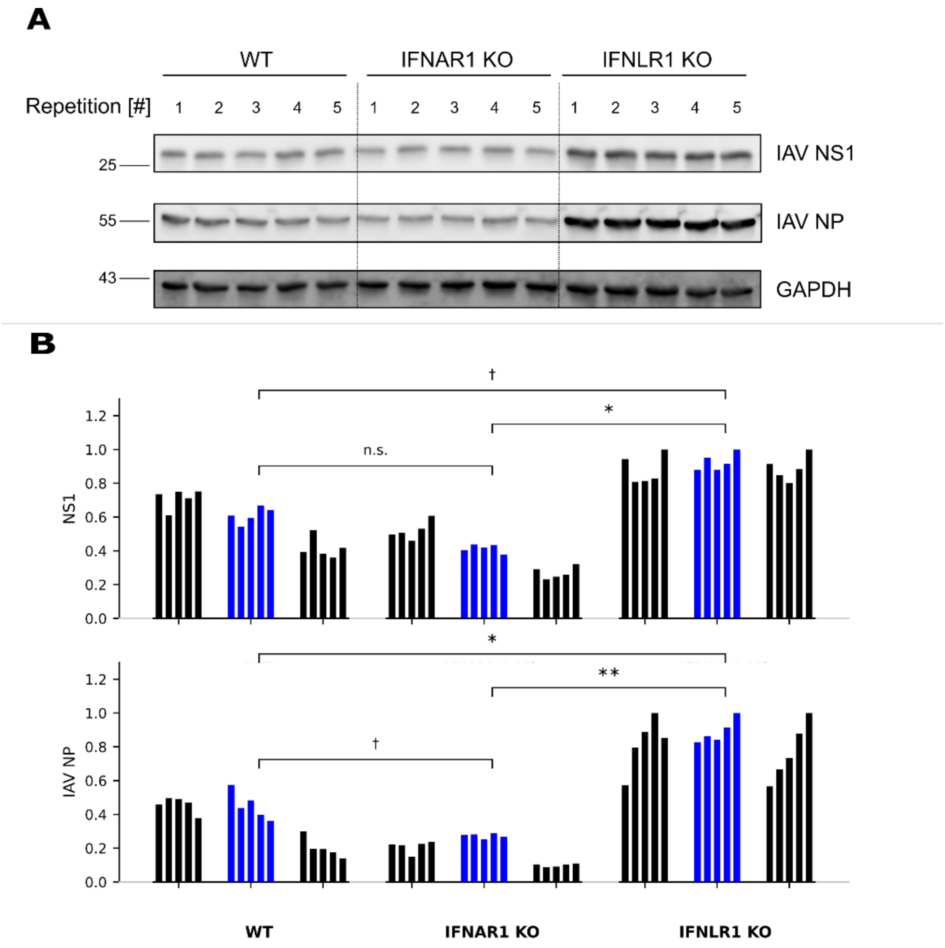


**Figure S7. Expression of IAV proteins, NS1 and NP, in WT, IFNAR1 KO, and IFNLR1 KO cells.**

A. A549 WT, IFNAR1 KO, IFNLR1 KO cells were infected with IAV at an MOI of 0.1 and lysed 48 h p.i. Each lane corresponds to a single technical replicate (separate well). Western blots of the two remaining biological replicates are provided in Source Data 1.
B. Quantifications of three biological replicates (each consisting of five technical replicates). Each blot quantification is normalized to the maximum. Quantification of the shown blot is marked in blue. Differences between cell lines were determined using multiple ANOVA tests on log intensities, corrected with 5% Benjamini-Hochberg false discovery rate. Tukey post-hoc tests were performed to obtain p-values: † $p < 0.1$, * $p < 0.05$, ** $p < 0.01$ (see Methods for details).

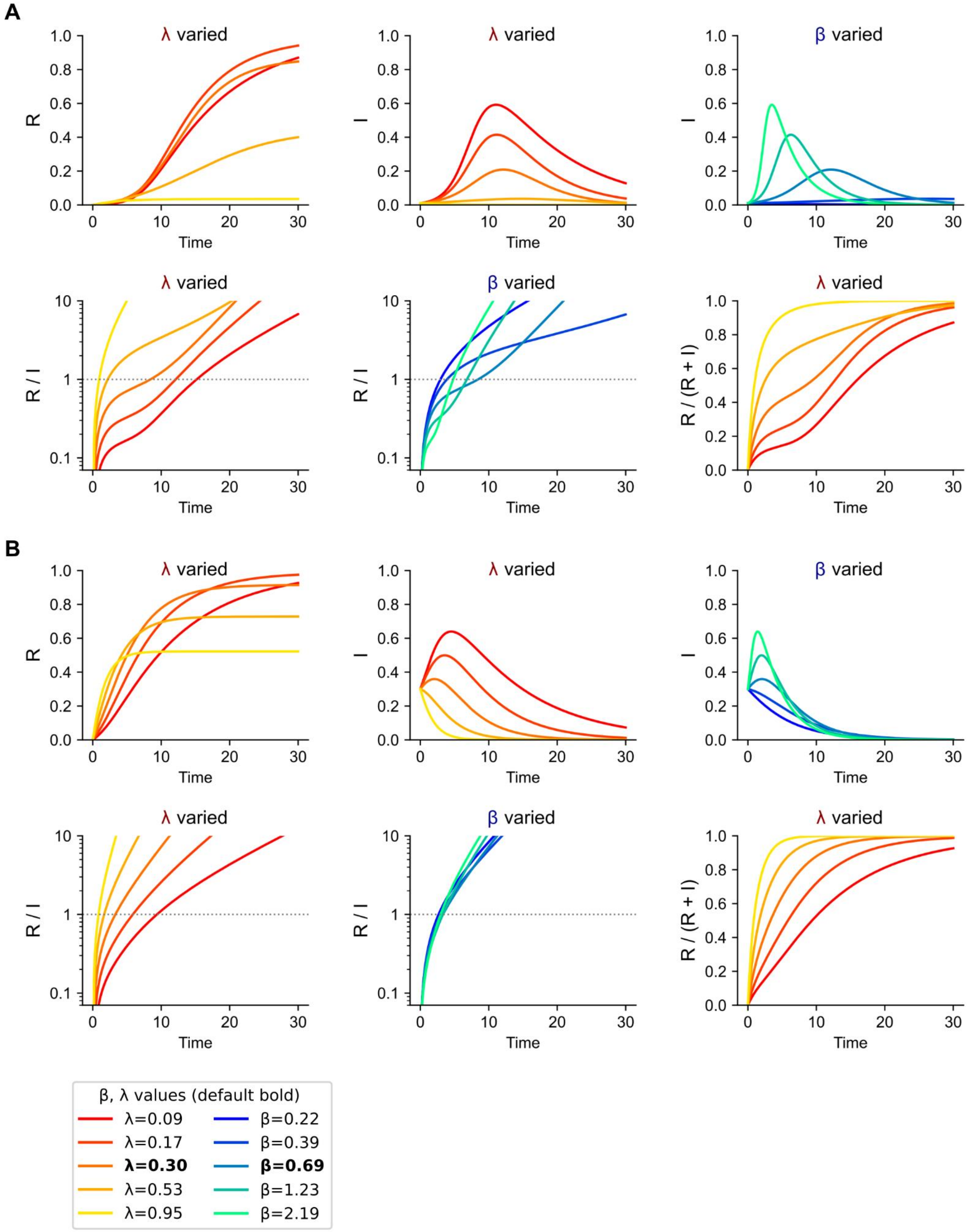


**Figure S8. Kinetics of infections in the SIR model.** The figure supplements Fig. 5 with data for an MOI of 0.01 (A) and 0.3 (B).

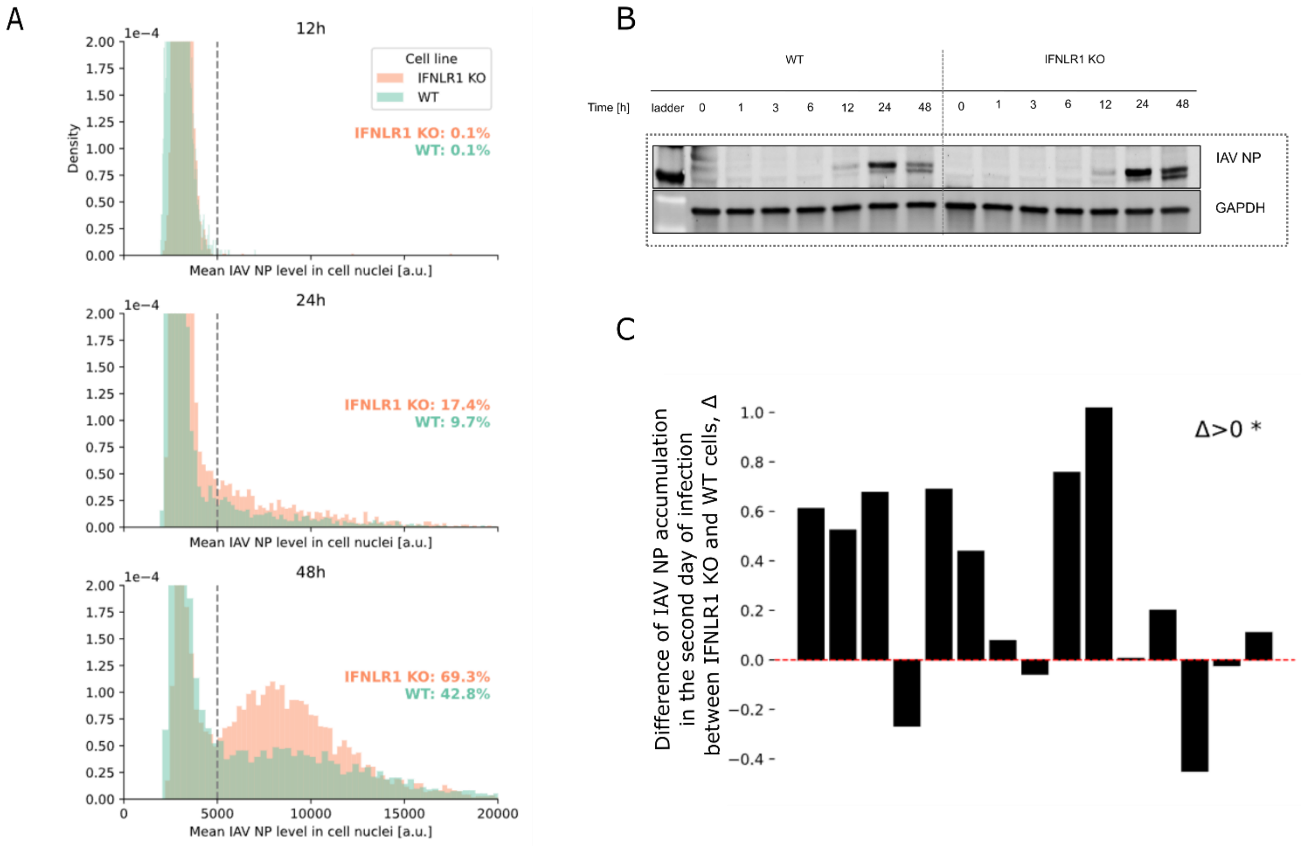


**Figure S9. Kinetics of infection with IAV in WT and IFNLR1 KO cells.**

A. Cells were infected with IAV at an MOI of 0.01, then fixed (at times indicated) and immunostained with an antibody for IAV NP to determine the fraction of virus-positive cell. Normalized histograms show fraction of cells as a function of the mean IAV NP signal in the nucleus. The analysis of additional two replicates is provided in Source Data S1.
B. Cells were infected with IAV at an MOI of 0.01 and lysed at indicated times p.i.. Western blots for three additional replicates are provided in Source Data S1.
C. Statistical analysis of accumulation of IAV nucleoprotein (between 24 and 48 h p.i.) in WT and IFNLR1 KO cells, infected with IAV at an MOI of 0.1. All 14 Western blots, in which levels of IAV NP was determined for WT and IFNLR1 KO cells, at 24 and 48 h p.i., are used for analysis. Height of the bar is equal to $\Delta=\ln(KO_{48}/KO_{24})-\ln(WT_{48}/WT_{24})$, were $KO_{48}$, $KO_{24}$, $WT_{48}$, and $WT_{24}$ denote normalized IAV NP levels in IFNLR1 KO and WT at 48 and 24 h. Positive Δ value indicates faster accumulation of IAV NP in IFNLR1 KO than in WT population. One-sampled, two-tailed Student t-test indicated that $\Delta>0$, * $p<0.05$.

#### 2 Supplementary Methods

In this section, statistical methods for the analysis of the significance of results presented in the main and supplementary figures are described.

In Fig. S1B, the factors in ANOVA were: stimulation agent (IFN-β or IFN-λ), stimulation time (1, 2, 6, 24, 48 h for p-STAT1/2; 24, 48 h for STAT1/2), replicate ID (assuming no interaction with other factors). For each considered protein, the significance of the stimulation agent was tested, and the results were corrected using the Benjamini–Hochberg procedure (with a 5% false discovery rate). In Fig. 1, the factors in ANOVA were concentration of poly(I:C), presence or absence of IFN-β costimulation, and cell line (WT or IFNLR1 KO). The significance of the cell line was tested for a non-zero concentration of poly(I:C), separately for the presence or absence of IFN-β costimulation. In Figs. 2B and 2D, the factors in ANOVA were: stimulation agent (mock, IFN-β, IFN-λ), stimulation conditions (MOI 0.1 24 h, MOI 0.1 48 h, MOI 1 24 h, MOI 1 48 h), replicate ID (assuming no interaction with other factors). For each considered protein, the significance of the stimulation agent was tested, and the results were corrected using the Benjamini–Hochberg procedure (5% false discovery rate). For proteins in which a significant difference was observed, pairwise significance levels between stimulation agents obtained by the Tukey post-hoc test are reported. In Figs. 3B and 3D, the factors in ANOVA were: cell line (WT, IFNAR1 KO, IFNLR1 KO), stimulation time (24 h, 48 h), replicate ID (assuming no interaction with other factors); with the exception of RSV F in Fig. 3D, where only measurements at 48 h were considered and thus only two factors remain. For each considered protein, the significance of the cell line was tested, and the results were corrected using the Benjamini–Hochberg procedure (with a 5% false discovery rate). In all cases a significant difference was observed, and pairwise significance levels between cell lines obtained by the Tukey post-hoc are reported (for each pair of cell lines).

In Fig. S1C, RT-PCR was similarly analyzed using ANOVA. Data for each mRNA were analyzed independently, with following factors: stimulation agent (IFN-β or IFN-λ), stimulation time (6, 10, 24 h), replicate ID (assuming no interaction with other factors). For each mRNA the significance of the stimulation agent was tested, and the results were corrected using the Benjamini-Hochberg procedure (with 5% false discovery rate).

Statistical significance in Figs. 4A, 4B, and Fig. S5 was assessed using the Student t-test. In Fig. 6A we performed both a Welch t-test (unpaired Student t-test without assuming equal variances) and Mann–Whitney U test, while in Fig. 6B we performed a Student t-test and Wilcoxon signed-rank U test. All tests assume two-tailed alternatives.

In the case of Fig. 4A, Fig. 6A, Fig. S5, the statistical tests were performed using experimental replicates obtained for two or three time points pooled together; similarly, in Fig. 4A and Fig. S5 two MOI values (0.01 and 0.1) are considered jointly. In these cases our data indicates that the studied relative differences between cell lines do not change over time or with MOI (for the considered time periods and MOIs).

In particular, in Fig. 6A, the left column consists of IAV NP level ratios between IFNLR1 KO and WT cells, where both levels are measured in a single Western blot. The right column consists of ratios between the fraction of dying/dead cells between IFNLR1 KO and WT cells; both fractions are measured in a single FACS experiment. However, ratios from the left column are not paired with ratios from the right column (since the Western blot and FACS measurements were not paired).

Thus, we use a (parametric) Welch's t-test and a (non-parametric) Mann–Whitney U test to compare log ratios from the left column with log ratios from the right column. Data from time points 24 h p.i. and 48 h p.i. are considered jointly.

In Fig. 6B, ratios from the left column (obtained from immunostaining images of fixed cells) and ratios from the right column (obtained from live-cell imaging) are paired, since both types of measurement were performed for the same cell population (for each experimental replicate). For each left–right pair, we can thus compute a ratio-of-ratios. We then perform a (parametric) two-tailed Student t-test and a (non-parametric) two-tailed Wilcoxon signed-rank test on log ratio-of-ratios. Data from time points 24 h p.i. and 48 h p.i. and cell lines WT and IFNLR1 KO are considered jointly, since the death rate in IAV is always higher than in RSV (which is equivalent to ratio-of-ratios greater than 1).

#### 3 Supplementary Data

Source Data S1 (PDF File) containing:

Images of all Western blot replicates,

Figures with quantifications of replicates of the experiments shown in Fig. 1, Fig. S9,

Supplementary Tables with quantifications of the experiments shown in Fig. 6 and Fig. S9C.

Source Data S2 (Zipped folder). Data quantifications in CSV format (Western blots, dPCR, RT-PCR, FACS, cell counts from immunostaining) and Python codes to generate figures.
