## Supplementary Data S1 for "Type III interferons may suppress viral infections by triggering cell death"

**Figure 1** / rep2 (Influence of IFN-λ signaling on poly(I:C) induced cell death)

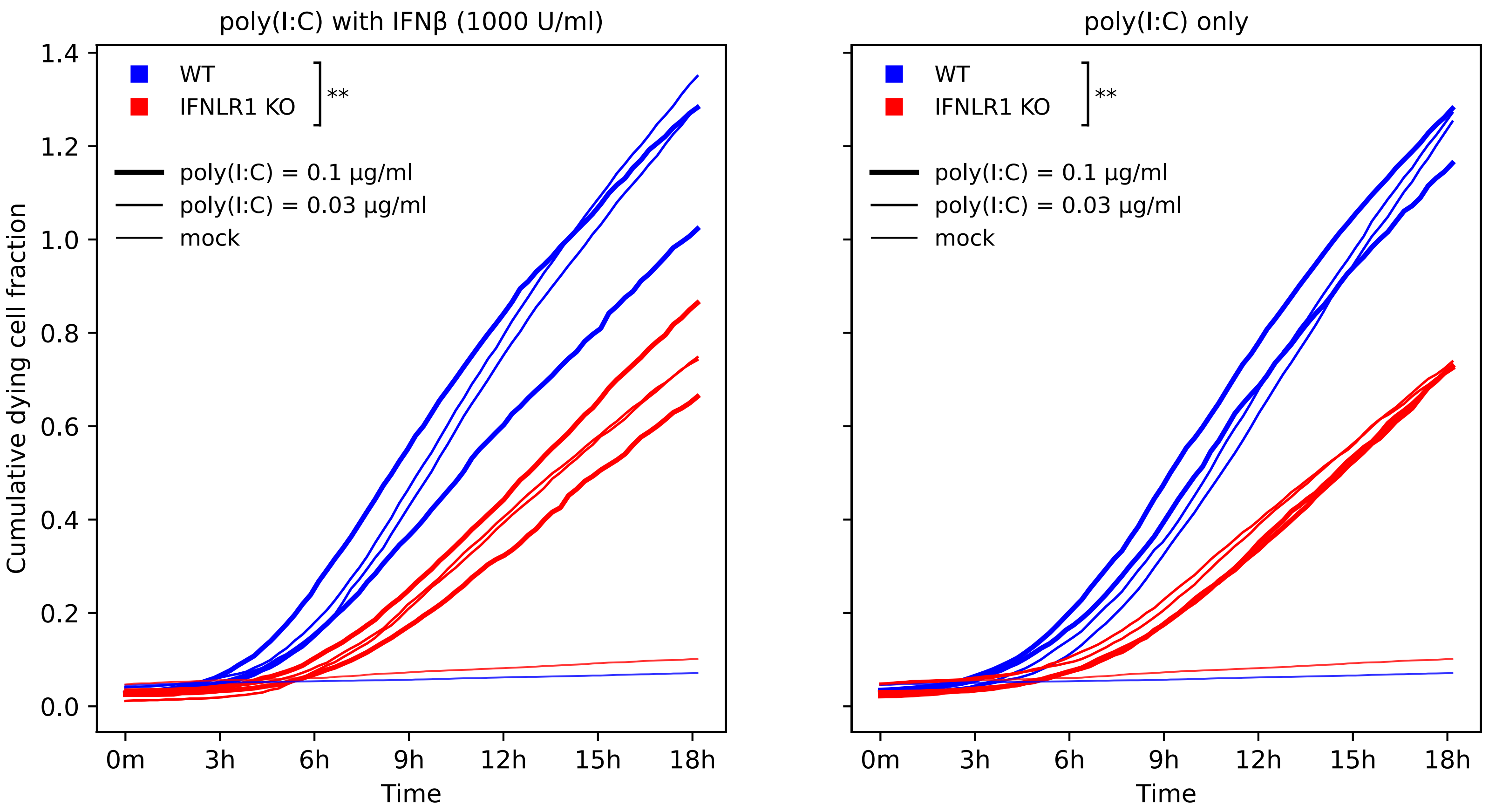

**Figure 1** / rep3 (Influence of IFN-λ signaling on poly(I:C) induced cell death)

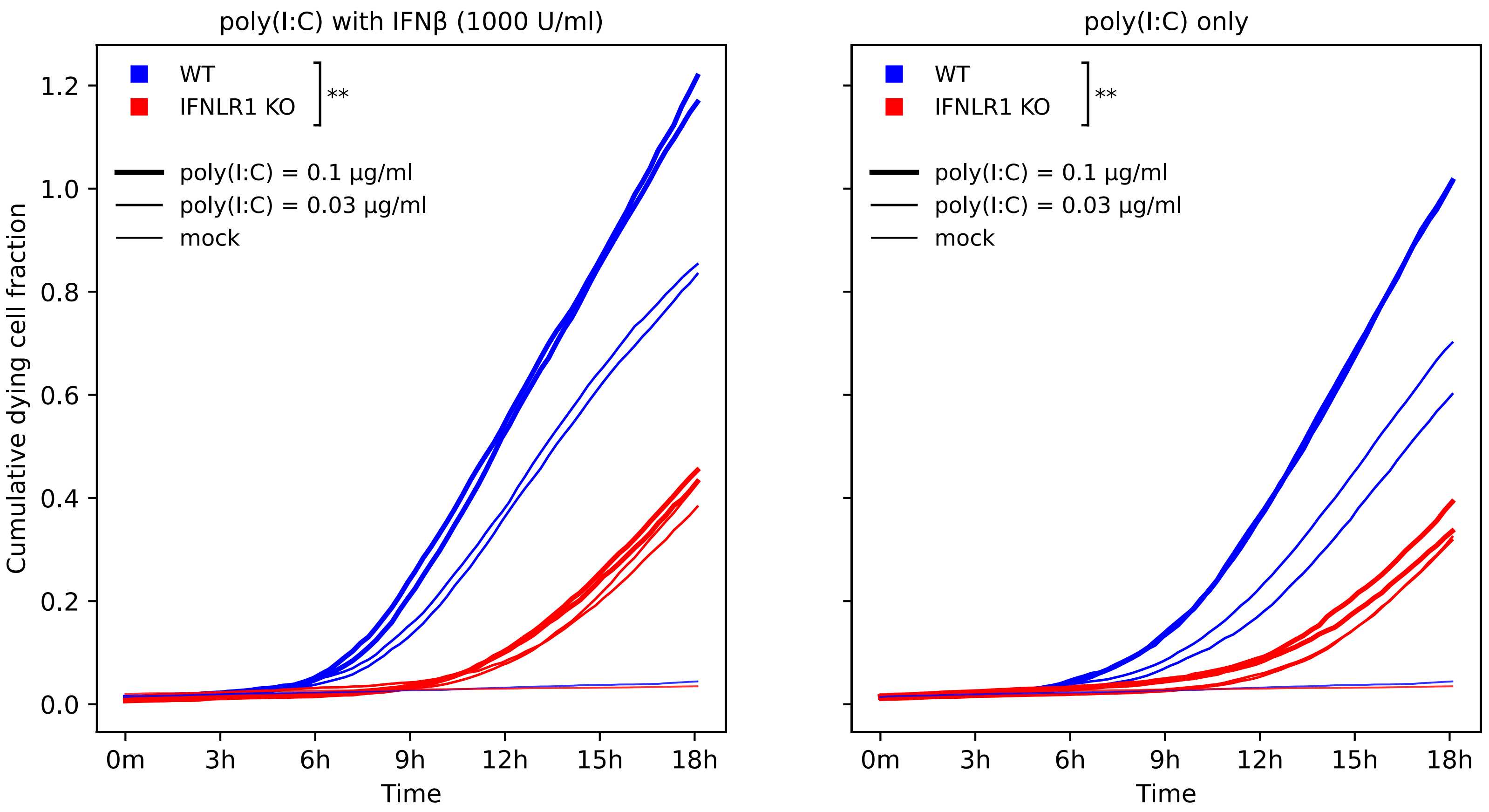

### **Figure S3A** / rep1 (**IAV** after IFN-β and IFN-λ1)

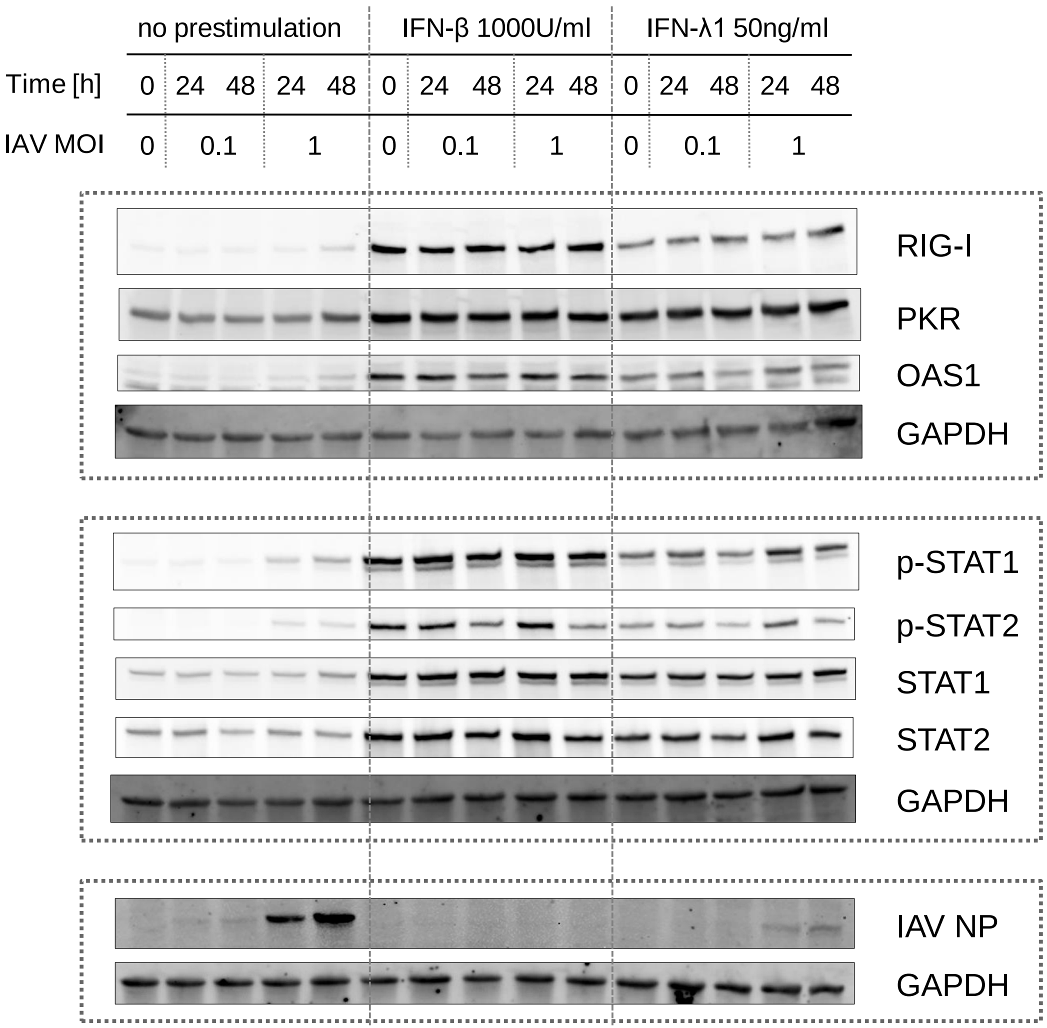


Replicate 1, quantified for Fig. S3A

### **Figure S3A** / rep2 (**IAV** after IFN-β and IFN-λ1)

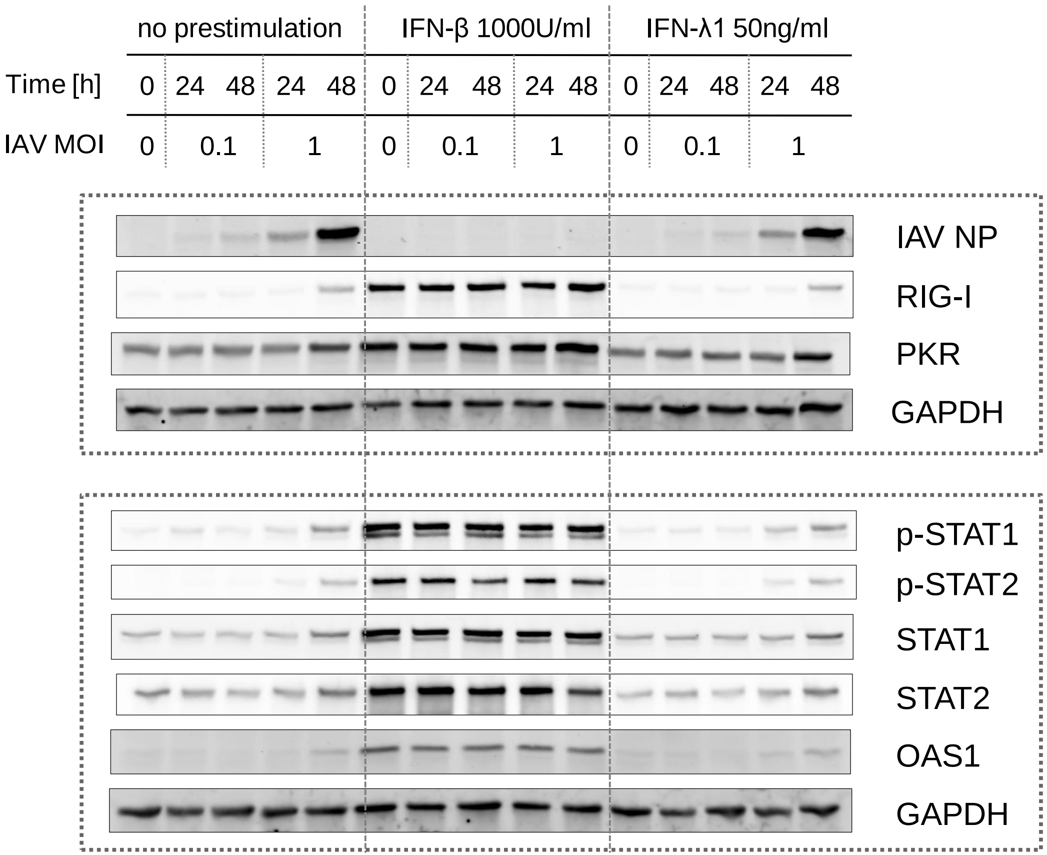


Replicate 2,
quantified for Fig. S3A

### **Figure S3A** / rep3 (**IAV** after IFN-β and IFN-λ1)

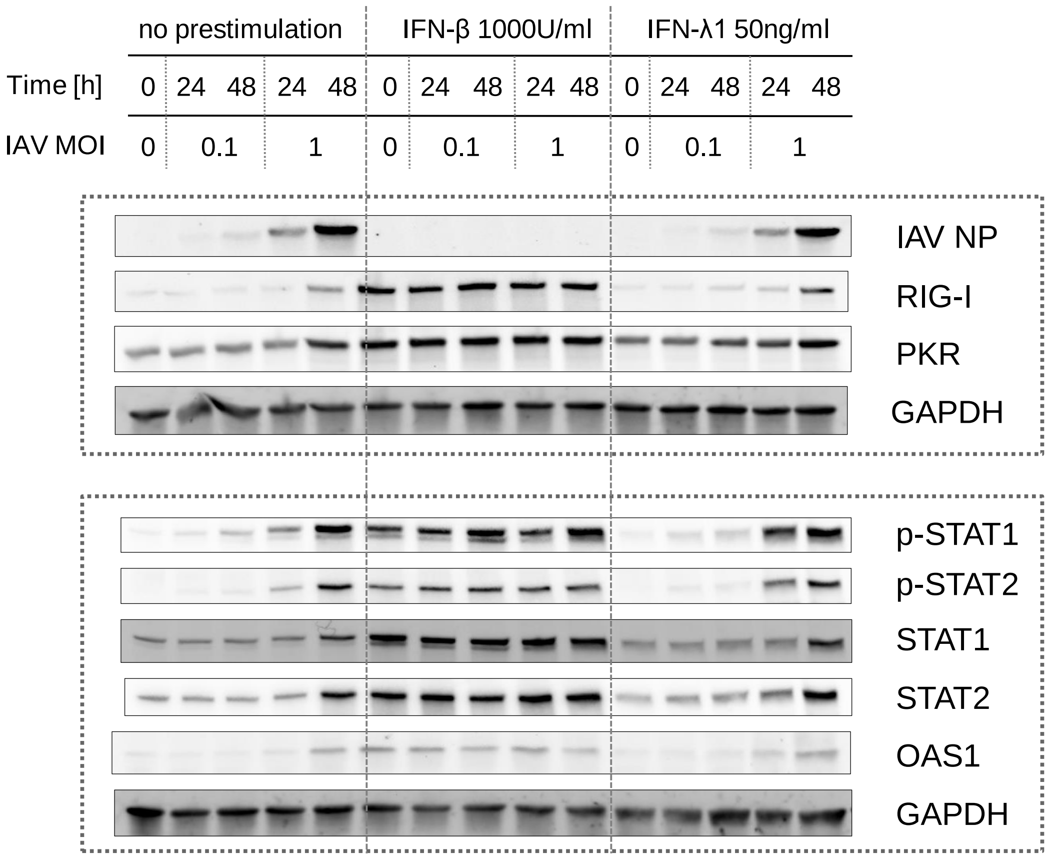


Replicate 3, quantified for Fig. S3A

### **Figure S3B** / rep1 (**RSV** after IFN-β and IFN-λ1)

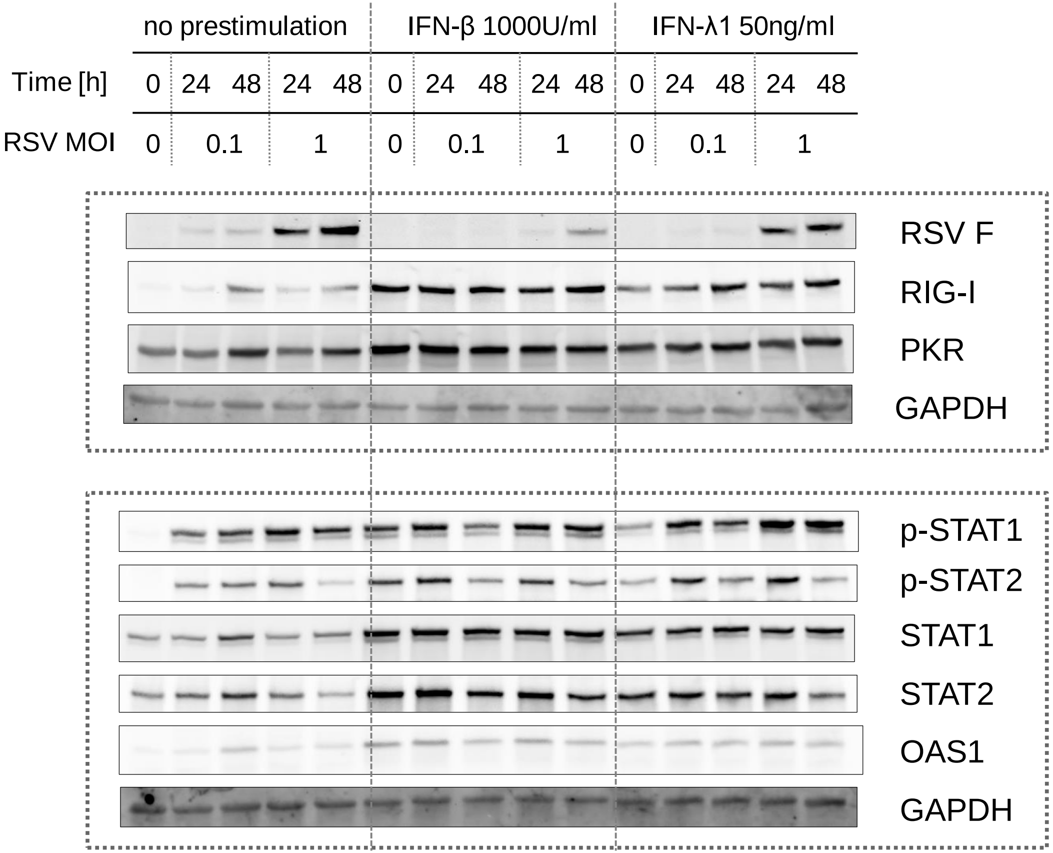


Replicate 1, quantified for Fig. S3B

### **Figure S3B** / rep2 (**RSV** after IFN-β and IFN-λ1)

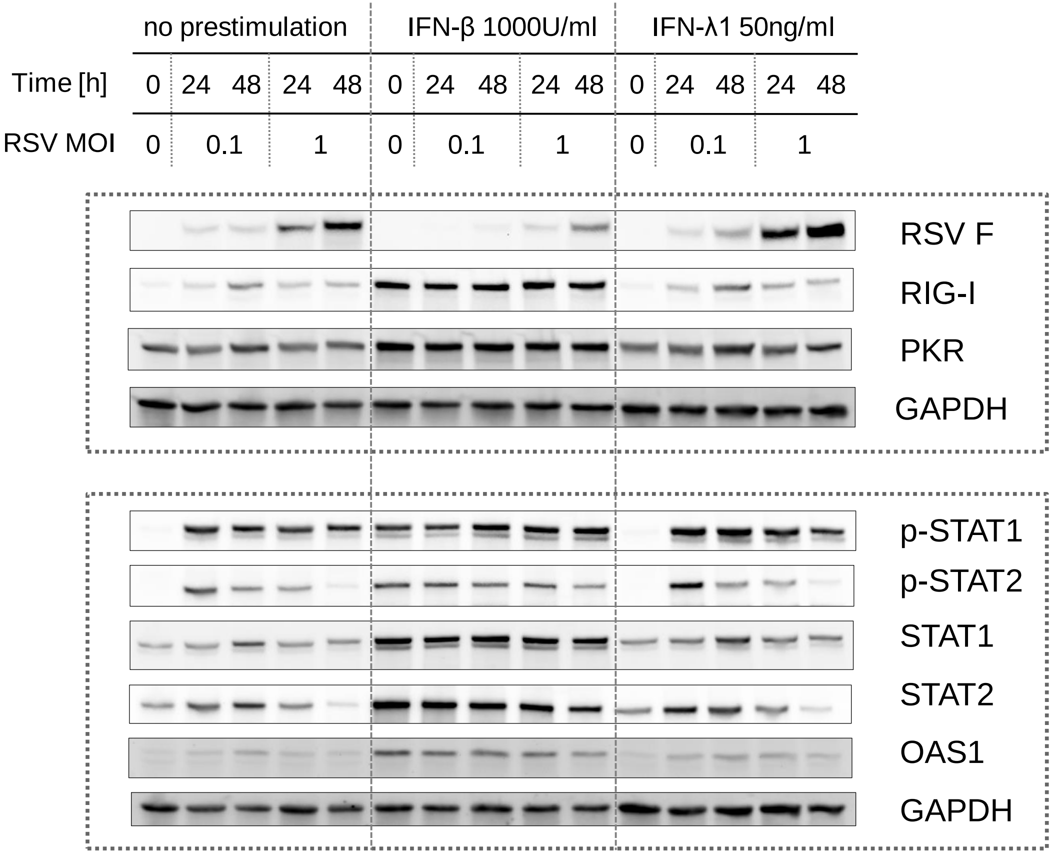


Replicate 2, quantified for Fig. S3B

**Figure S3B** / rep3 (**RSV** after IFN-β and IFN-λ1)

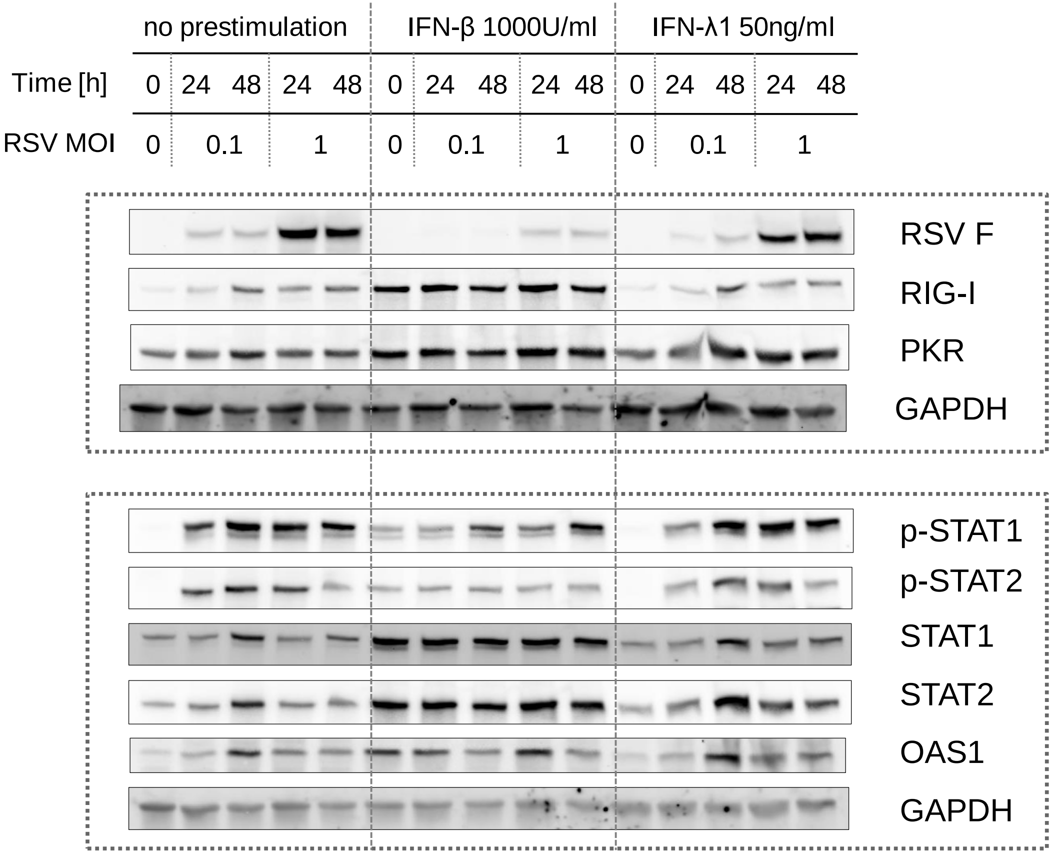


Replicate 3, quantified for Fig. S3B

### **Figure 3B** / rep1 (**IAV** in IFN_R KOs)

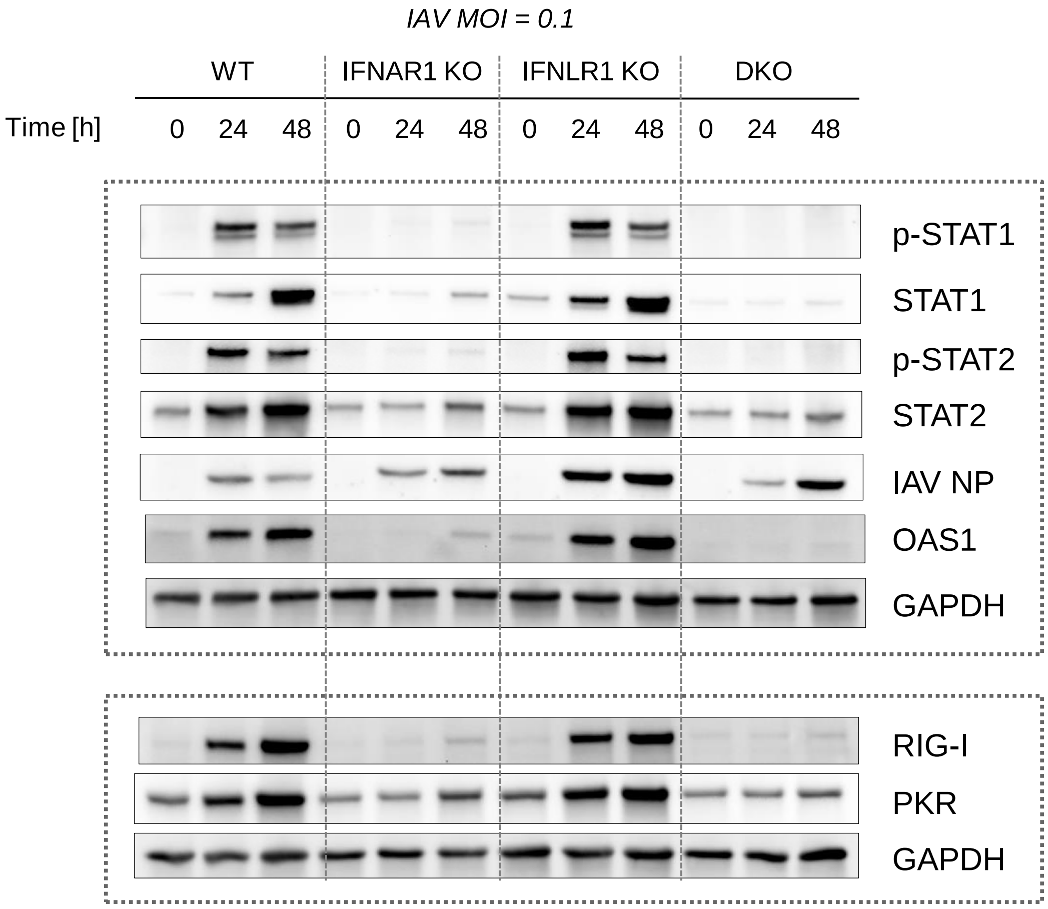


Replicate 1,
quantified in Fig. 3B

**Figure 3B** / rep2 (**IAV** in IFN_R Kos)

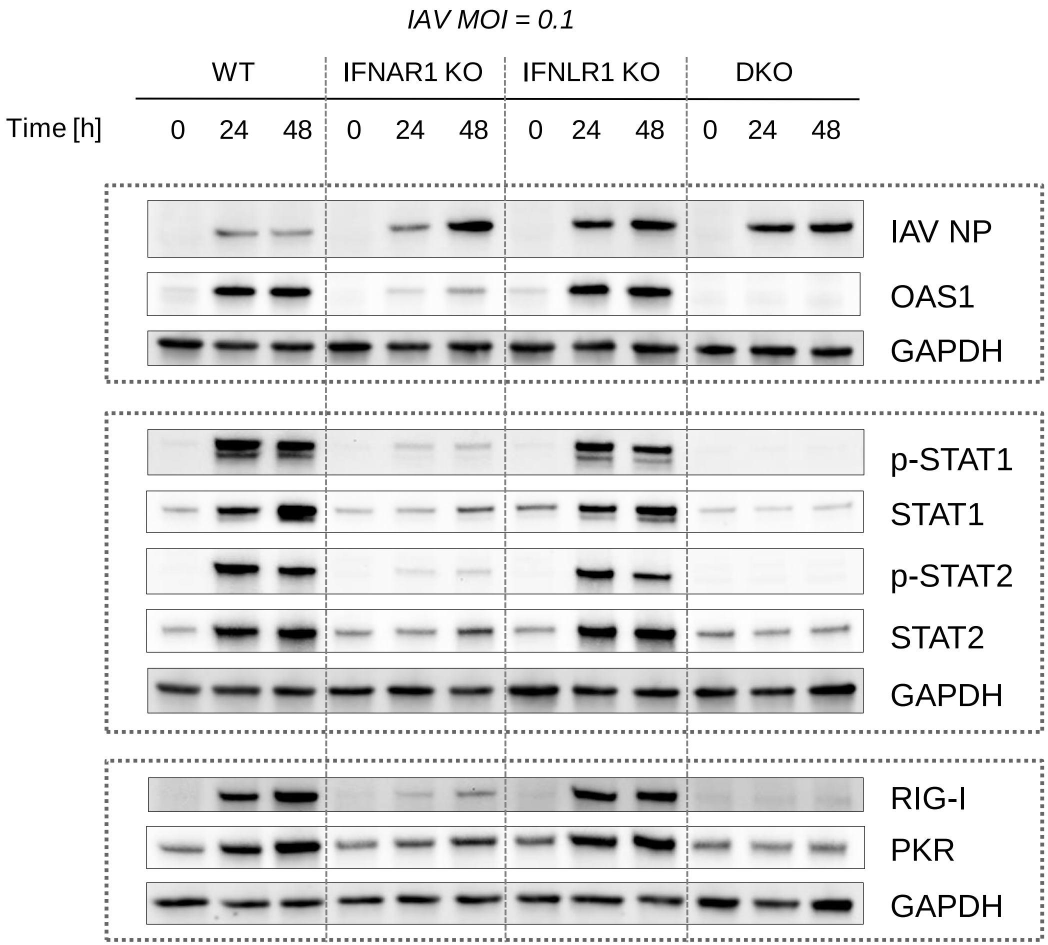


Replicate 2, quantified for Fig. 3B

### **Figure 3B** / rep3 (**IAV** in IFN_R KOs)

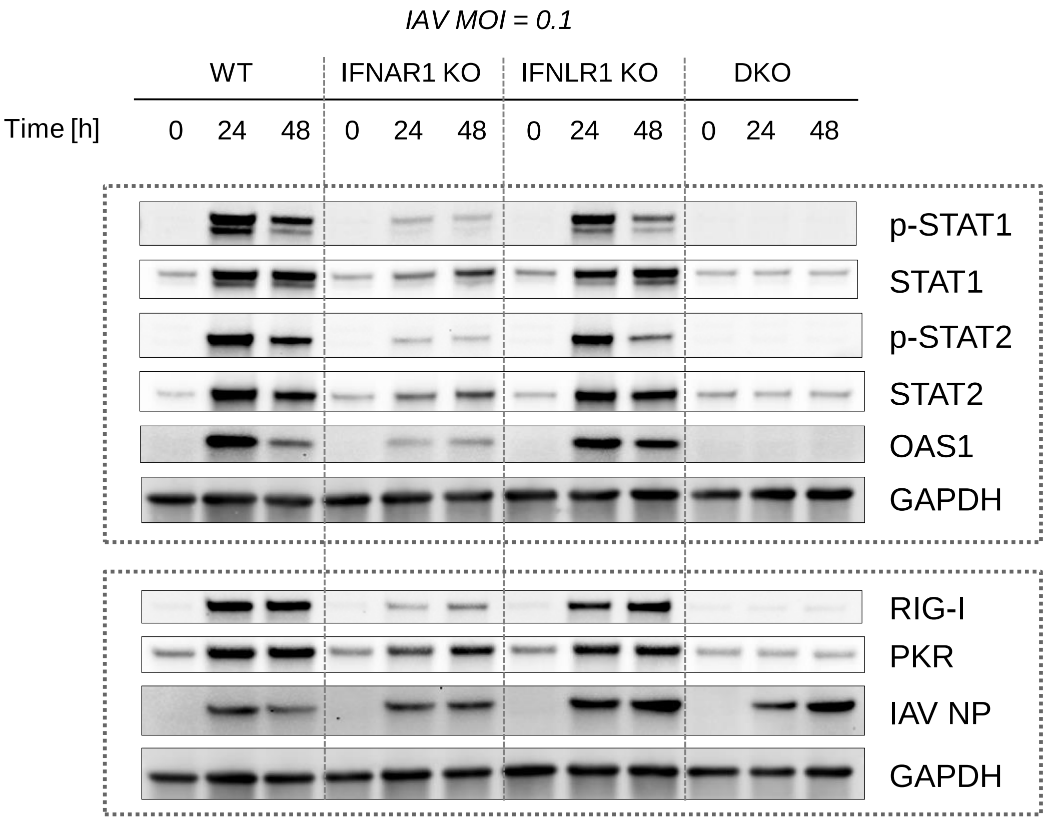


Replicate 3, quantified for Fig. 3B

### **Figure 3B** / rep4 (**IAV** in IFN_R KOs)

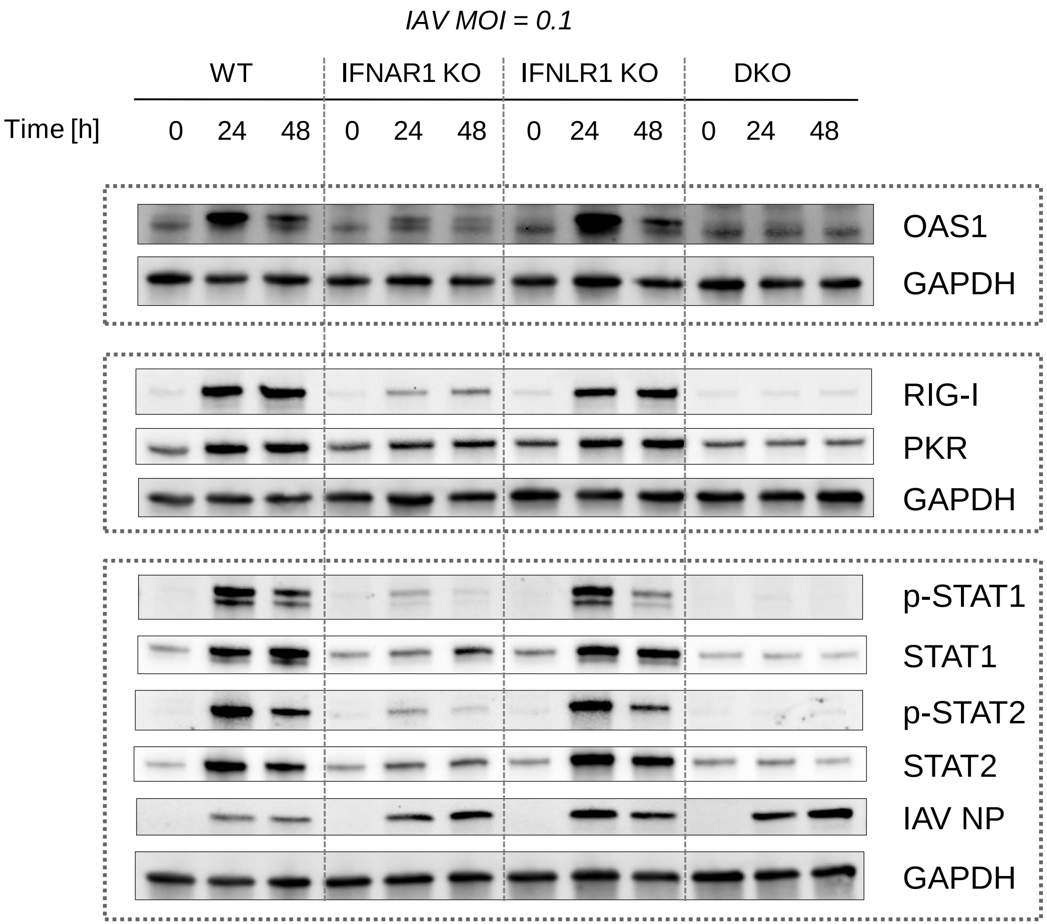


Replicate 4, quantified for Fig. 3B

**Figure 3B** / rep5 (**IAV** in IFN_R KOs)

Replicate 5, quantified for Fig. 3B

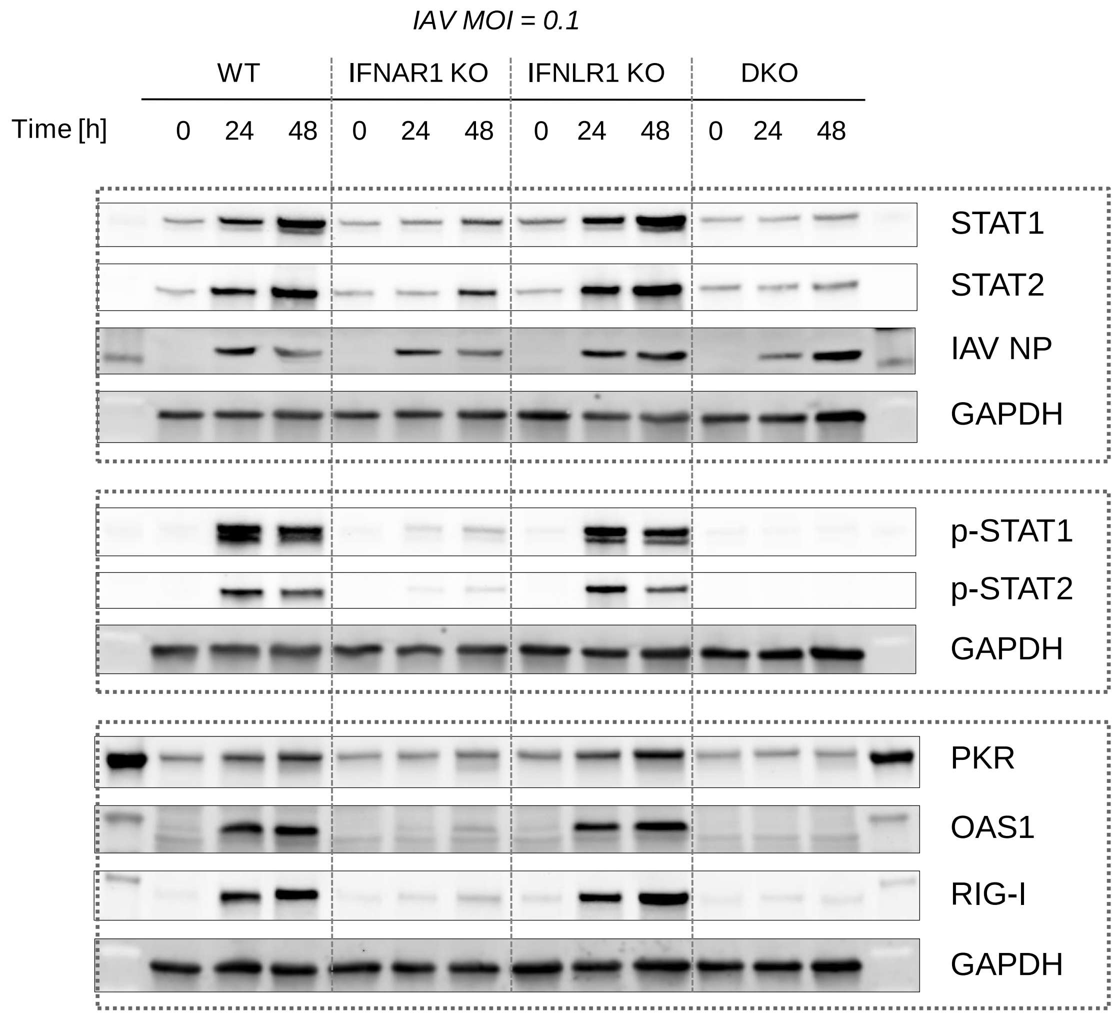

**Figure 3B** / rep6 (**IAV** in IFN_R KOs)

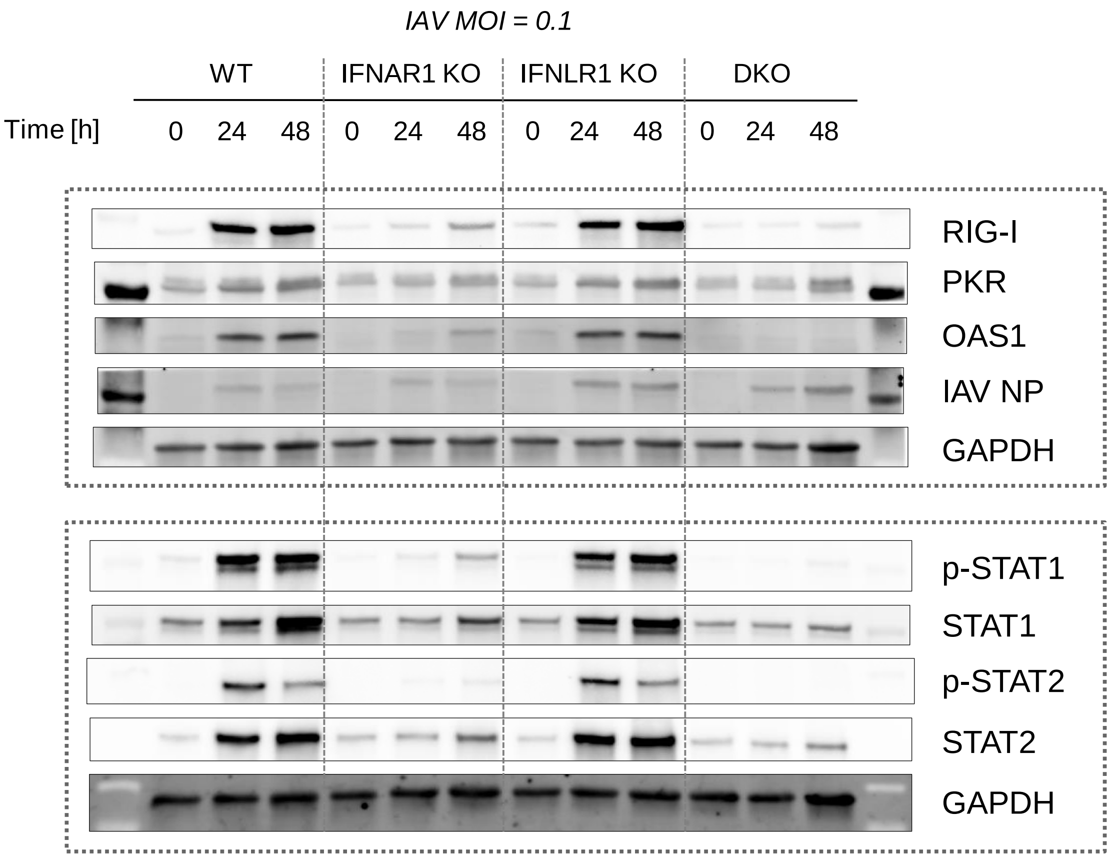


Replicate 6, quantified for Fig. 3B

**Figure 3B** / rep7 (**IAV** in IFN_R KOs)

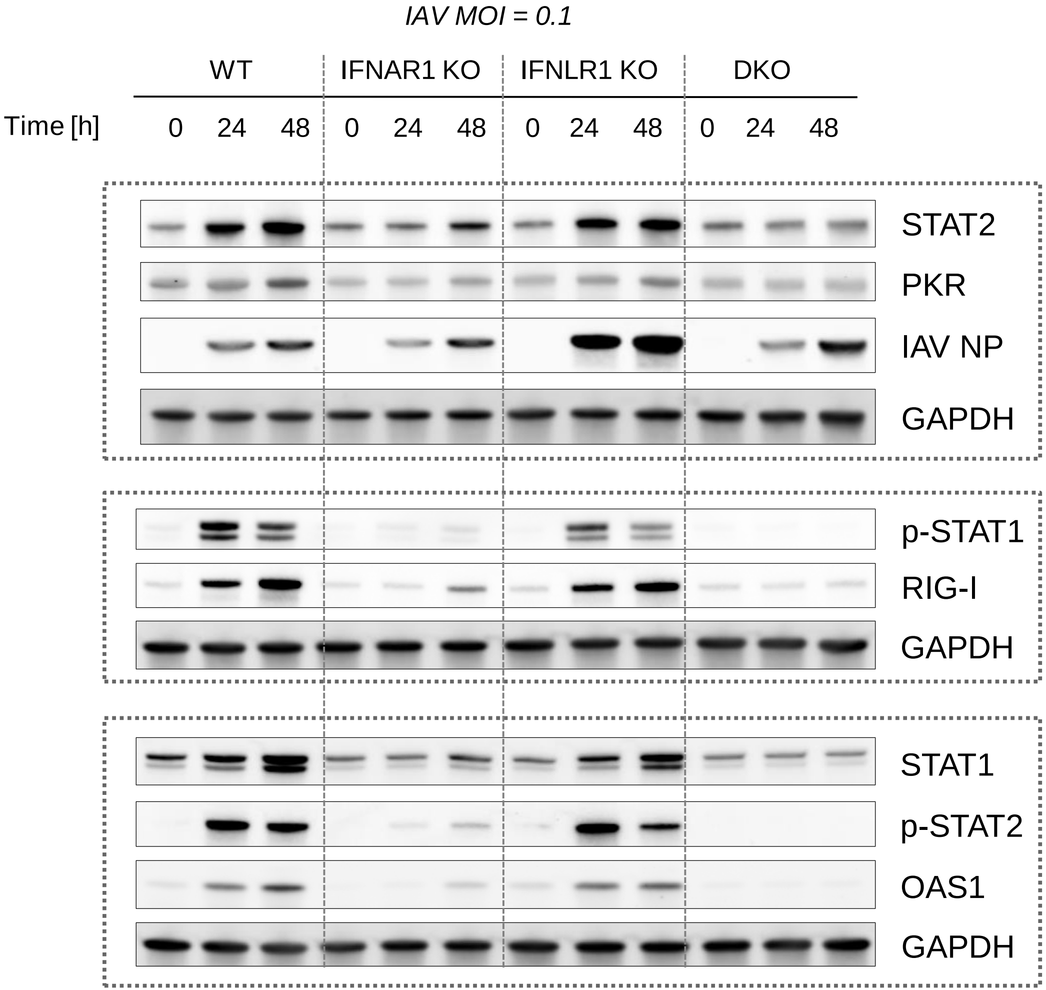


Replicate 7, quantified for Fig. 3B

**Figure 3B** / rep8 (**IAV** in IFN_R KOs)

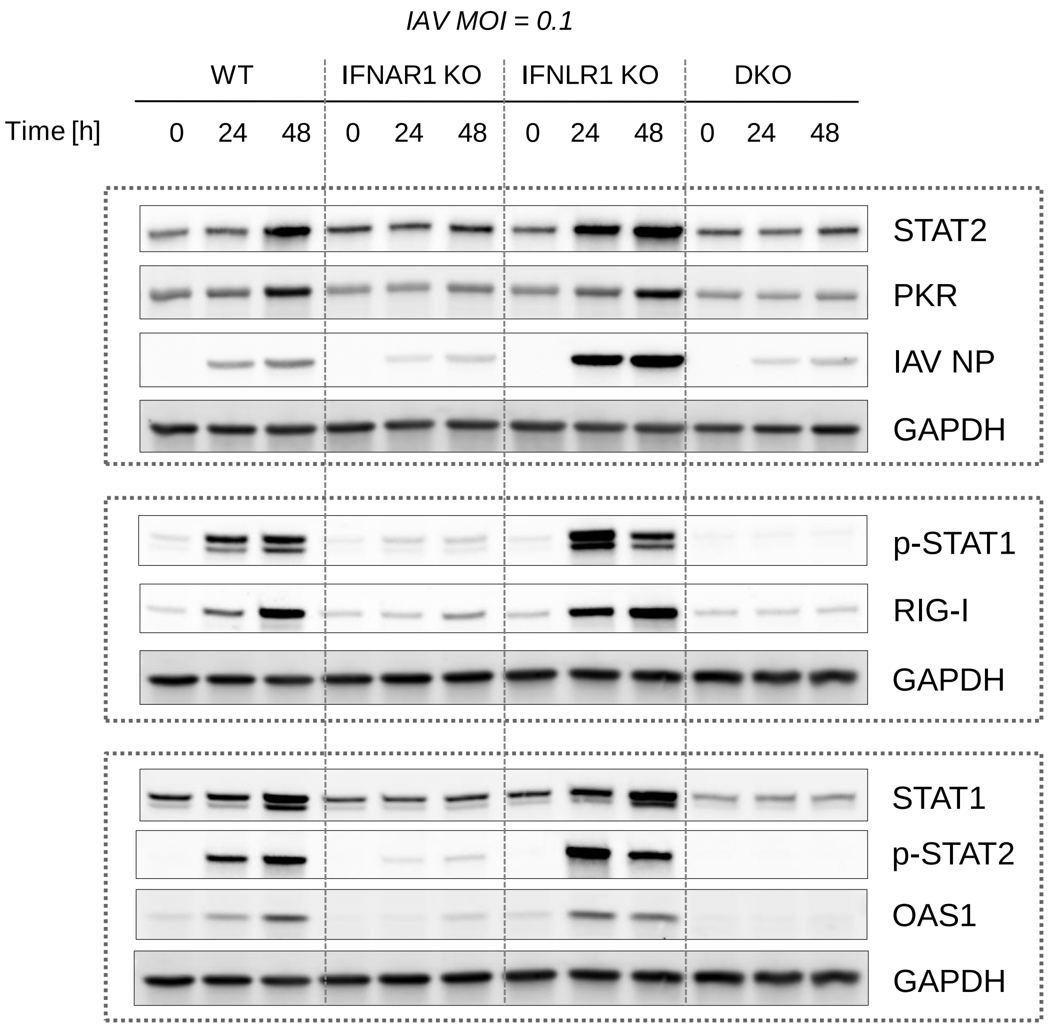


Replicate 8, quantified for Fig. 3B

### **Figure 3D** / rep1 (**RSV** in IFN_R KOs)

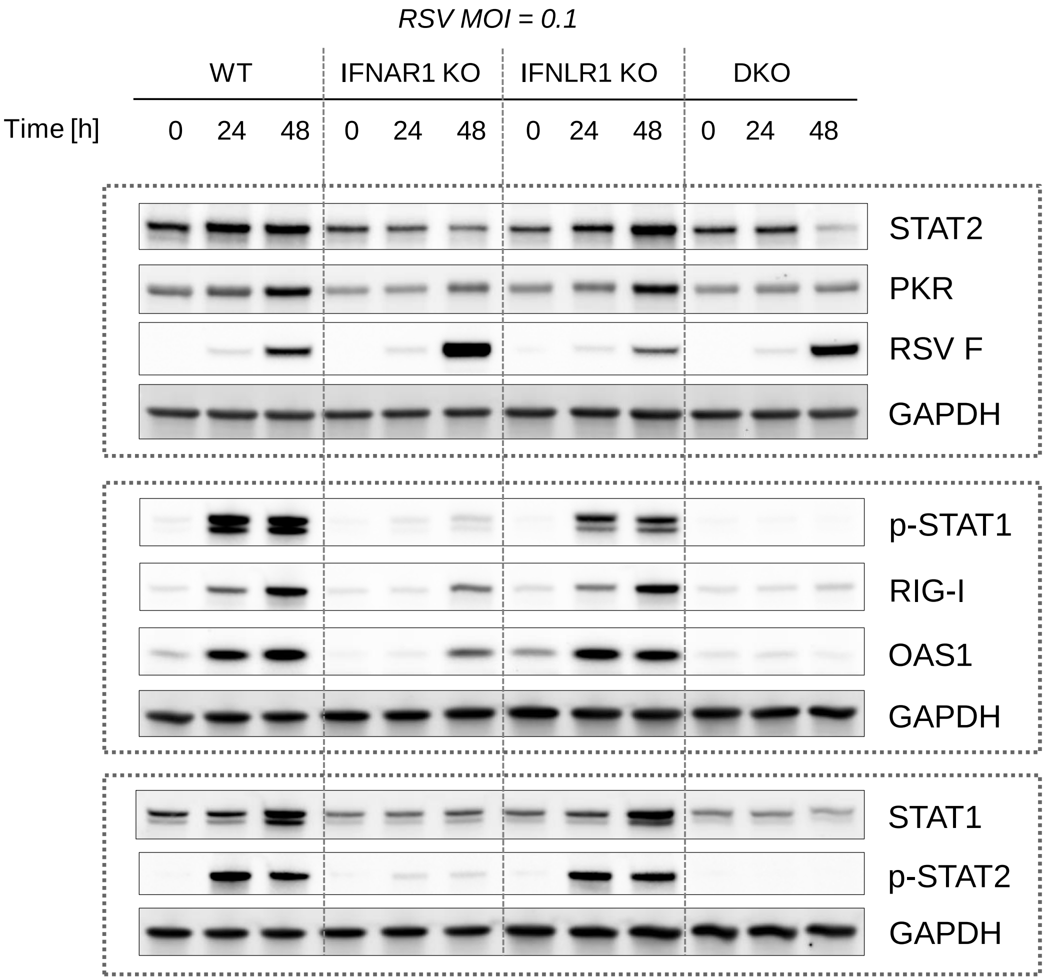


Replicate 1, quantified for Fig. 3D

### **Figure 3D** / rep2 (**RSV** in IFN_R KOs)

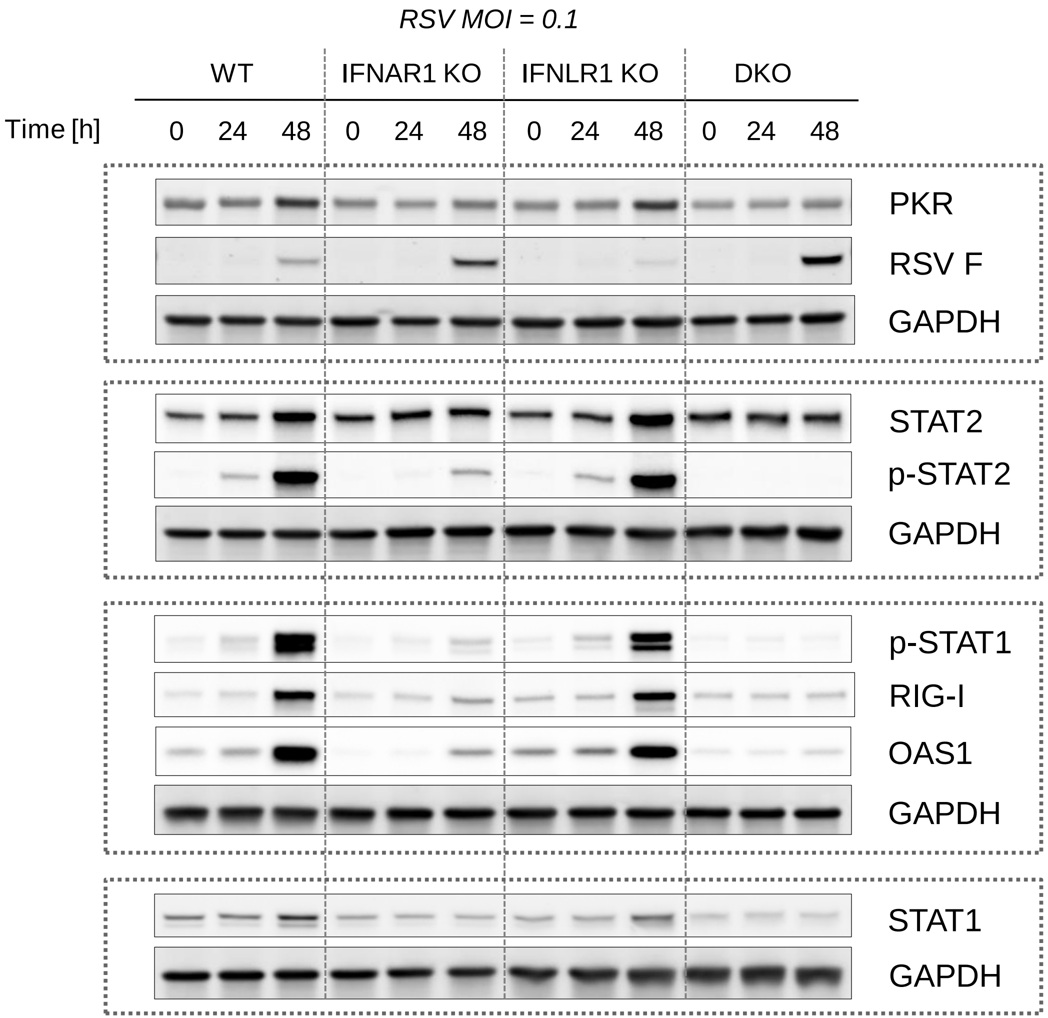


Replicate 2,
quantified for Fig. 3D

### **Figure 3D** / rep3 (**RSV** in IFN_R KOs)

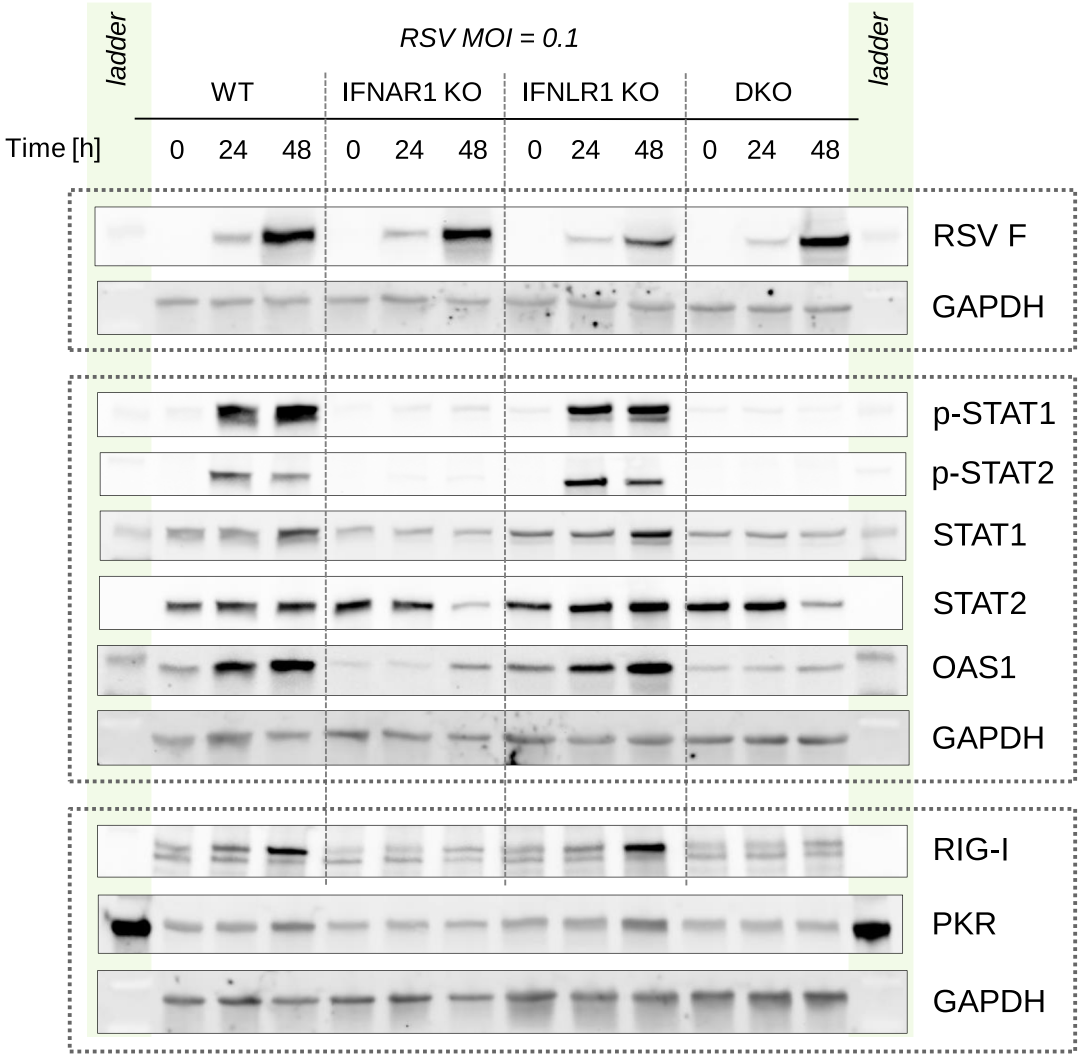


Replicate 3, quantified for Fig. 3D

### **Figure 3D** / rep4 (**RSV** in IFN_R KOs)

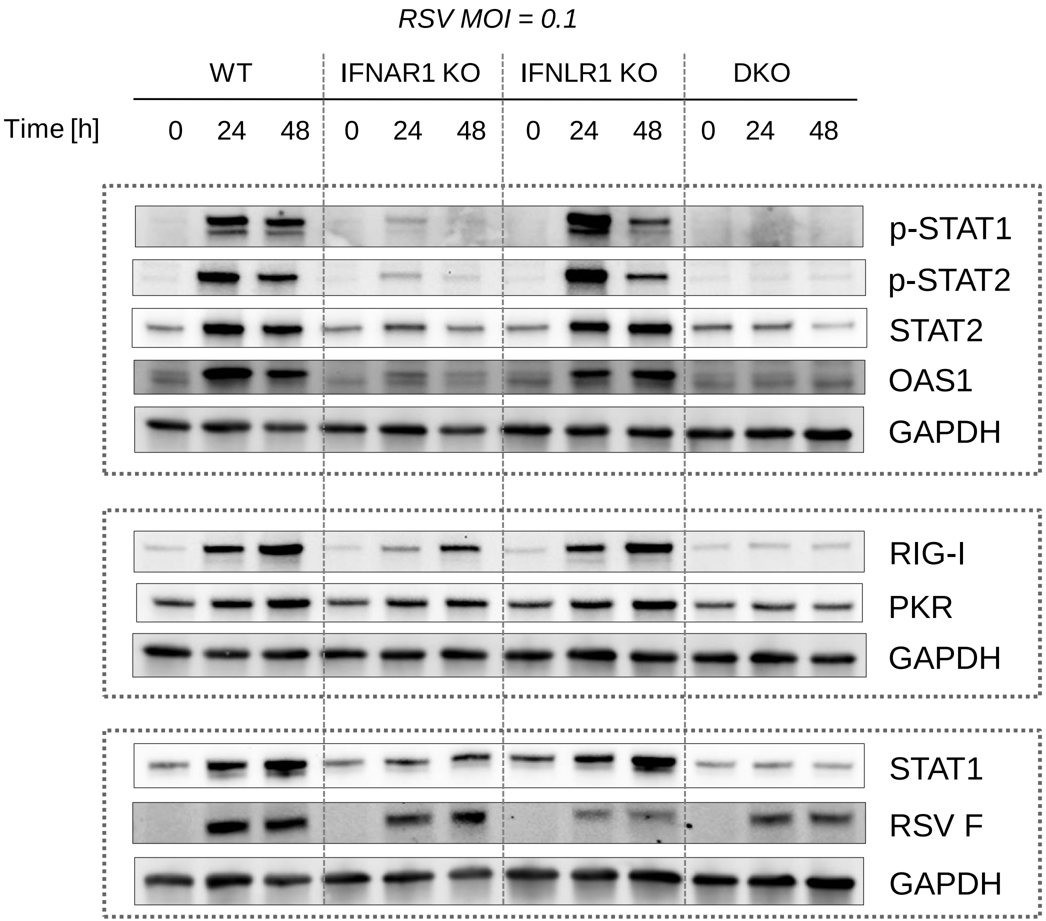


Replicate 4, quantified for Fig. 3D

### **Figure 3D** / rep5 (**RSV** in IFN_R KOs)

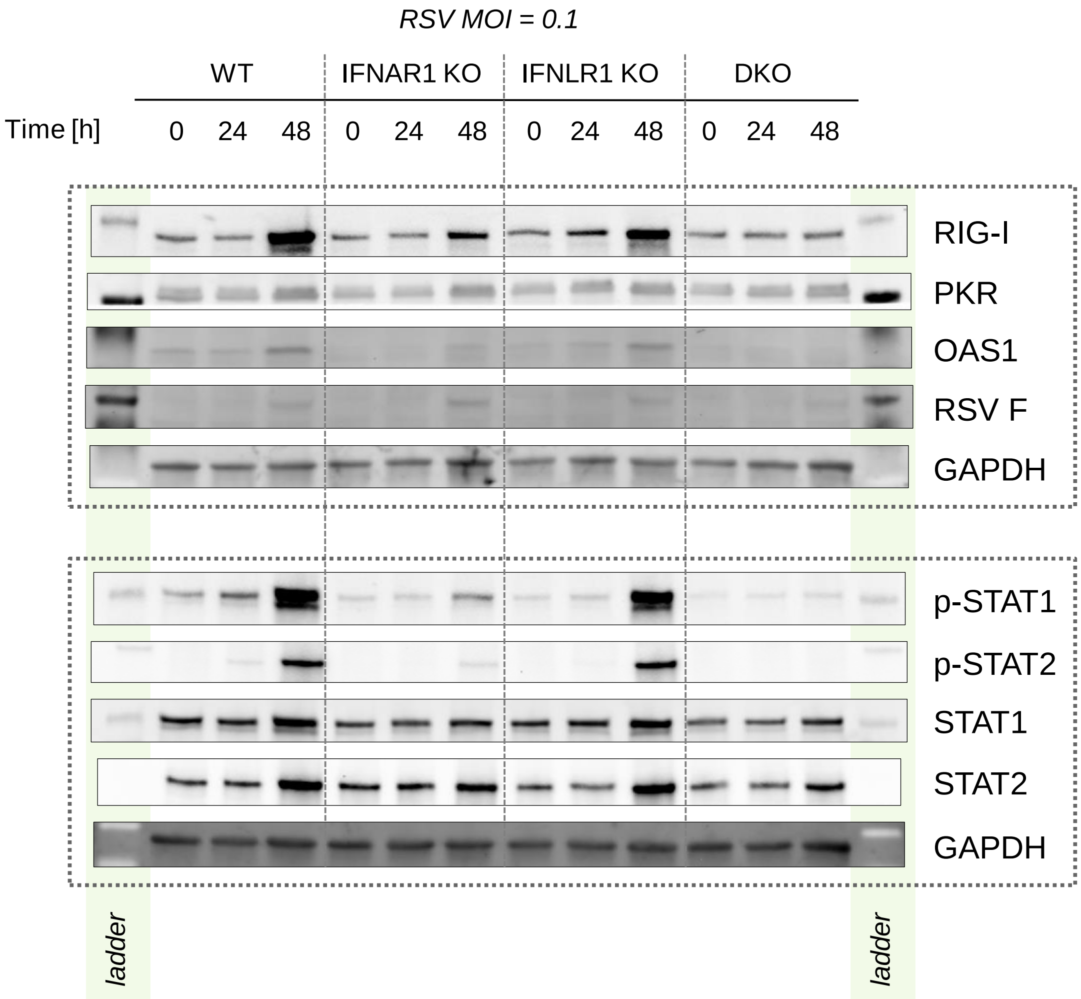


Replicate 5,
quantified for Fig. 3D

**Figure 3D** / rep6 (**RSV** in IFN_R KOs)

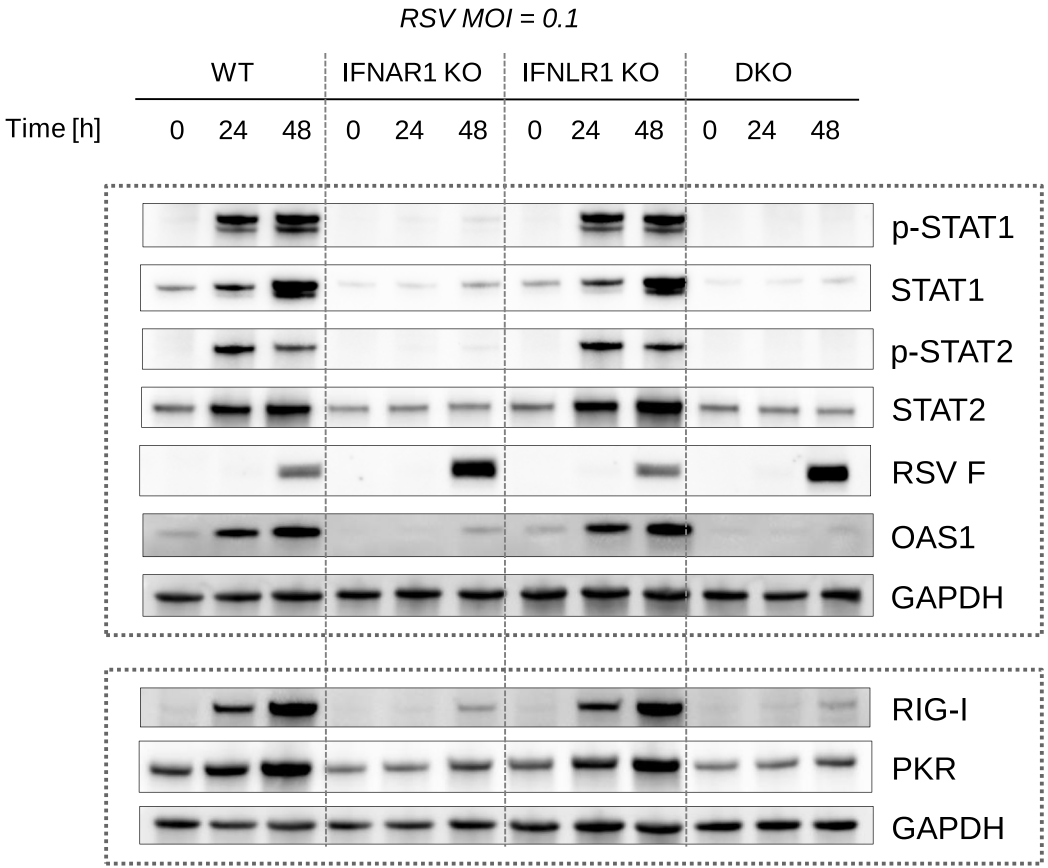


Replicate 6, quantified for Fig. 3D

### **Figure 3D** / rep7 (**RSV** in IFN_R KOs)

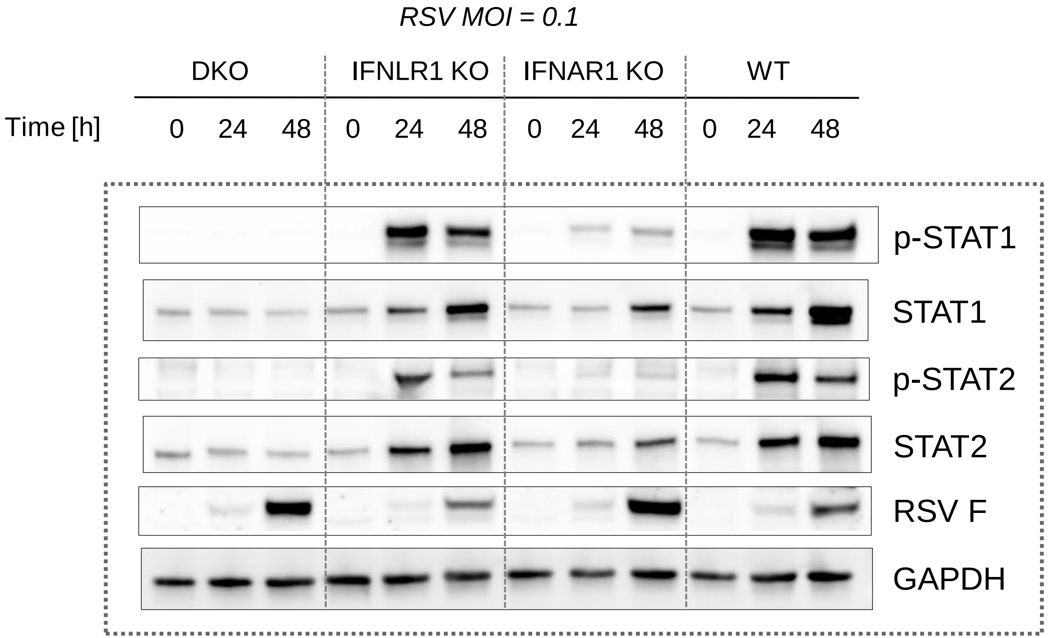


Replicate 7, quantified for Fig. 3D

### **Figure 3D** / rep7 continued (**RSV** in IFN_R KOs)

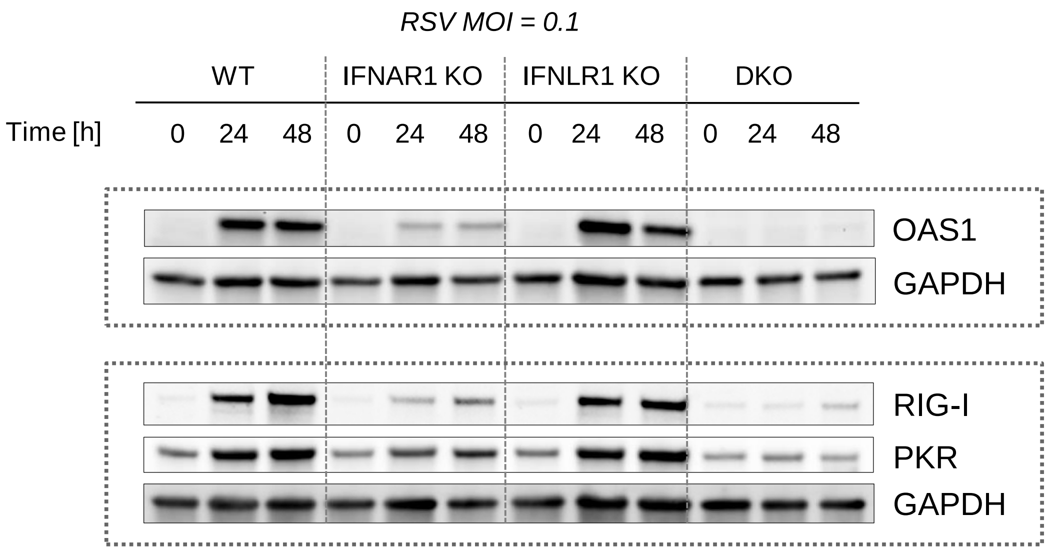


Replicate 7 continued, quantified for Fig. 3D

### Corresponding to Figures 4C and 4D

Immunostaining images from
Figures 4C and 4D without pSTAT1 channel

C

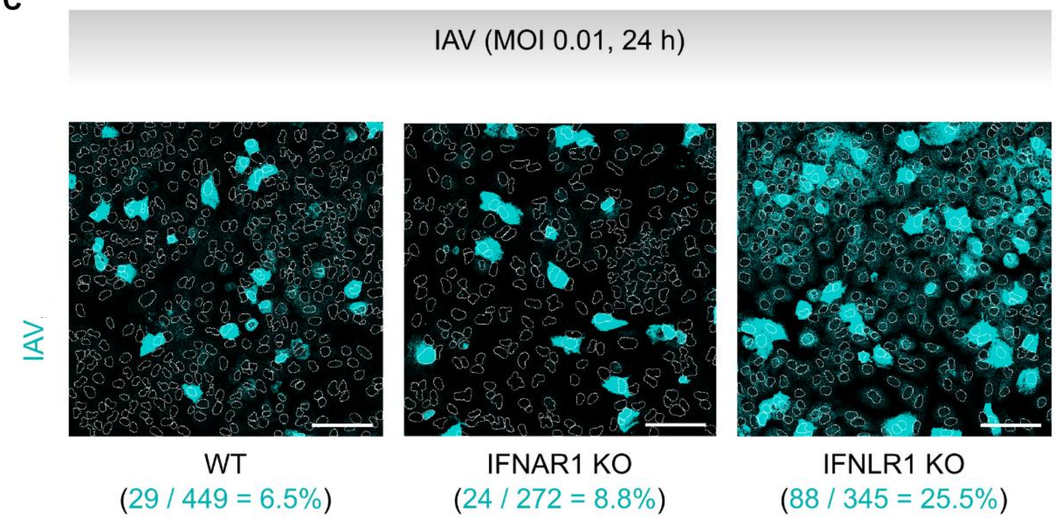


D

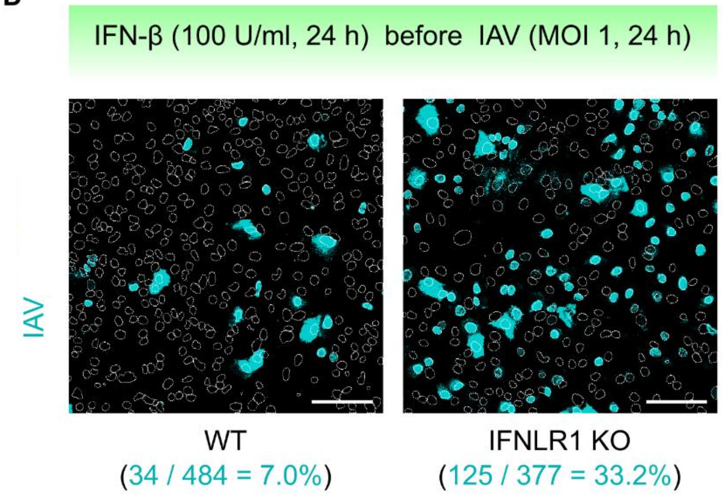

#### **Figure 6A** / rep1 (**IAV** in IFNLR1 KO)

Replicate 1,
quantified for Fig. 6A

(*in addition to replicates quantified for Fig. 4B*)

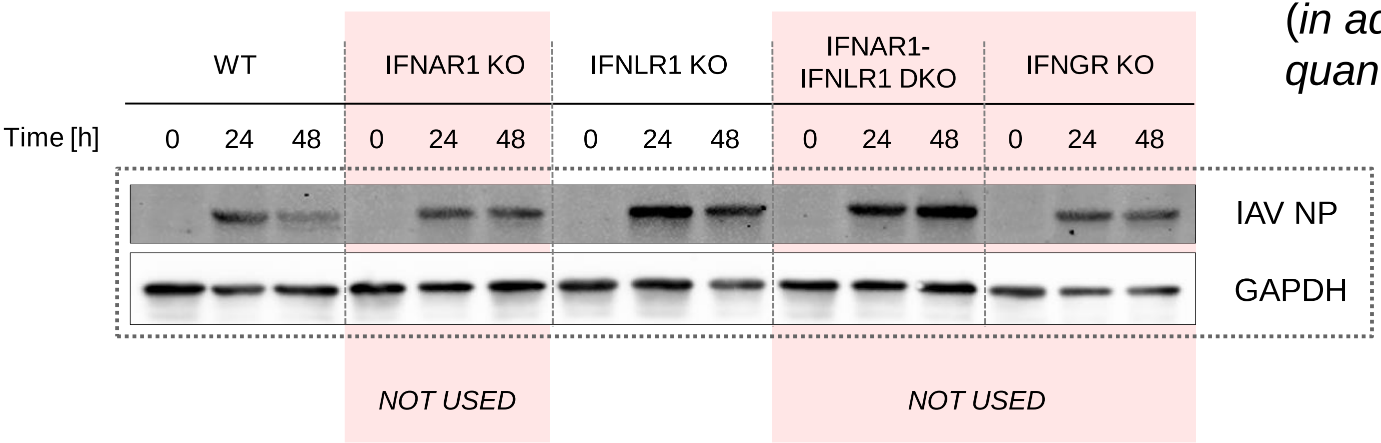

### **Figure 6A** / rep2 (**IAV** in IFNLR1 KO)

Replicate 2,
quantified for Fig. 6A

(*in addition to replicates quantified for Fig. 4B*)

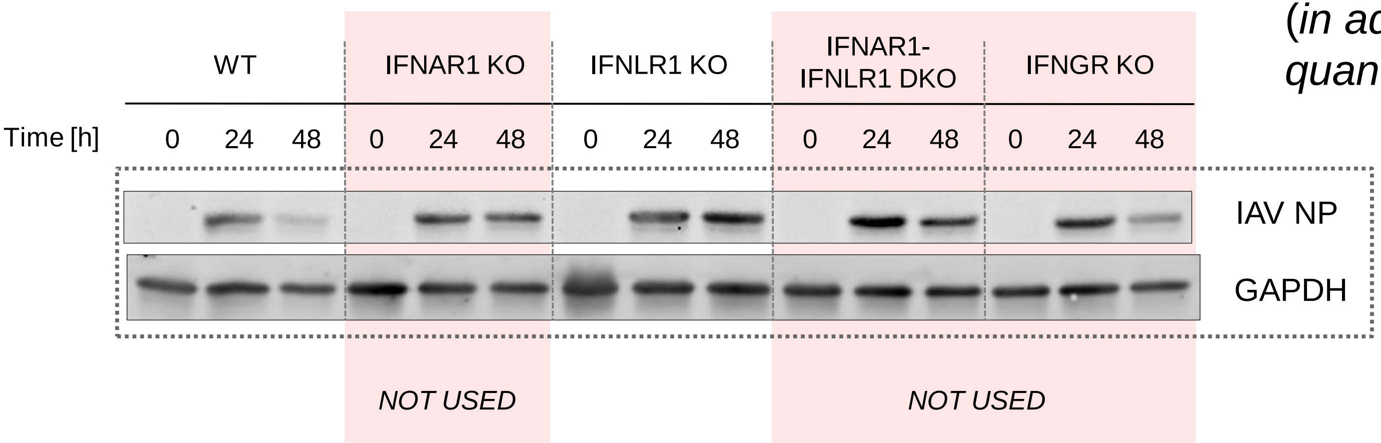

**Figure 6A** / rep3 (**IAV** in IFNLR1 KO)

Replicate 3,
quantified for Fig. 6A

(*in addition to replicates quantified for Fig. 4B*)

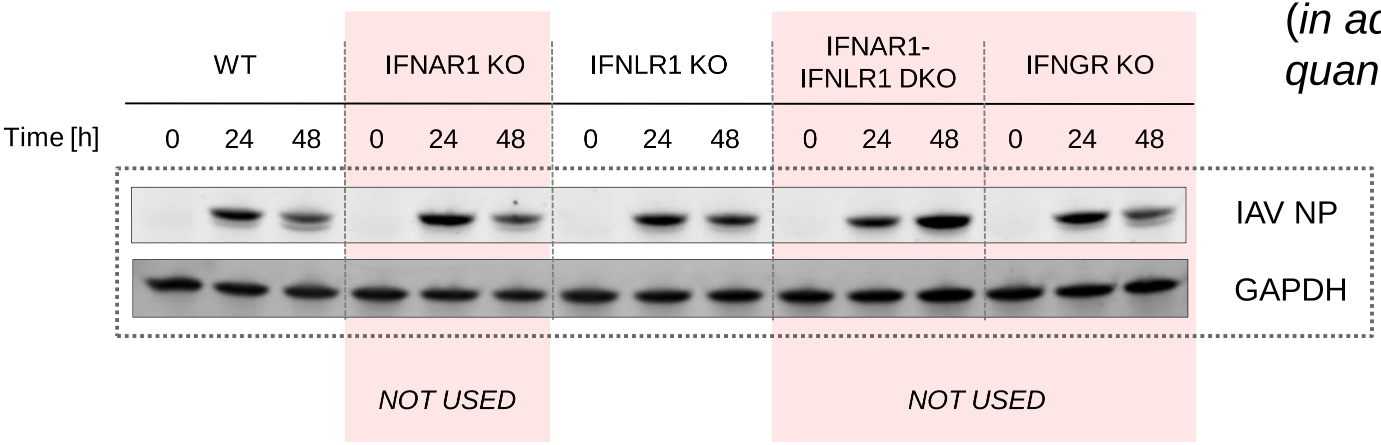

**Figure 6A** / Supplementary Table FACS

| Experiment | MOI | Time | Fraction dead/dying IFNLR1-KO | Fraction dead/dying WT |
|---|---|---|---|---|
| **rep1** | **0.1** | **24h** | 19.40% | 15.07% |
| | | **48h** | 24.02% | 13.65% |
| **rep2** | **0.1** | **24h** | 29.10% | 28.98% |
| | | **48h** | 42.58% | 29.26% |
| **rep3** | **0.1** | **24h** | 14.83% | 16.21% |
| | | **48h** | 17.51% | 19.98% |
| **rep4** | **0.1** | **24h** | 16.22% | 15.67% |

**Figure 6A** / Supplementary Table — Western blot quantifications **Part 1**

| Experiment | MOI | Time | IAV NP level IFNLR1-KO | IAV NP level WT |
|---|---|---|---|---|
| **3B-rep1** | **0.1** | **24h** | 78.19 | 32.87 |
| | | **48h** | 100.00 | 21.71 |
| **3B-rep2** | **0.1** | **24h** | 59.68 | 28.15 |
| | | **48h** | 78.79 | 21.07 |
| **3B-rep3** | **0.1** | **24h** | 58.20 | 39.48 |
| | | **48h** | 93.78 | 31.18 |
| **3B-rep4** | **0.1** | **24h** | 64.94 | 29.54 |
| | | **48h** | 42.18 | 25.24 |
| **3B-rep5** | **0.1** | **24h** | 89.20 | 92.24 |
| | | **48h** | 100.00 | 50.46 |

**Figure 6A** / Supplementary Table — Western blot quantifications **Part 2**

| Experiment | MOI | Time | IAV NP level IFNLR1-KO | IAV NP level WT |
|---|---|---|---|---|
| 3B-rep6 | 0.1 | 24h | 98.26 | 46.78 |
| | | 48h | 100.00 | 29.67 |
| 3B-rep7 | 0.1 | 24h | 66.41 | 14.20 |
| | | 48h | 100.00 | 20.63 |
| 3B-rep8 | 0.1 | 24h | 75.17 | 13.92 |
| | | 48h | 100.00 | 20.60 |
| 6A-rep1 | 0.1 | 24h | 100.00 | 59.85 |
| | | 48h | 93.09 | 24.52 |
| 6A-rep2 | 0.1 | 24h | 64.00 | 56.85 |
| | | 48h | 79.38 | 23.62 |
| 6A-rep3 | 0.1 | 24h | 82.25 | 83.83 |
| | | 48h | 66.15 | 67.08 |

**Figure 6B** / Supplementary Table — Imaging

| Experiment | Cell Line | Time | Fraction Virus Positive | | Cumulated Deaths Fraction | |
|---|---|---|---|---|---|---|
| | | | IAV | RSV | IAV | RSV |
| rep1 | IFNLR1-KO | 24h | 5.59% | 38.63% | 7.63% | 6.68% |
| | | 48h | 7.73% | 71.96% | 41.38% | 41.51% |
| | WT | 24h | 1.28% | 47.52% | 3.37% | 2.81% |
| | | 48h | 0.91% | 69.95% | 17.31% | 25.37% |
| rep2 | IFNLR1-KO | 24h | 4.71% | 14.57% | 4.11% | 3.89% |
| | | 48h | 4.53% | 54.76% | 13.42% | 14.87% |
| | WT | 24h | 2.03% | 18.31% | 4.67% | 4.98% |
| | | 48h | 1.92% | 64.40% | 12.00% | 10.52% |

**Figure 7C and D** / rep2

(Caspase 3/7 activation in dying cells)

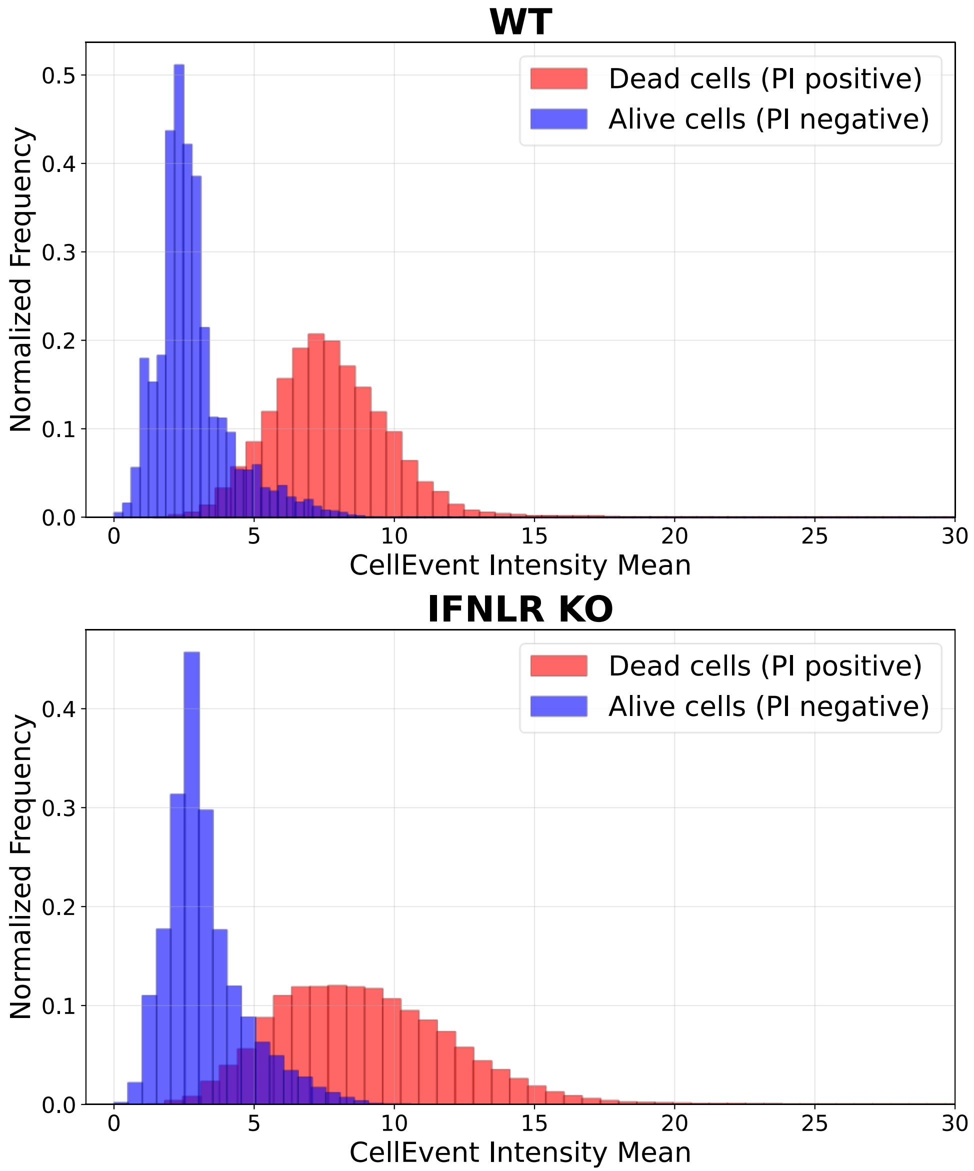

**Figure 7C and D** / rep3

(Caspase 3/7 activation in dying cells)

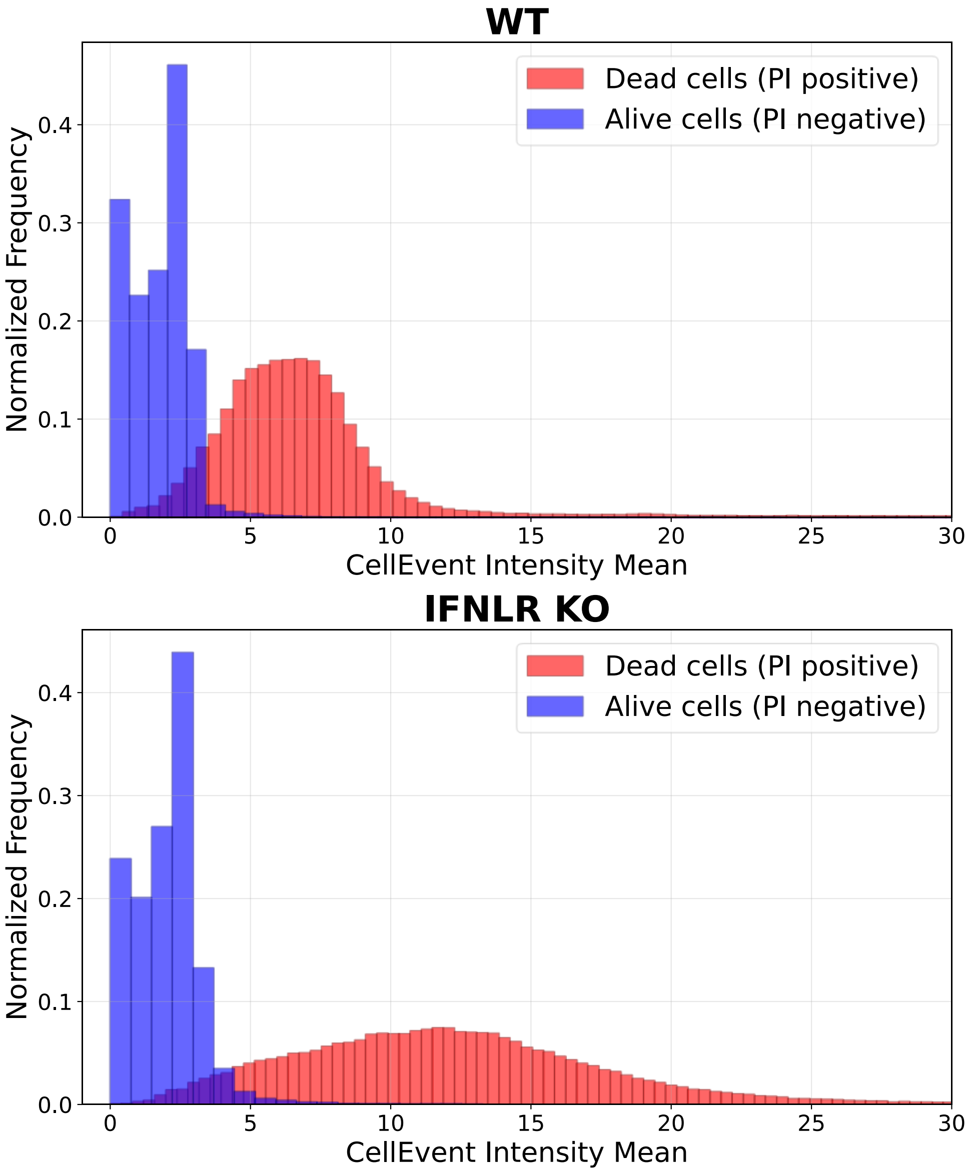

**Figure S1B** / rep1 (STAT1/2 & ISGs after IFN-β and IFN-λ1)

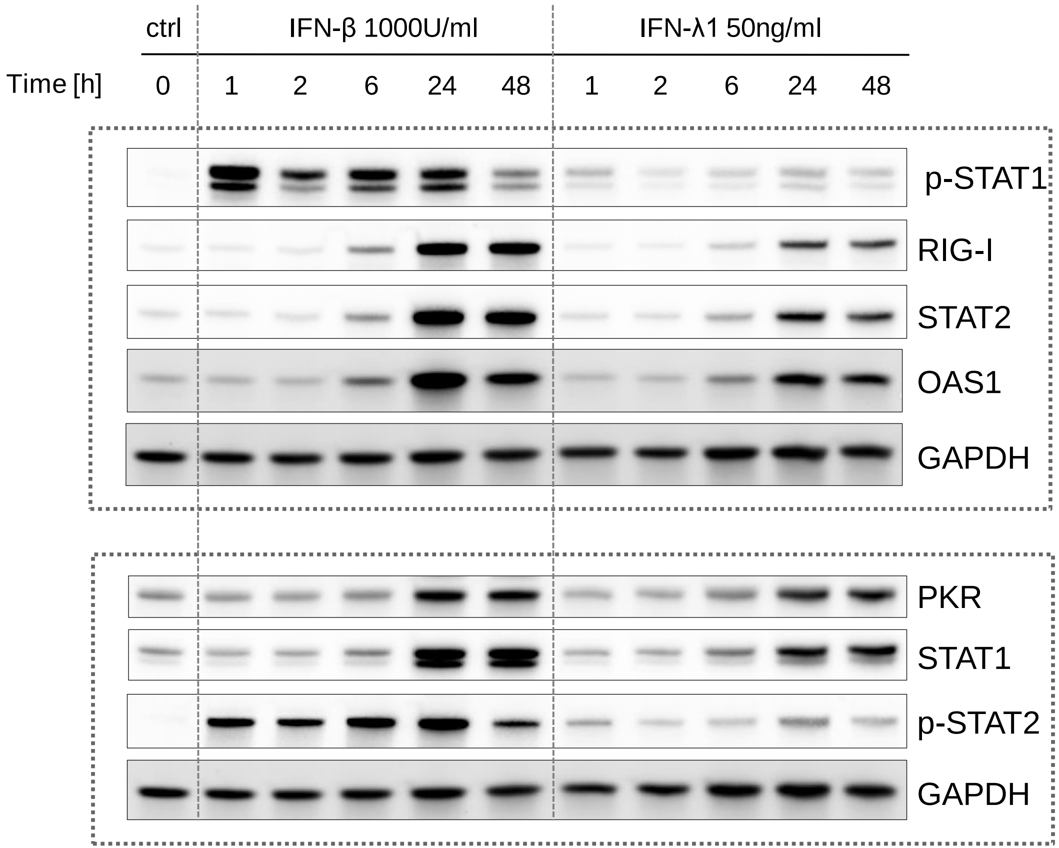


Replicate 1, quantified for Fig. S1B

**Figure S1B** / rep2 (STAT1/2 & ISGs after IFN-β and IFN-λ1)

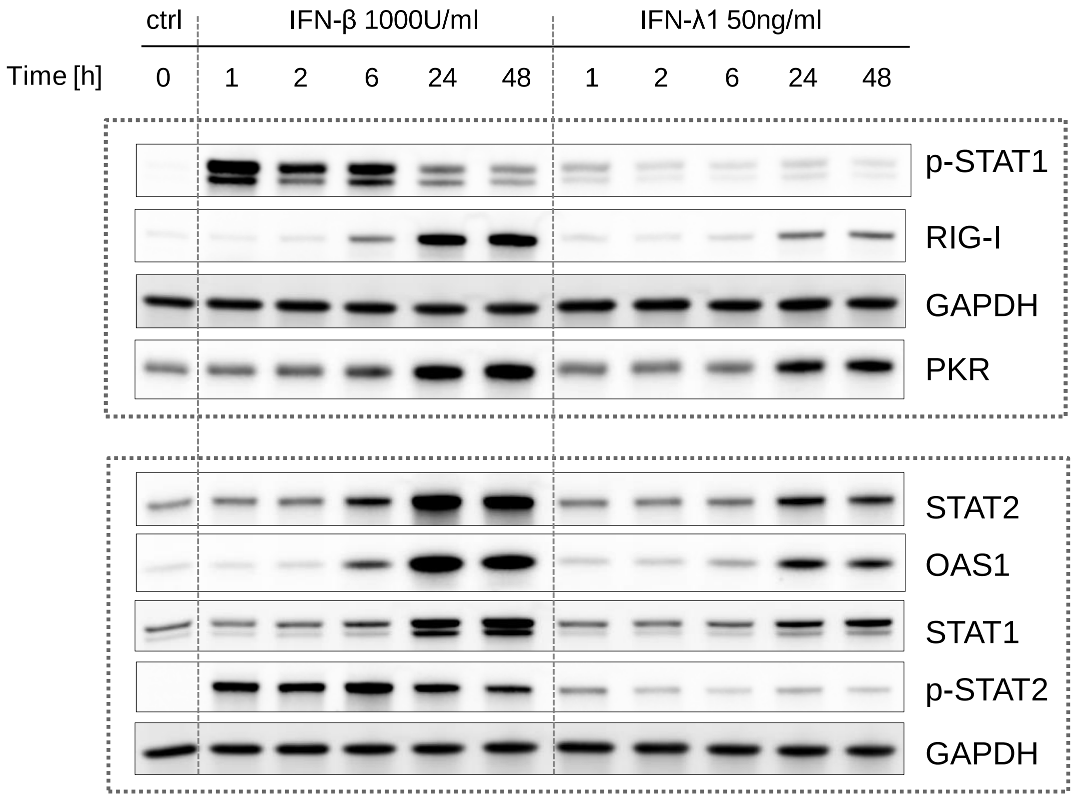


Replicate 2,
quantified for Fig. S1B

### **Figure S1B** / rep3 (STAT1/2 & ISGs after IFN-β and IFN-λ1)

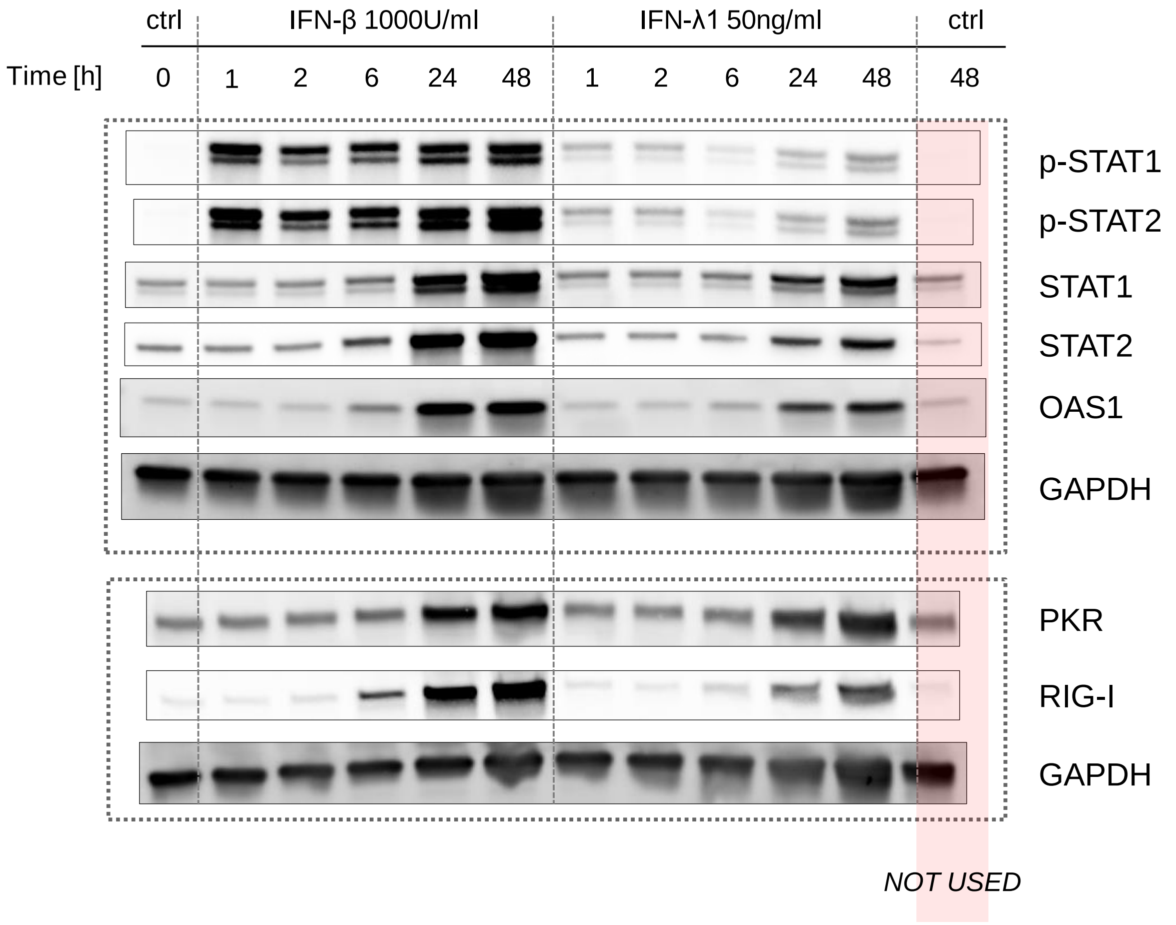


Replicate 3,
quantified for Fig. S1B

**Figure S2C** / rep1 (IFN-λ1 doses)

Replicate 1,
quantified for Fig. S2C

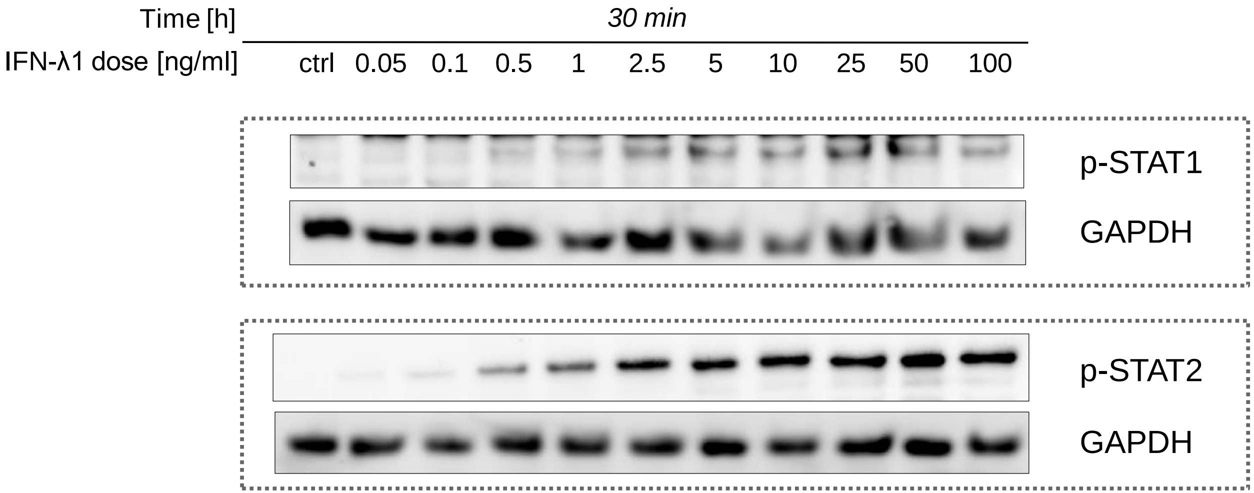

### **Figure S2C** / rep2 (IFN-λ1 doses)

Replicate 2,
quantified for Fig. S2C

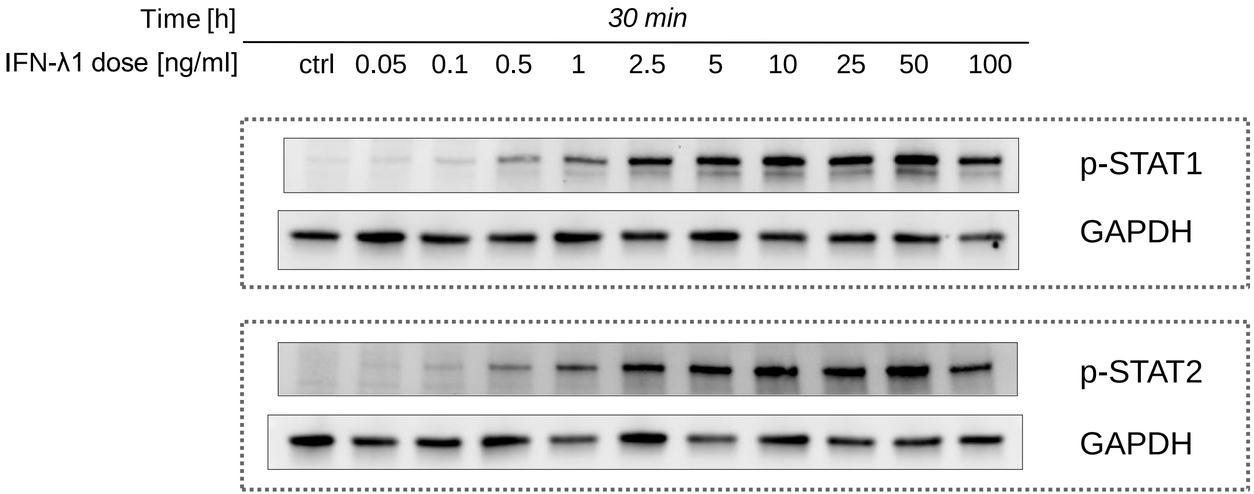

**Figure S2D** / rep1 & rep2 (IFN-β doses)

Replicate 1 and 2 quantified for Fig. S2D

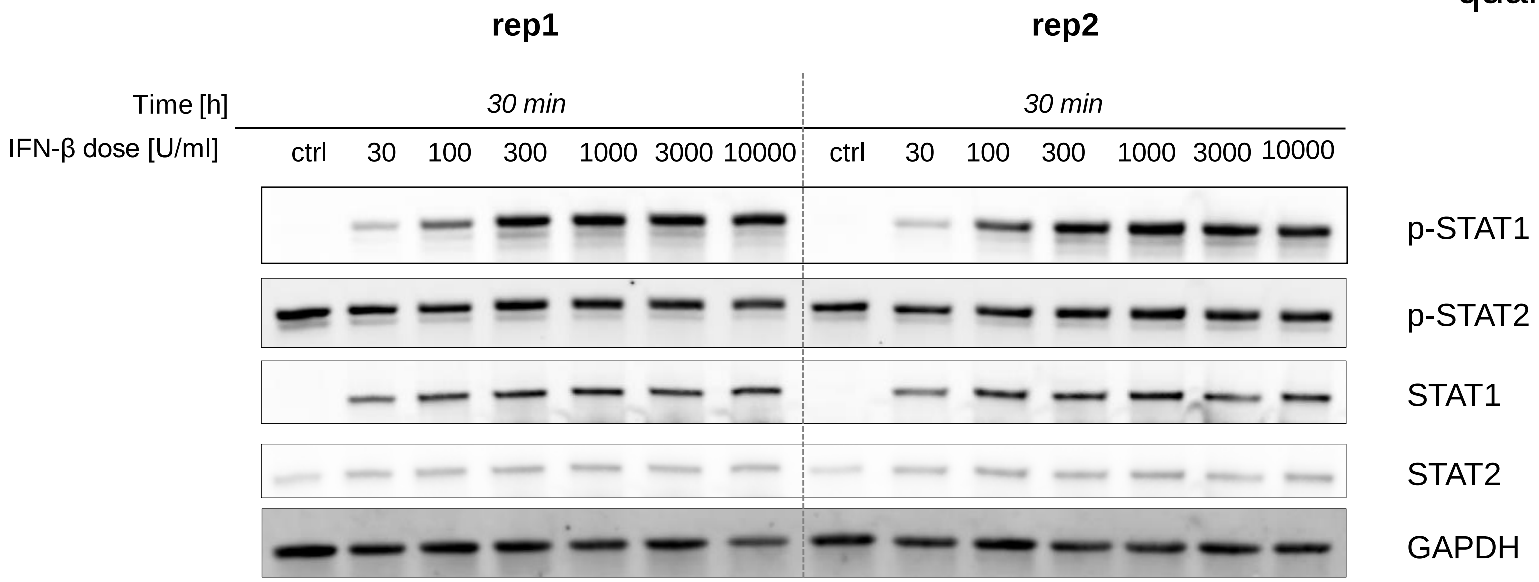

**Figure S7** / rep1 (**IAV** in IFN_R KOs)

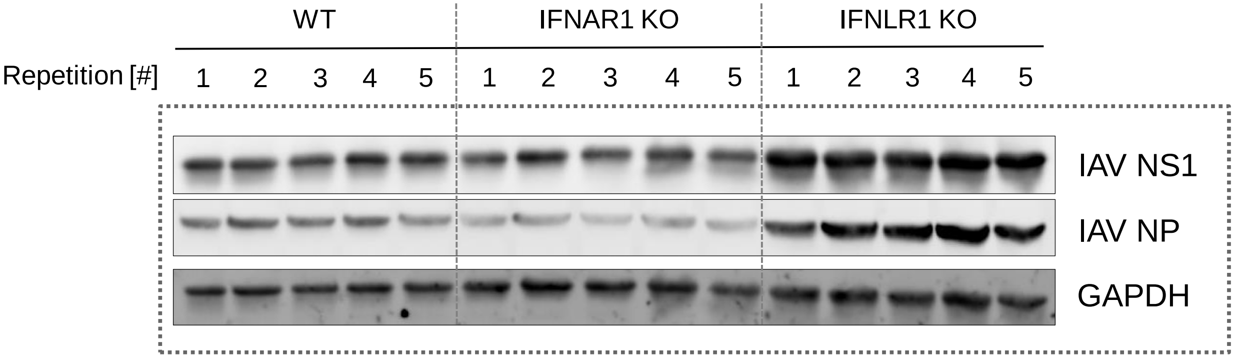


Replicate 1,
quantified for Figs. S7

**Figure S7** / rep2 (**IAV** in IFN_R KOs)

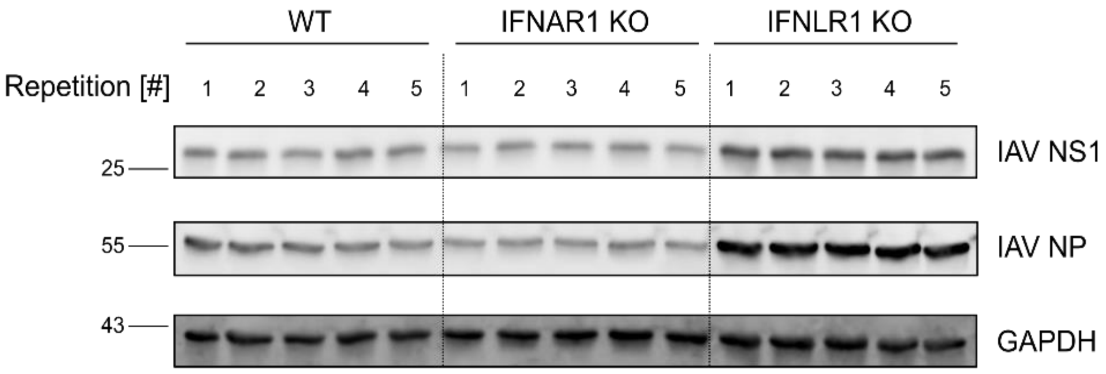


Replicate 2,
quantified for Figs. S7

**Figure S7** / rep3 (**IAV** in IFN_R KOs)

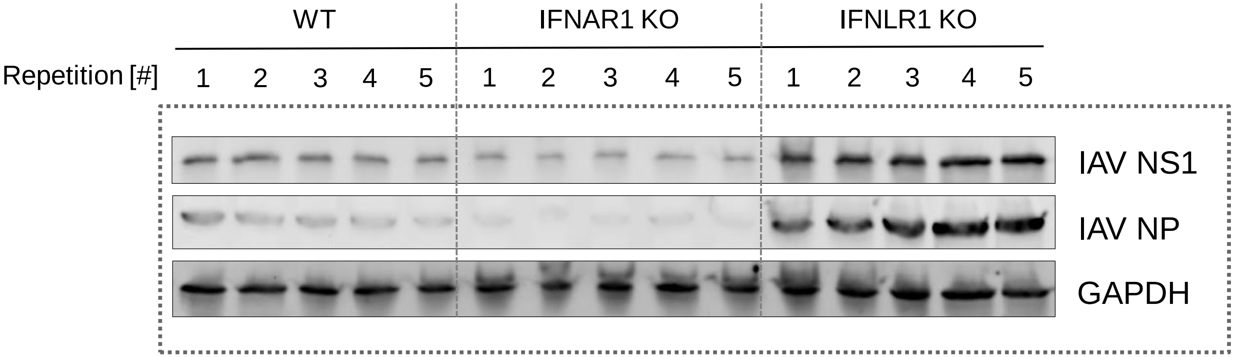


Replicate 3,
quantified for Figs. S7

**Figure S9A** / rep1 (**IAV** in WT and IFNLR1 KO)

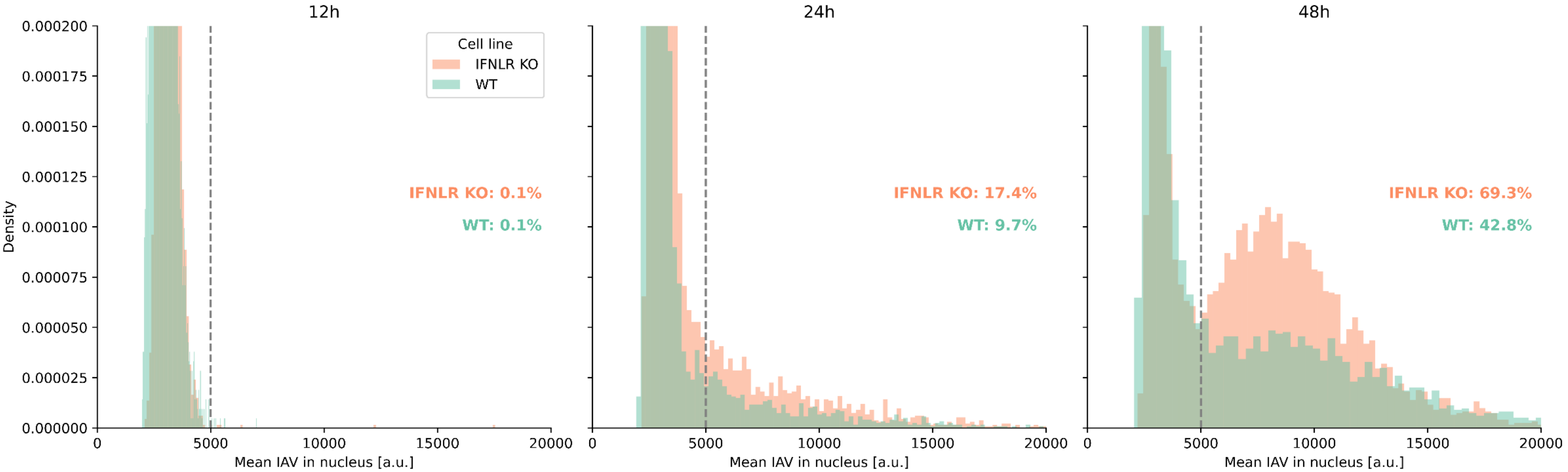

**Figure S9A** / rep2 (**IAV** in WT and IFNLR1 KO)

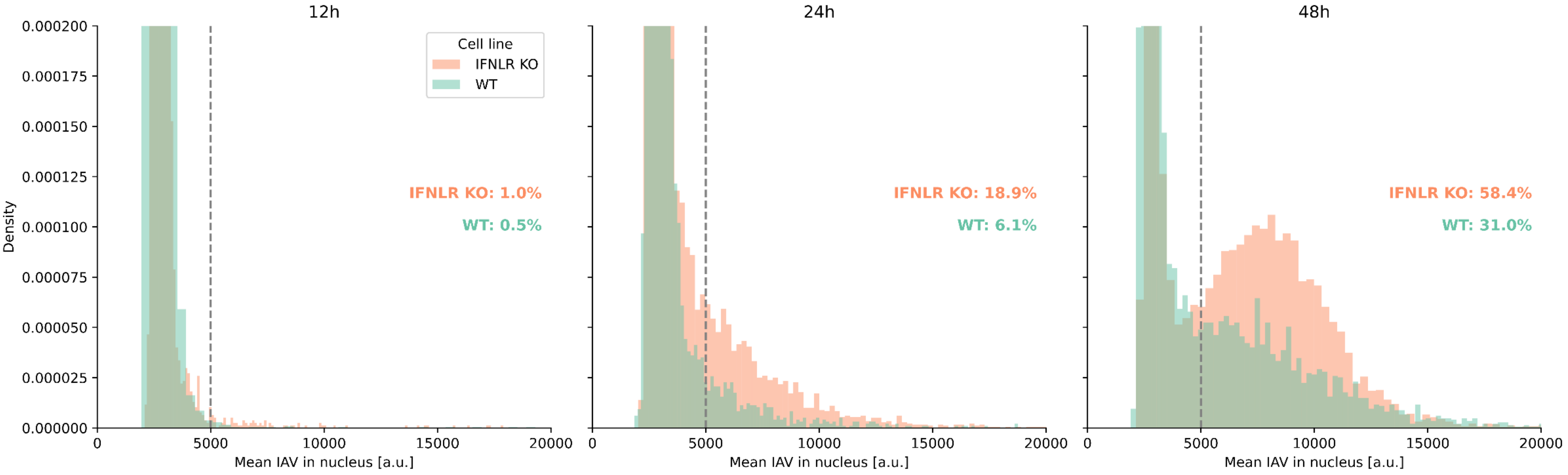

### **Figure S9A** / rep3 (**IAV** in WT and IFNLR1 KO)

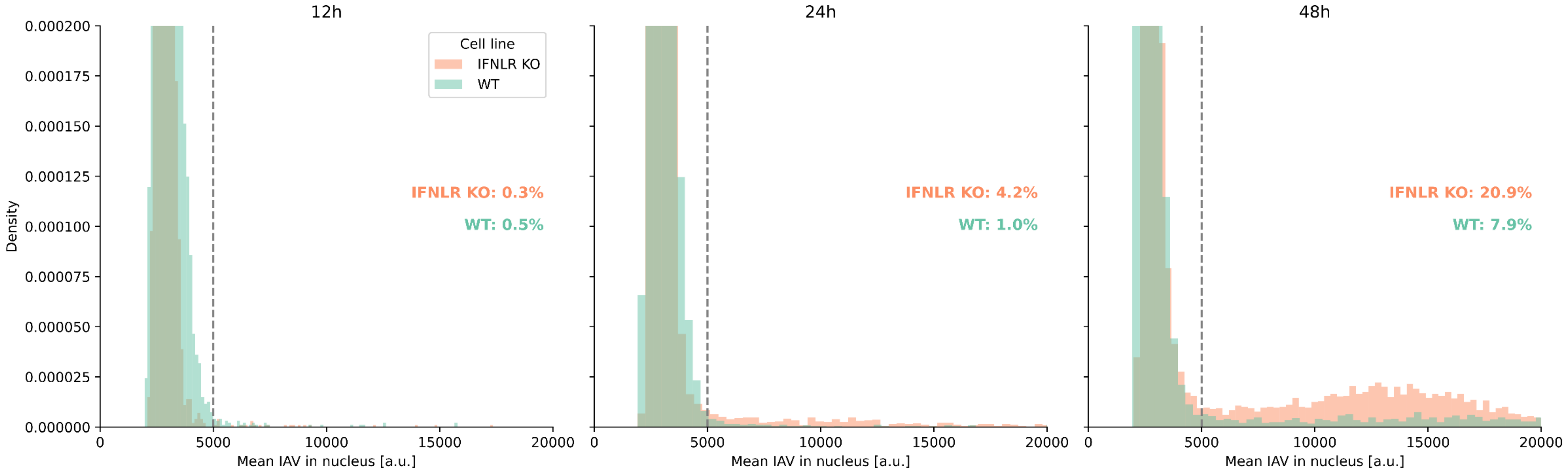

**Figure S9B** / rep1 (**IAV** in WT and IFNLR1 KO)

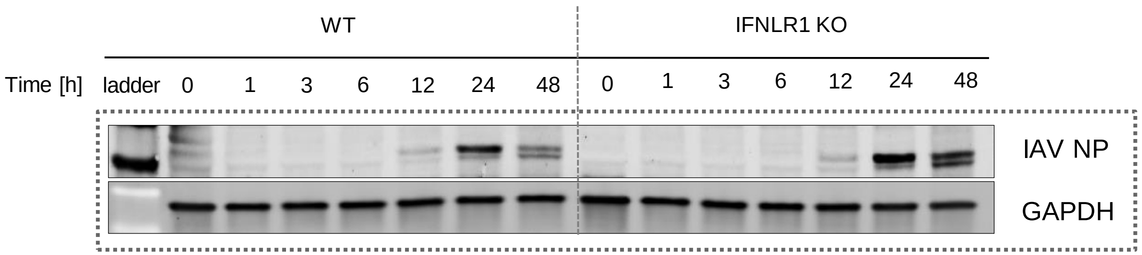

**Figure S9B** / rep1 (**IAV** in WT and IFNLR1 KO)

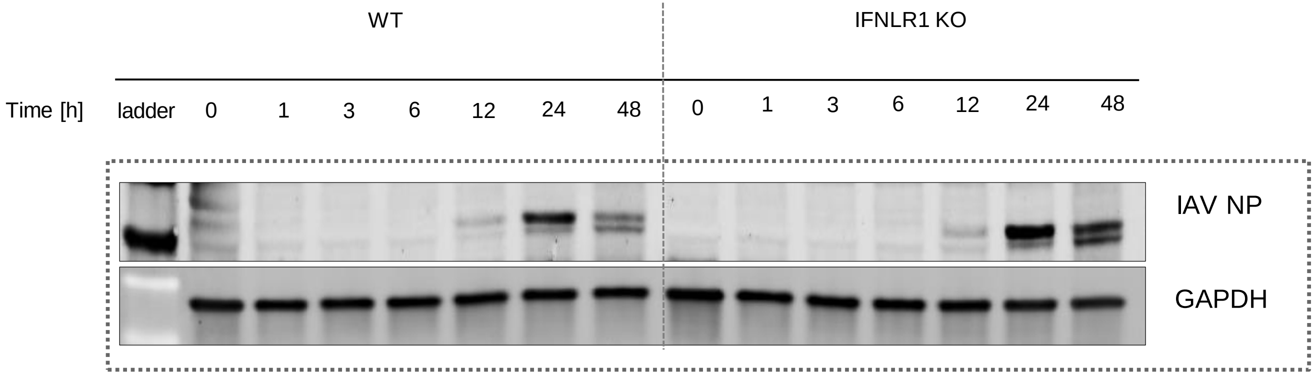

**Figure S9B** / rep2 (**IAV** in WT and IFNLR1 KO)

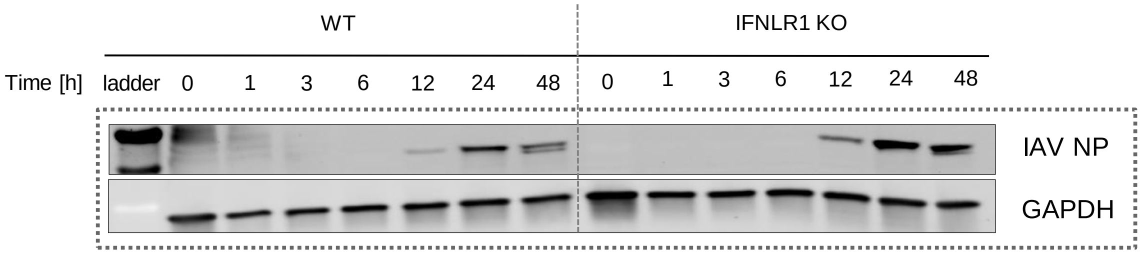

**Figure S9B** / rep3 (**IAV** in WT and IFNLR1 KO)

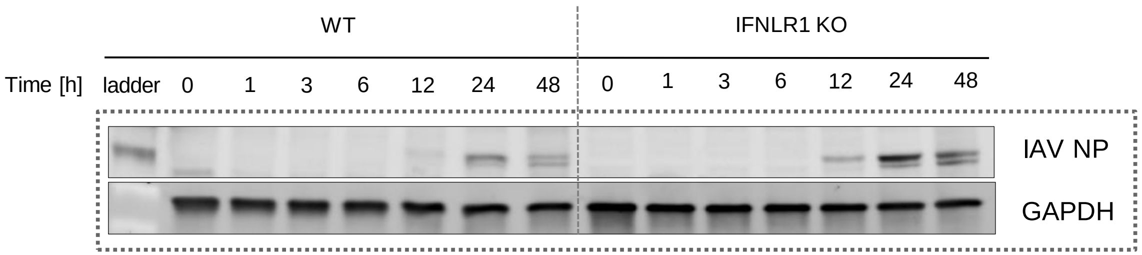

### **Figure S9B** / rep4 (**IAV** in WT and IFNLR1 KO)

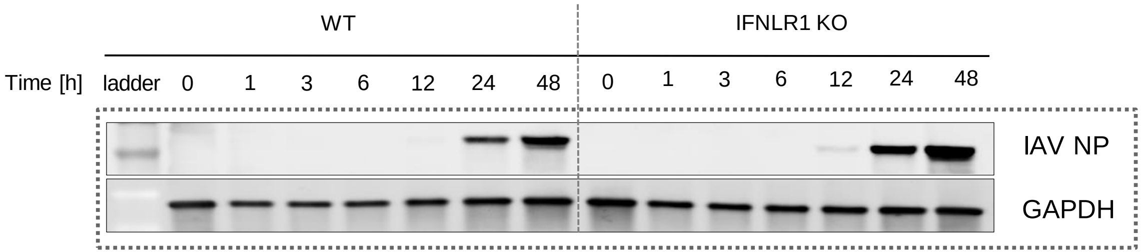

**Figure S9C**/ Supplementary Table — Western blot quantifications

| Experiment | MOI | Time | IAV NP level IFNLR1-KO | IAV NP level WT |
|---|---|---|---|---|
| **S8B-rep1** | **0.1** | **12h** | 7.95 | 7.16 |
| | | **24h** | 100.00 | 52.2 |
| | | **48h** | 79.42 | 33.17 |
| **S8B-rep2** | **0.1** | **12h** | 19.42 | 3.95 |
| | | **24h** | 100.00 | 27.1 |
| | | **48h** | 69.59 | 30.22 |
| **S8B-rep3** | **0.1** | **12h** | 17.44 | 4.79 |
| | | **24h** | 100.00 | 35.48 |
| | | **48h** | 76.98 | 27.56 |
| **S8B-rep4** | **0.1** | **12h** | 1.80 | 0.37 |
| | | **24h** | 51.35 | 22.75 |
| | | **48h** | 100.00 | 40.72 |
