## Supplementary Data S2 for "Type III interferons may suppress viral infections by triggering cell death": Figure 1 plot polyIC.ipynb

```
# %% [code]
from pathlib import Path

import numpy as np
import pandas as pd

from scipy.stats import levene
from scipy.interpolate import interp1d

import matplotlib.pyplot as plt
import matplotlib.ticker as ticker
from matplotlib.lines import Line2D

from r_stats import r_anova_and_tukey

# %% [code]
SOURCE_DATA_DIR = Path('./Quantifications')

# %% [code]
!mkdir -p "Figures/Figure 1"

# %% [code]
def format_time(x, pos='None'):
    h, rem = divmod(x, 60)
    m = int(rem)
    if h >= 1:
        return f"{int(h)}h {m}m" if m else f"{int(h)}h"
    else:
        return f"{m}m"

# %% [code]
def plot_all_experiments(deaths):
    # Cell type (color)
    celltype_legend = [
        Line2D([0], [0], color='blue', marker='s', linestyle='', label='WT'),
        Line2D([0], [0], color='red', marker='s', linestyle='', label='IFNLR1 KO'),
    ]

    spacer = Line2D([0], [0], color='white', linestyle='')

    # pIC concentration (line width)
    pIC_legend = [
        Line2D([0], [0], color='black', linewidth=2, label='poly(I:C) = 0.1 µg/ml'),
        Line2D([0], [0], color='black', linewidth=1, label='poly(I:C) = 0.03 µg/ml'),
        Line2D([0], [0], color='black', linewidth=0.5, label='poly(I:C) = 0 µg/ml'),
    ]

    # Costimulation (line style)
    costimulation_legend = [
        Line2D([0], [0], color='black', linestyle='--', label='Costimulation with IFNβ (1000 U/ml)'),
        Line2D([0], [0], color='black', linestyle='-', label='No costimulation'),
    ]

    # Combine with spacers
    full_legend = (
        celltype_legend + [spacer] +
        pIC_legend + [spacer] +
        costimulation_legend
    )


    celltype_color_map = {
        "A549 WT": "blue",
        "A549 IFNLR KO": "red",
    }

    prestimulation_linestyle_map = {
        "mock": "-",
        "IFNb 1000 u/ml": "--",
    }

    pIC_linewidth_map = {
        "mock": 0.5,
        "pIC 0,03 ug/ml": 1,
        "pIC 0,1 ug/ml": 2,
    }
    
    # Create figure and subplots
    fig, axs = plt.subplots(1, 3, figsize=(15, 5), sharex=True, sharey=True)
    for rep, deaths_group in enumerate(deaths.groupby('experiment')):
        _, deaths = deaths_group
        for well, group in deaths.groupby(['well', 'cellline', 'stimulation', 'pIC']):
            color = celltype_color_map.get(well[1], "black")
            linestyle = prestimulation_linestyle_map.get(well[2], None)
            linewidth = pIC_linewidth_map.get(well[3], None)

            axs[rep].plot(
                group['time'],
                group['cumulated_deaths'],
                linestyle=linestyle,
                linewidth=linewidth,
                color=color,
            )

        axs[rep].set_title(f"Replicate {rep+1}", fontsize=10)
        
        tick_hours = [0, 180, 360, 540, 900, 720, 1080]
        axs[rep].set_xticks(tick_hours)
        axs[rep].set_xlabel("Time", fontsize=10)
        axs[rep].xaxis.set_major_formatter(ticker.FuncFormatter(format_time))

    # Add labels and titles
    axs[0].set_ylabel("Cumulative dying cell fraction", fontsize=10)
    
    # Add to first subplot
    axs[0].legend(handles=full_legend, loc='upper left', frameon=False, fontsize=9)
    
    # Add vertical line
    fig.lines.append(plt.Line2D(
        xdata=[0.140, 0.140],  
        ydata=[0.86, 0.91],  
        transform=fig.transFigure,
        color='black',
        linewidth=1,
    ))

    # Top horizontal cap
    fig.lines.append(plt.Line2D(
        xdata=[0.140, 0.137],
        ydata=[0.91, 0.91],
        transform=fig.transFigure,
        color='black',
        linewidth=1,
    ))

    # Bottom horizontal cap
    fig.lines.append(plt.Line2D(
        xdata=[0.140, 0.137],
        ydata=[0.86, 0.86],
        transform=fig.transFigure,
        color='black',
        linewidth=1,
    ))
    
    # Add stars above the line
    fig.text(0.150, 0.88, '**', ha='center', va='center', fontsize=9)

    return fig, axs

# %% [code]
data =  pd.read_csv(SOURCE_DATA_DIR / 'Imaging/Figure 1/all_reps.csv', dtype={'well': str})
data

# %% [code]
fig, axs = plot_all_experiments(data)

plt.tight_layout()
fig.savefig('./Figures/Figure 1/figure_1--polyIC_WT_vs_IFNLR_KO.svg')

# %% [markdown]
# Revision

# %% [code]
def add_significance(fig, x, y):
        # Add vertical line
    fig.lines.append(plt.Line2D(
        xdata=[x, x],  
        ydata=[y, y+0.07],  
        transform=fig.transFigure,
        color='black',
        linewidth=1,
    ))

    # Top horizontal cap
    fig.lines.append(plt.Line2D(
        xdata=[x, x-0.003],
        ydata=[y+0.07, y+0.07],
        transform=fig.transFigure,
        color='black',
        linewidth=1,
    ))

    # Bottom horizontal cap
    fig.lines.append(plt.Line2D(
        xdata=[x, x-0.003],
        ydata=[y, y],
        transform=fig.transFigure,
        color='black',
        linewidth=1,
    ))
    # Add stars above the line
    fig.text(x+0.01, y+0.03, '**', ha='center', va='center', fontsize=9)
    return fig

def plot_one_experiment(deaths):
    # Cell type (color)
    celltype_legend = [
        Line2D([0], [0], color='blue', marker='s', linestyle='', label='WT'),
        Line2D([0], [0], color='red', marker='s', linestyle='', label='IFNLR1 KO'),
    ]

    spacer = Line2D([0], [0], color='white', linestyle='')

    # pIC concentration (line width)
    pIC_legend = [
        Line2D([0], [0], color='black', linewidth=2, label='poly(I:C) = 0.1 µg/ml'),
        Line2D([0], [0], color='black', linewidth=1, label='poly(I:C) = 0.03 µg/ml'),
        Line2D([0], [0], color='black', linewidth=0.5, label='mock'),#'poly(I:C) = 0 µg/ml'),
    ]

    # Combine with spacers
    full_legend = (
        celltype_legend + [spacer] +
        pIC_legend + [spacer]
    )


    celltype_color_map = {
        "A549 WT": "blue",
        "A549 IFNLR KO": "red",
    }


    pIC_linewidth_map = {
        "mock": 0.5,
        "pIC 0,03 ug/ml": 1,
        "pIC 0,1 ug/ml": 2,
    }
    
    # Create figure and subplots
    fig, axs = plt.subplots(1, 2, figsize=(10, 5), sharex=True, sharey=True)
    for rep, deaths_group in enumerate(deaths.groupby('stimulation')):
        stimulation, deaths = deaths_group
        for well, group in deaths.groupby(['well', 'cellline', 'pIC']):
            color = celltype_color_map.get(well[1], "black")
            linewidth = pIC_linewidth_map.get(well[2], None)

            axs[rep].plot(
                group['time'],
                group['cumulated_deaths'],
                linewidth=linewidth,
                color=color,
            )
        if stimulation=='mock':
            axs[rep].set_title(f"No costimulation", fontsize=10)
        else:
            axs[rep].set_title(f"Costimulation with IFNβ (1000 U/ml)", fontsize=10)
        
        tick_hours = [0, 180, 360, 540, 900, 720, 1080]
        axs[rep].set_xticks(tick_hours)
        axs[rep].set_xlabel("Time", fontsize=10)
        axs[rep].xaxis.set_major_formatter(ticker.FuncFormatter(format_time))

    # Add labels and titles
    axs[0].set_ylabel("Cumulative dying cell fraction", fontsize=10)
    
    # Add to first subplot
    axs[0].legend(handles=full_legend, loc='upper left', frameon=False, fontsize=9)
    axs[1].legend(handles=full_legend, loc='upper left', frameon=False, fontsize=9)
    

    add_significance(fig, 0.25, 0.79)
    add_significance(fig, 0.68, 0.79)
    

    return fig, axs

fig, axs = plot_one_experiment(data[data['experiment']=='rep1'])
#fig.savefig('/mnt/nas/group/PAPER/LambdaPtosis/Manuscript_Frontiers/Prus_et_al_2025__Source_Data_S2/Prus_et_al_2025_v3--Source_Data_S2/Figures/Figure 1/figure1-revision-rep1.svg')

# %% [code]
fig, axs = plot_one_experiment(data[data['experiment']=='rep2'])
fig.savefig('/mnt/nas/group/PAPER/LambdaPtosis/Manuscript_Frontiers/Prus_et_al_2025__Source_Data_S2/Prus_et_al_2025_v3--Source_Data_S2/Figures/Figure 1/figure1-revision-rep2.svg')
fig, axs = plot_one_experiment(data[data['experiment']=='rep3'])
fig.savefig('/mnt/nas/group/PAPER/LambdaPtosis/Manuscript_Frontiers/Prus_et_al_2025__Source_Data_S2/Prus_et_al_2025_v3--Source_Data_S2/Figures/Figure 1/figure1-revision-rep3.svg')

# %% [markdown]
# Statistical test

Run ANOVA on the cumulated number of deaths with factors: CellType, stimulation, pIC. \
Then, repeat it on some (or all) timepoints after the first half to ensure robustness of the result.


# %% [code]
def interpolate_deaths(df, t):
    f = interp1d(
        df['time'],
        df['cumulated_deaths'],
        bounds_error=False,
    )
    return float(f(t))

def df_with_interpolated_deaths(df, t):
    cumulated_deaths_at_t = df.groupby([
            'experiment',
            'well',
            'cellline',
            'pIC',
            'stimulation',
    ]).apply(lambda dg: interpolate_deaths(dg, t=t), include_groups=False)

    return cumulated_deaths_at_t.to_frame('cumulated_deaths').reset_index()

# example: interpolate cumulated_deaths at 16h
df_with_interpolated_deaths(data, t=16*60)

# %% [code]
def compute_pvals_over_time(
    data, 
    formula, 
    factors, 
    groupby_cols,
    times,
):
    """
    Compute p-values over time using ANOVA.

    Parameters:
    - data: Input DataFrame
    - formula: ANOVA formula string
    - factors: List of factors to extract p-values for (e.g., ['cellline', 'pIC'])
    - groupby_cols: List of columns to groupby before averaging (e.g., ['experiment', 'cellline', 'pIC'])
    - times: times at which to evaluate

    Returns:
    - pvals: Dictionary with p-values for each factor and time
    """
    pvals = {factor: [] for factor in factors}
    pvals['time'] = []
    
    for t in times:
        # Interpolate deaths at time t (for each experiment and well separately)
        df = df_with_interpolated_deaths(data, t=t)

        # Aggregate
        data_test = df.groupby(groupby_cols)['cumulated_deaths'].mean().reset_index()

        # Check Levene
        stat, p = levene(*data_test.groupby(factors)['cumulated_deaths'].agg(list))
        if p < 0.05:
            print(f"WARNING: Levene rejected at t = {t/60}h with p = {p:.2e}")

        try:
            # Perform ANOVA
            result_anova, _ = r_anova_and_tukey(
                formula,
                factors,
                factors[0],  # Main factor for posthoc
                data_test
            )

            # Store p-values
            for factor in factors:
                if factor in result_anova.index:
                    pvals[factor].append(result_anova.loc[factor, 'Pr(>F)'])
                else:
                    pvals[factor].append(np.nan)
            pvals['time'].append(t)

        except Exception as e:
            print(f"Error at t={t}: {e}")
            for factor in factors:
                pvals[factor].append(np.nan)
            pvals['time'].append(t)

    return pvals

# %% [code]
# Interpolate to times of the shortest experiment
shortest_experiment = data.groupby('experiment')['time'].max().idxmin()
times = sorted(data[data['experiment'] == shortest_experiment]['time'].unique())

# %% [code]
%%time

pvals_full = compute_pvals_over_time(
    data=data[((data['pIC'] != 'mock') | (data['stimulation'] != 'mock'))],
    formula="cumulated_deaths ~ cellline*pIC*stimulation",
    factors=["cellline", "pIC", "stimulation"],
    groupby_cols=["experiment", "cellline", "pIC", "stimulation"],
    times=times,
)

# %% [code]
%%time

pvals_filtered = compute_pvals_over_time(
    data=data[((data['pIC'] != 'mock') | (data['stimulation'] != 'mock')) & (data['stimulation'] != 'mock')],
    formula="cumulated_deaths ~ cellline*pIC",
    factors=["cellline", "pIC"],
    groupby_cols=["experiment", "cellline", "pIC"],
    times=times,
)

# %% [code]
%%time

pvals_mockonly = compute_pvals_over_time(
    data=data[((data['pIC'] != 'mock') | (data['stimulation'] != 'mock')) & (data['stimulation'] == 'mock')],
    formula="cumulated_deaths ~ cellline*pIC",
    factors=["cellline", "pIC"],
    groupby_cols=["experiment", "cellline", "pIC"],
    times=times,
)

# %% [code]
fig, axs = plt.subplots(1, 2, figsize=(16, 6), sharey=True)
colorfactor = {'cellline': 'blue', 'pIC': 'orange', 'stimulation': 'green'}


for factor in ["cellline", "pIC", "stimulation"]:
    axs[0].plot(pvals_full["time"], pvals_full[factor], label=factor)

axs[0].axhline(0.05, color="red", linestyle="--", label="p = 0.05")
axs[0].set_yscale("log")
axs[0].set_xlabel("Time")
axs[0].set_ylabel("p-value")
axs[0].set_title("ANOVA: cellline * pIC * stimulation")
axs[0].legend()
axs[0].grid(True)
axs[0].xaxis.set_major_formatter(ticker.FuncFormatter(format_time))

for factor in ["cellline", "pIC"]:
    axs[1].plot(pvals_filtered["time"], pvals_filtered[factor], label=factor, color = colorfactor[factor],)
    axs[1].plot(
        pvals_mockonly["time"],
        pvals_mockonly[factor],
        linestyle="--",
        label=f"{factor} (mock only)",
        color = colorfactor[factor],
    )

axs[1].axhline(0.05, color="red", linestyle="--")
axs[1].set_yscale("log")
axs[1].set_xlabel("Time")
axs[1].set_title("ANOVA: cellline * pIC - separately for costimulation")
axs[1].legend()
axs[1].xaxis.set_major_formatter(ticker.FuncFormatter(format_time))
axs[1].grid(True)

plt.tight_layout()
plt.show()

# %% [code]
```
