## Supplementary Data S2 for "Type III interferons may suppress viral infections by triggering cell death": Figure 3, S1, S2, S3, S4, S5 plot Western blot.ipynb

```
# %% [code]
from pathlib import Path

import numpy as np
import pandas as pd

import scipy.stats as ss

import matplotlib as mpl
import matplotlib.pyplot as plt
import matplotlib.patches as mpatches

from subplots_from_axsize import subplots_from_axsize

from utils import load_rep, p_stars
from r_stats import r_anova_and_tukey

# %% [code]
plt.rcParams["font.size"] = 7
color_ns = 'grey'

# %% [code]
SOURCE_DATA_DIR = Path('./Quantifications')

# %% [code]
!mkdir -p "Figures/Figure S3"
!mkdir -p "Figures/Figure 3"
!mkdir -p "Figures/Figure S1"
!mkdir -p "Figures/Figure S2"
!mkdir -p "Figures/Figure S4"
!mkdir -p "Figures/Figure S5"

# %% [markdown]
# blot manipulation utils

# %% [code]
def blot_renorm_max(data, cutoff=0.03):
    data_norm = data.copy().set_index(['Experiment', 'Lane'])

    for col in data_norm:
        if col[0] != 'Measurement':
            continue

        data_norm[col] = data_norm[col] / data_norm[col].groupby(level=0).max()
        data_norm[col] += cutoff
        data_norm[col] = data_norm[col] / data_norm[col].groupby(level=0).max()

    return data_norm.reset_index()

# %% [markdown]
# blot plotting utils

# %% [code]
mol_to_ylabel = {
    mol: mol for mol in [
        'p-STAT1', 'STAT1',
        'p-STAT2', 'STAT2',
        'RIG-I', 'PKR', 'OAS1',
        'IAV nucleoprotein',
    ]
} | {
    'RSV F protein': 'RSV F',
    'IAV nucleoprotein': 'IAV NP',
}

def plot_blots(
    data,
    exps=None,
    mols=None,
    lanes=None,
    lane_labels=None,
    exp_to_color={},
    mol_to_ylim_max={},
    scalex=1.0,
    scaley=1.0,
    scaleoffset=1.0,
    scalebarwidth=1.0,
    **subplots_kwargs,
):
    exps = exps or data['Experiment'].unique()
    mols = mols or [col[1] for col in data if col[0] == 'Measurement']
    lanes = lanes or sorted(data['Lane'].unique())
    lane_labels = lane_labels or [f"lane {l}" for l in lanes]
    
    axx = len(lane_labels) + 1
    axy = 1
    fig, axs = subplots_from_axsize(**dict(
        nrows=len(mols),
        ncols=1,
        axsize=(scalex * 0.25 * axx, scaley * 0.25 * axy),
        left=0.4,
        right=0.05,
        top=0.05,
        hspace=scaley * 0.4,
        bottom=0.35,   
    ) | subplots_kwargs)
    
    for ax, mol in zip(axs, mols):
        ax.set_ylabel(mol_to_ylabel[mol], fontsize=9)

        ylim_max = mol_to_ylim_max.get(mol, 1.33)
        ax.set_ylim(0, ylim_max)
        ax.set_yticks([0, 1, ylim_max] if ylim_max > 1.5 else [0, 1])
        
        for lane_i, lane in enumerate(lanes):
            data_lane = data[(data['Lane'] == lane) & (~data[('Measurement', mol)].isna())]
            offsets = scaleoffset * 0.15 * np.arange(len(data_lane))
            offsets -= offsets.mean()
            for (_, lane), offset in zip(data_lane.iterrows(), offsets):
                color = exp_to_color.get(lane['Experiment'], 'k')
                ax.bar(
                    1 + lane_i + offset,
                    lane[('Measurement', mol)],
                    color=color,
                    width=scalebarwidth * 0.125,
                    zorder=2,
                    clip_on=False,
                )

    for ax in axs:
        ax.set_xlim(0.5, len(lane_labels) + 0.5)
        ax.set_xticks(range(1, 1 + len(lane_labels)))
        ax.set_xticklabels(lane_labels)
        ax.spines[['top', 'right']].set_visible(False)
        #ax.grid(axis='y', ls=':')
        
    return fig, axs

# %% [code]
def plot_groups(
    axs,
    start_ends=[],
    margin=0.25,
    **kwargs
):
    for ax in axs:
        ax.spines['bottom'].set_color('lightgray')
        kwargs = dict(color='black', lw=1, clip_on=False, zorder=10) | kwargs
        for start, end in start_ends:
            ax.hlines(0, start-margin, end+margin, **kwargs)

# %% [code]
def test_blot(data, mol):
    data_test = data[['FactorTest', 'FactorCondition', 'Experiment']].copy()
    data_test['Measurement'] = np.log(data[('Measurement', mol)])

    if data_test['FactorCondition'].nunique() > 1:
        result_anova, result_tukey = r_anova_and_tukey(
            'Measurement ~ FactorTest * FactorCondition + Experiment',
            ['FactorTest', 'FactorCondition', 'Experiment'],
            'FactorTest',
            data_test,
        )
    else:
        result_anova, result_tukey = r_anova_and_tukey(
            'Measurement ~ FactorTest + Experiment',
            ['FactorTest', 'Experiment'],
            'FactorTest',
            data_test,
        )
    
    p_anova = result_anova.loc['FactorTest']['Pr(>F)']
    ps_tukey = dict(result_tukey['p adj'])
    
    return p_anova, ps_tukey

# %% [code]
def plot_p_value(
    ax,
    p,
    pos_x1,
    pos_x2,
    height,
    yscale=1.33,
    thickness=0.05,
    group_line=None,
    always_plot_frame=True,
):
    s = p_stars(p)
    
    if not always_plot_frame and not s:
        return
        
    h0 = (1.0 + 0 * thickness + height) * yscale
    h1 = (1.0 + 1 * thickness + height) * yscale
    h2 = (1.0 + 2 * thickness + height) * yscale
    color = 'k' if s else color_ns

    ax.plot(
        [pos_x1, pos_x1, pos_x2, pos_x2],
        [h0, h1, h1, h0],
        color=color,
        lw=0.75,
        clip_on=False,
    )

    ax.annotate(
        s,
        ((pos_x1 + pos_x2) / 2, h2),
        ha='center',
        va='bottom',
        fontsize=5,
        annotation_clip=False,
    )

    if not group_line:
        return

    for pos in [pos_x1, pos_x2]:
        ax.plot(
            [pos - group_line / 2, pos + group_line / 2],
            [h0, h0],
            color=color,
            lw=0.75,
            clip_on=False,
        )

# %% [code]
def plot_tests(
    axs,
    mols,
    data,
    plot_p_value_kwargs_dict,
):
    if isinstance(data, list):
        datas = data
    else:
        datas = len(mols) * [data]

    if isinstance(plot_p_value_kwargs_dict, list):
        plot_p_value_kwargs_dicts = plot_p_value_kwargs_dict
    else:
        plot_p_value_kwargs_dicts = len(mols) * [plot_p_value_kwargs_dict]

    tests = [
        test_blot(data, mol)
        for data, mol in zip(datas, mols)
    ]
    
    ps_anova = [p for p, _ in tests]
    pss_tukey = [ps for _, ps in tests]
    ps_bh_anova = ss.false_discovery_control(ps_anova, method='bh')

    for ax, p_bh_anova, ps_tukey, plot_p_value_kwargs_dict in zip(axs, ps_bh_anova, pss_tukey, plot_p_value_kwargs_dicts):
        if p_bh_anova >= 0.05:
            ax.annotate(
                'ANOVA not significant',
                (0.5, 1.2),
                xycoords='axes fraction',
                va='center',
                ha='center',
                color=color_ns,
                style='italic',
            )
            continue

        for comparison, p in ps_tukey.items():
            kwargs = plot_p_value_kwargs_dict[comparison]
            plot_p_value(ax, p, **kwargs)

# %% [markdown]
# Figure S3A. IFNbeta & IFNlambda prestimulation + IAV

# %% [code]
data_dir = SOURCE_DATA_DIR / 'Western blot/Figure S3A'

for path_rep in sorted(data_dir.glob('rep*')):
    print(path_rep)

# %% [code]
data = pd.concat([
    blot_renorm_max(load_rep(path_rep))
    for path_rep in sorted(data_dir.glob('rep*'))
])

data = data.sort_values(by=['Experiment', 'Lane'])

# %% [code]
mols = [
    'p-STAT1', 'STAT1',
    'p-STAT2', 'STAT2',
    'RIG-I', 'PKR', 'OAS1',
    'IAV nucleoprotein',
]

# %% [code]
data_test = data.copy()
data_test = data_test[data_test[('Stimulation', 'Duration')] > '0 hour']

data_test['FactorTest'] = data_test[('Prestimulation', 'Agent')].fillna('mock')
data_test['FactorCondition'] = (
    data_test[('Stimulation', 'Amount')].apply(lambda m: f"MOI_{m}")
  + data_test[('Stimulation', 'Duration')].apply(lambda h: f"_{h / pd.Timedelta('1 hour'):.0f}h")
)

plot_p_value_kwargs_dict = {
    'IFNlambda-IFNbeta': dict(pos_x1=6.65, pos_x2=10.5, height=0.0),
    'mock-IFNlambda': dict(pos_x1=2.5, pos_x2=10.5, height=0.40),
    'mock-IFNbeta': dict(pos_x1=2.5, pos_x2=6.35, height=0.0),
}

# %% [code]
lanes = [
     2,  3,  4,  5,
     7,  8,  9, 10,
    12, 13, 14, 15,
]

lane_labels = [
    '24h',
    '48h',
    '24h',
    '48h',

    '24h',
    '48h',
    '24h',
    '48h',

    '24h',
    '48h',
    '24h',
    '48h',
]

exp_to_color = {
    'rep2': 'blue',
}

fig, axs = plot_blots(
    data[data['Lane'].isin(lanes)],
    mols=mols,
    lanes=lanes,
    lane_labels=lane_labels,
    exp_to_color=exp_to_color,
    hspace=0.35,
    scalex=0.9,
    scaley=0.8,
    top=0.15,
    bottom=0.45,
)

plot_groups(
    axs,
    [(1, 4), (5, 8), (9, 12)]
)

plot_tests(
    axs,
    mols,
    data=data_test,
    plot_p_value_kwargs_dict=plot_p_value_kwargs_dict,
)

# hide all tick labels but last
for ax in axs[:-1]:
    ax.tick_params(labelbottom=False)

# add custom 3-layer ticks
ax = axs[-1]
sec = ax.secondary_xaxis(location=0)
sec.set_xticks([1.5, 3.5, 5.5, 7.5, 9.5, 11.5], labels=3*['0.1', '1'])
sec.tick_params('x', length=0, pad=15)
thr = ax.secondary_xaxis(location=0)
thr.set_xticks([2.5, 6.5, 10.5], labels=['mock', 'IFN-β', 'IFN-λ1'])
thr.tick_params('x', length=0, pad=25)

ax.annotate('MOI', (-0.105, -1.4), xycoords='axes fraction', annotation_clip=False)
ax.annotate('Prestim.', (-0.105, -2.1), xycoords='axes fraction', annotation_clip=False)

fig.savefig('./Figures/Figure 2/figure_2b--IAV_IFNbeta_IFNlambda_prestim.svg', transparent=True)

# %% [markdown]
# Figure S3B. IFNbeta & IFNlambda prestimulation + RSV

# %% [code]
data_dir = SOURCE_DATA_DIR / 'Western blot/Figure S3B'

for path_rep in sorted(data_dir.glob('rep*')):
    print(path_rep)

# %% [code]
data = pd.concat([
    blot_renorm_max(load_rep(path_rep))
    for path_rep in sorted(data_dir.glob('rep*'))
])

data = data.sort_values(by=['Experiment', 'Lane'])

# %% [code]
mols = [
    'p-STAT1', 'STAT1',
    'p-STAT2', 'STAT2',
    'RIG-I', 'PKR', 'OAS1',
    'RSV F protein',
]

# %% [code]
data_test = data.copy()
data_test = data_test[data_test[('Stimulation', 'Duration')] > '0 hour']

data_test['FactorTest'] = data_test[('Prestimulation', 'Agent')].fillna('mock')
data_test['FactorCondition'] = (
    data_test[('Stimulation', 'Amount')].apply(lambda m: f"MOI_{m}")
  + data_test[('Stimulation', 'Duration')].apply(lambda h: f"_{h / pd.Timedelta('1 hour'):.0f}h")
)

plot_p_value_kwargs_dict = {
    'IFNlambda-IFNbeta': dict(pos_x1=6.65, pos_x2=10.5, height=0.0),
    'mock-IFNlambda': dict(pos_x1=2.5, pos_x2=10.5, height=0.40),
    'mock-IFNbeta': dict(pos_x1=2.5, pos_x2=6.35, height=0.0),
}

# %% [code]
lanes = [
     2,  3,  4,  5,
     7,  8,  9, 10,
    12, 13, 14, 15,
]

lane_labels = [
    '24h',
    '48h',
    '24h',
    '48h',

    '24h',
    '48h',
    '24h',
    '48h',

    '24h',
    '48h',
    '24h',
    '48h',
]

exp_to_color = {
    'rep1': 'blue',
}

fig, axs = plot_blots(
    data[data['Lane'].isin(lanes)],
    mols=mols,
    lanes=lanes,
    lane_labels=lane_labels,
    exp_to_color=exp_to_color,
    hspace=0.35,
    scalex=0.9,
    scaley=0.8,
    top=0.15,
    bottom=0.45,
)

plot_groups(
    axs,
    [(1, 4), (5, 8), (9, 12)]
)

plot_tests(
    axs,
    mols,
    data=data_test,
    plot_p_value_kwargs_dict=plot_p_value_kwargs_dict,
)

# hide all tick labels but last
for ax in axs[:-1]:
    ax.tick_params(labelbottom=False)

# add custom 3-layer ticks
ax = axs[-1]
sec = ax.secondary_xaxis(location=0)
sec.set_xticks([1.5, 3.5, 5.5, 7.5, 9.5, 11.5], labels=3*['0.1', '1'])
sec.tick_params('x', length=0, pad=15)
thr = ax.secondary_xaxis(location=0)
thr.set_xticks([2.5, 6.5, 10.5], labels=['mock', 'IFN-β', 'IFN-λ1'])
thr.tick_params('x', length=0, pad=25)

ax.annotate('MOI', (-0.105, -1.4), xycoords='axes fraction', annotation_clip=False)
ax.annotate('Prestim.', (-0.105, -2.1), xycoords='axes fraction', annotation_clip=False)

fig.savefig('./Figures/Figure 2/figure_2d--RSV_IFNbeta_IFNlambda_prestim.svg', transparent=True)

# %% [markdown]
# Figure 3B. IFN_R KOs + IAV

# %% [code]
data_dir = SOURCE_DATA_DIR / 'Western blot/Figure 3B'

for path_rep in sorted(data_dir.glob('rep*')):
    print(path_rep)

# %% [code]
data = pd.concat([
    blot_renorm_max(load_rep(path_rep))
    for path_rep in sorted(data_dir.glob('rep*'))
])

# %% [code]
mols = [
    'p-STAT1', 'STAT1',
    'p-STAT2', 'STAT2',
    'RIG-I', 'PKR', 'OAS1',
    'IAV nucleoprotein',
]

# %% [code]
data_test = data.copy()
data_test = data_test[data_test[('Stimulation', 'Duration')] > '0 hour']
data_test = data_test[data_test['CellLine'] != 'IFNAR-IFNLR-KO']

data_test['FactorTest'] = data_test['CellLine']
data_test['FactorCondition'] = data_test[('Stimulation', 'Duration')].apply(lambda h: f"{h / pd.Timedelta('1 hour'):.0f}h")

plot_p_value_kwargs_dict = {
    'IFNLR-KO-IFNAR-KO': dict(pos_x1=4.65, pos_x2=6.5, height=0.0),
    'WT-IFNLR-KO': dict(pos_x1=2.5, pos_x2=6.5, height=0.40),
    'WT-IFNAR-KO': dict(pos_x1=2.5, pos_x2=4.35, height=0.0),
}

# %% [code]
lanes = [1, 2, 3, 5, 6, 8, 9]

lane_labels = [
    'ctrl',
    '24h',
    '48h',
    '24h',
    '48h',
    '24h',
    '48h',
]

exp_to_color_short = {
    'rep5': 'blue',
}

mol_to_ylim_max = {}

fig, axs = plot_blots(
    data,
    mols=mols,
    lanes=lanes,
    lane_labels=lane_labels,
    exp_to_color=exp_to_color_short,
    mol_to_ylim_max=mol_to_ylim_max,
    hspace=0.55,
    scalex=1.5,
    scaley=0.8,
    scalebarwidth=0.4,
    scaleoffset=0.4,
    top=0.15,
)

plot_groups(
    axs,
    [(1, 1), (2, 3), (4, 5), (6, 7)],
    margin=0.33
)

plot_tests(
    axs,
    mols,
    data=data_test,
    plot_p_value_kwargs_dict=plot_p_value_kwargs_dict,
)

for ax in axs:
    sec = ax.secondary_xaxis(location=0)
    sec.set_xticks([1, 2.5, 4.5, 6.5], labels=['WT', 'WT', 'IFNAR1 KO', 'IFNLR1 KO'])
    sec.tick_params('x', length=0, pad=15)

fig.savefig('./Figures/Figure 3/figure_3b--IAV_IFNxR_KOs.svg', transparent=True)

# %% [markdown]
# Figure 3D. IFN_R KOs + RSV

# %% [code]
data_dir = SOURCE_DATA_DIR / 'Western blot/Figure 3D'

for path_rep in sorted(data_dir.glob('rep*')):
    print(path_rep)

# %% [code]
data = pd.concat([
    blot_renorm_max(load_rep(path_rep))
    for path_rep in sorted(data_dir.glob('rep*'))
])

# %% [code]
mols = [
    'p-STAT1', 'STAT1',
    'p-STAT2', 'STAT2',
    'RIG-I', 'PKR', 'OAS1',
    'RSV F protein',
]

# %% [code]
data_test = data.copy()
data_test = data_test[data_test[('Stimulation', 'Duration')] > '0 hour']
data_test = data_test[data_test['CellLine'] != 'IFNAR-IFNLR-KO']

data_test['FactorTest'] = data_test['CellLine']
data_test['FactorCondition'] = data_test[('Stimulation', 'Duration')].apply(lambda h: f"{h / pd.Timedelta('1 hour'):.0f}h")

data_tests = len(mols[:-1]) * [data_test] + [
    data_test[data_test[('Stimulation', 'Duration')] == '48 hour']
]

plot_p_value_kwargs_dicts = len(mols[:-1]) * [{
    'IFNLR-KO-IFNAR-KO': dict(pos_x1=4.65, pos_x2=6.5, height=0.0),
    'WT-IFNLR-KO': dict(pos_x1=2.5, pos_x2=6.5, height=0.40),
    'WT-IFNAR-KO': dict(pos_x1=2.5, pos_x2=4.35, height=0.0),
}] + [{
    'IFNLR-KO-IFNAR-KO': dict(pos_x1=5.1, pos_x2=7.0, height=0.0),
    'WT-IFNLR-KO': dict(pos_x1=3.0, pos_x2=7.0, height=0.40),
    'WT-IFNAR-KO': dict(pos_x1=3.0, pos_x2=4.9, height=0.0),
}]

# %% [code]
lanes = [1, 2, 3, 5, 6, 8, 9]

lane_labels = [
    'ctrl',
    '24h',
    '48h',
    '24h',
    '48h',
    '24h',
    '48h',
]

exp_to_color_short = {
    'rep3': 'blue',
}

fig, axs = plot_blots(
    data,
    mols=mols,
    lanes=lanes,
    lane_labels=lane_labels,
    exp_to_color=exp_to_color_short,
    mol_to_ylim_max=mol_to_ylim_max,
    hspace=0.55,
    scalex=1.5,
    scaley=0.8,
    scalebarwidth=0.4,
    scaleoffset=0.4,
    top=0.15,
)

plot_groups(
    axs,
    [(1, 1), (2, 3), (4, 5), (6, 7)],
    margin=0.33
)

plot_tests(
    axs,
    mols,
    data=data_tests,
    plot_p_value_kwargs_dict=plot_p_value_kwargs_dicts,
)

for ax in axs:
    sec = ax.secondary_xaxis(location=0)
    sec.set_xticks([1, 2.5, 4.5, 6.5], labels=['WT', 'WT', 'IFNAR1 KO', 'IFNLR1 KO'])
    sec.tick_params('x', length=0, pad=15)

fig.savefig('./Figures/Figure 3/figure_3d--RSV_IFNxR_KOs.svg', transparent=True)

# %% [markdown]
# Figure S1B. IFNbeta & IFNlambda prestimulation, no virus

# %% [code]
data_dir = SOURCE_DATA_DIR / 'Western blot/Figure S1B'

for path_rep in sorted(data_dir.glob('rep*')):
    print(path_rep)

# %% [code]
data = pd.concat([
    blot_renorm_max(load_rep(path_rep))
    for path_rep in sorted(data_dir.glob('rep*'))
])

data = data.sort_values(by=['Experiment', 'Lane'])

# %% [code]
data_test = data.copy()

data_test = data_test[~data_test[('Prestimulation', 'Agent')].isna()]
data_test['FactorTest'] = data_test[('Prestimulation', 'Agent')]
data_test['FactorCondition'] = data_test[('Prestimulation', 'Duration')].apply(lambda h: f"{h / pd.Timedelta('1 hour'):.0f}h")

data_test_24h = data_test[data_test[('Prestimulation', 'Duration')] >= '24 hour']
data_tests = [data_test, data_test_24h, data_test, data_test_24h, data_test_24h, data_test_24h, data_test_24h]

plot_p_value_kwargs_dicts = [
    {'IFNlambda-IFNbeta': dict(pos_x1=4, pos_x2=9, height=0.05, thickness=0.15, group_line=4.5)},
    {'IFNlambda-IFNbeta': dict(pos_x1=5.5, pos_x2=10.5, height=0.05, thickness=0.15, group_line=1.5)},
    {'IFNlambda-IFNbeta': dict(pos_x1=4, pos_x2=9, height=0.05, thickness=0.15, group_line=4.5)},
    {'IFNlambda-IFNbeta': dict(pos_x1=5.5, pos_x2=10.5, height=0.05, thickness=0.15, group_line=1.5)},
    {'IFNlambda-IFNbeta': dict(pos_x1=5.5, pos_x2=10.5, height=0.05, thickness=0.15, group_line=1.5)},
    {'IFNlambda-IFNbeta': dict(pos_x1=5.5, pos_x2=10.5, height=0.05, thickness=0.15, group_line=1.5)},
    {'IFNlambda-IFNbeta': dict(pos_x1=5.5, pos_x2=10.5, height=0.05, thickness=0.15, group_line=1.5)},
]

# %% [code]
experiments = data['Experiment'].unique()

mols = [
    'p-STAT1', 'STAT1',
    'p-STAT2', 'STAT2',
    'RIG-I', 'PKR', 'OAS1',
]

lane_labels = [
    'ctrl\nmock',
    
    '1h',
    '2h',
    '6h\nIFN-β',
    '24h',
    '48h',
    
    '1h',
    '2h',
    '6h\nIFN-λ1',
    '24h',
    '48h',
]

exp_to_color = {
    'rep1': 'blue',
}

fig, axs = plot_blots(
    data,
    mols=mols,
    lane_labels=lane_labels,
    exp_to_color=exp_to_color,
    hspace=0.5,
    top=0.15,
)

plot_groups(
    axs,
    [(1, 1), (2, 6), (7, 11)]
)

plot_tests(
    axs,
    mols,
    data=data_tests,
    plot_p_value_kwargs_dict=plot_p_value_kwargs_dicts,
)

fig.savefig('./Figures/Figure S1/figure_s1b--IFNbeta_IFNlambda_prestim.svg', transparent=True)

# %% [markdown]
# Figure S2C. IFN-λ saturation dose

# %% [code]
data_dir = SOURCE_DATA_DIR / 'Western blot/Figure S2C'

for path_rep in sorted(data_dir.glob('rep*')):
    print(path_rep)

# %% [code]
data = pd.concat([
    load_rep(path_rep)
    for path_rep in sorted(data_dir.glob('rep*'))[::-1]
])

# %% [code]
lane_labels = [
    '0',
    '0.05',
    '0.1',
    '0.5',
    '1',
    '2.5',
    '5',
    '10',
    '25',
    '50',
    '100',
]
lane_labels = [f"{l} ng/ml" for l in lane_labels]

exp_to_color = {
    'rep2': 'blue',
}

mols = [
    'p-STAT1',
    'p-STAT2',
]

fig, axs = plot_blots(data, mols=mols, lane_labels=lane_labels, exp_to_color=exp_to_color, scalex=1.75, top=0.1)
fig.savefig('./Figures/Figure S2/figure_s2c--IFNlambda_doses.svg', transparent=True)

# %% [markdown]
# Figure S2D. IFN-β saturation dose

# %% [code]
data_dir = SOURCE_DATA_DIR / 'Western blot/Figure S2D'

for path_rep in sorted(data_dir.glob('rep*')):
    print(path_rep)

# %% [code]
data = pd.concat([
    load_rep(path_rep)
    for path_rep in sorted(data_dir.glob('rep*'))
])

# %% [code]
lane_labels = [
    '0',
    '30',
    '100',
    '300',
    '1000',
    '3000',
    '10000',
]
lane_labels = [f"{l} U/ml" for l in lane_labels]

exp_to_color = {
    'rep1': 'blue',
}

mols = [
    'p-STAT1',
    'p-STAT2',
]

fig, axs = plot_blots(data, mols=mols, lane_labels=lane_labels, exp_to_color=exp_to_color, scalex=1.75, top=0.1)
fig.savefig('./Figures/Figure S2/figure_s2d--IFNbeta_doses.svg', transparent=True)

# %% [markdown]
# Figure S4. IFN_R KOs + IAV

# %% [code]
data_dir = SOURCE_DATA_DIR / 'Western blot/Figure S4'

for path_rep in sorted(data_dir.glob('rep*')):
    print(path_rep)

# %% [code]
data = pd.concat([
    blot_renorm_max(load_rep(path_rep))
    for path_rep in sorted(data_dir.glob('rep*'))
])

# %% [code]
mols = [
    'p-STAT1', 'STAT1',
    'p-STAT2', 'STAT2',
    'RIG-I', 'PKR', 'OAS1',
    'IAV nucleoprotein',
]

lane_labels = [
    'ctrl',
    '24h\nWT',
    '48h',
    'ctrl',
    '24h\nIFNAR1 KO',
    '48h',
    'ctrl',
    '24h\nIFNLR1 KO',
    '48h',
    'ctrl',
    '24h\nDKO',
    '48h',
]

exp_to_color = {
    exp: f"C{i}"
    for i, exp in enumerate(data['Experiment'].unique())
}

fig, axs = plot_blots(
    data,
    mols=mols,
    lane_labels=lane_labels,
    exp_to_color=exp_to_color,
    top=0.15,
    hspace=0.4,
    scaley=0.8,
    scalebarwidth=0.6,
    scaleoffset=0.6,
)

plot_groups(
    axs,
    [(1, 3), (4, 6), (7, 9), (10, 12)],
    margin=0.33,
)

fig.savefig('./Figures/Figure S3/figure-s3.svg', transparent=True)

# %% [markdown]
# Figure S5. IFN_R KOs + RSV

# %% [code]
data_dir = SOURCE_DATA_DIR / 'Western blot/Figure S5'

for path_rep in sorted(data_dir.glob('rep*')):
    print(path_rep)

# %% [code]
data = pd.concat([
    blot_renorm_max(load_rep(path_rep))
    for path_rep in sorted(data_dir.glob('rep*'))
])

# %% [code]
mols = [
    'p-STAT1', 'STAT1',
    'p-STAT2', 'STAT2',
    'RIG-I', 'PKR', 'OAS1',
    'RSV F protein',
]

lane_labels = [
    'ctrl',
    '24h\nWT',
    '48h',
    'ctrl',
    '24h\nIFNAR1 KO',
    '48h',
    'ctrl',
    '24h\nIFNLR1 KO',
    '48h',
    'ctrl',
    '24h\nDKO',
    '48h',
]

exp_to_color = {
    exp: f"C{i}"
    for i, exp in enumerate(data['Experiment'].unique())
}

fig, axs = plot_blots(
    data,
    mols=mols,
    lane_labels=lane_labels,
    exp_to_color=exp_to_color,
    top=0.15,
    hspace=0.4,
    scaley=0.8,
    scalebarwidth=0.6,
    scaleoffset=0.6,
)

plot_groups(
    axs,
    [(1, 3), (4, 6), (7, 9), (10, 12)],
    margin=0.33,
)
fig.savefig('./Figures/Figure S4/figure-s4.svg')
```
