## Supplementary Data S2 for "Type III interferons may suppress viral infections by triggering cell death": Figure 4 and S6 plot dPCR, immunostaining.ipynb

```
# %% [code]
from pathlib import Path

import numpy as np
import pandas as pd
import scipy.stats as ss

import matplotlib.pyplot as plt
from matplotlib.lines import Line2D

from subplots_from_axsize import subplots_from_axsize

from utils import p_stars, time_norm, time_cmap, time_cmap_normed, moi_to_marker

# %% [code]
SOURCE_DATA_DIR = Path('./Quantifications')

# %% [code]
!mkdir -p "Figures/Figure 4"
!mkdir -p "Figures/Figure S6"

# %% [markdown]
# plot utils

# %% [code]
cell_line_names_default = {
    'WT': 'WT',
    'IFNAR-KO': 'IFNAR1 KO',
    'IFNLR-KO': 'IFNLR1 KO',
}

def plot_cols(
    data,
    cell_line_names=cell_line_names_default,
    yticks=[1/3, 1, 3, 10],
    yticklabels=['⅓', '1', '3', '10'],
    **subplots_kwargs,
):
    log_yticks = np.log10(yticks)
    cell_lines = list(data.columns)
    data = data.reset_index()

    axx = len(cell_lines) + 2

    subplots_kwargs = dict(
        axsize=(axx * 0.55, 2),
        left=0.65,
        right=0.15,
        bottom=0.5,
        top=0.5,
    ) | subplots_kwargs
    
    fig, ax = subplots_from_axsize(**subplots_kwargs)

    cell_lines_positions = list(range(1, len(cell_lines) + 1))

    # scatters
    for moi, data_moi in data.groupby("('Stimulation', 'Amount')"):
        for pos, cell_line in zip(cell_lines_positions, cell_lines):
            data_plot = data_moi[["('Stimulation', 'Duration')", cell_line]].dropna()
            ratios = data_plot[cell_line]
            days = data_plot["('Stimulation', 'Duration')"] / pd.Timedelta(days=1)
            ax.scatter(
                pos + np.random.uniform(-0.1, 0.1, size=(len(ratios),)),
                np.log10(ratios),
                marker=moi_to_marker[moi],
                c=days,
                norm=time_norm,
                cmap=time_cmap,
                alpha=0.7,
                clip_on=False,
            )

    # errors and p-stars
    for pos, cell_line in zip(cell_lines_positions, cell_lines):
        ratios = data[cell_line].dropna()
        log_ratios = np.log10(ratios)

        ax.errorbar(
            pos, log_ratios.mean(),
            yerr=(log_ratios.std() / len(log_ratios)**0.5),
            color='k',
            capsize=2,
            zorder=20,
        )

        p = ss.ttest_1samp(log_ratios, popmean=0).pvalue
        s = p_stars(p)
        s = s if s else 'n.s.'
        ax.annotate(
            s,
            (pos, log_yticks.max() + 0.15 * (log_yticks.max() - log_yticks.min())),
            ha='center',
            va='center',
            annotation_clip=False,
        )

    ax.spines[['bottom', 'top', 'right']].set_visible(False)
    
    ax.set_xlim(0.25, 0.50 + len(cell_lines))
    ax.set_xticks(cell_lines_positions)
    ax.set_xticklabels([cell_line_names[cell_line] for cell_line in cell_lines])

    ax.set_ylim(log_yticks.min(), log_yticks.max())
    ax.set_yticks(np.log10(yticks))
    ax.set_yticklabels(yticklabels)

    ax.axhline(0, color='gray', ls=':')

    return fig, ax

# %% [markdown]
# Figure 4A

# %% [code]
data = pd.read_csv(SOURCE_DATA_DIR / 'Imaging/Figure 4A/all_reps.csv')

data["('Stimulation', 'Duration')"] = pd.to_timedelta(data["('Stimulation', 'Duration')"])

data = data[data["('Stimulation', 'Agent')"] == 'IAV']
del data["('Stimulation', 'Agent')"]
del data["('Stimulation', 'Unit')"]

data['FractionVirusPositive'] = data['VirusPositive'] / data['Total']

col = 'FractionVirusPositive'
data = data.groupby(['Experiment', 'CellLine'] + [c for c in data if 'Stimulation' in c])[[col]].mean()
data = data.reset_index('CellLine').pivot(columns='CellLine', values=col)

data['IFNAR-KO'] = data['IFNAR-KO'] / data['WT']
data['IFNLR-KO'] = data['IFNLR-KO'] / data['WT']
del data['WT']

data

# %% [code]
fig, ax = plot_cols(data, right=1.3)

ax.set_ylabel('Ratio of the fraction of IAV-positive cells\nbetween KO and WT')

ax.legend(
    [
        Line2D([0], [0], marker='^', ls='none', color=time_cmap_normed(1), alpha=0.5),
        Line2D([0], [0], marker='^', ls='none', color=time_cmap_normed(2), alpha=0.5),
        Line2D([0], [0], marker='^', ls='none', color=time_cmap_normed(3), alpha=0.5),
        Line2D([0], [0], marker='^', ls='none', color=time_cmap_normed(4), alpha=0.5),
        Line2D([0], [0], marker='none', ls='none', color='k', alpha=0.5),
        Line2D([0], [0], marker=moi_to_marker[0.01], ls='none', color='k', alpha=0.5),
        Line2D([0], [0], marker=moi_to_marker[0.1], ls='none', color='k', alpha=0.5),
    ],
    ['24 h p.i.', '48 h p.i.', '72 h p.i.', '96 h p.i.', '', 'MOI 0.01', 'MOI 0.1'],
    bbox_to_anchor=(1.03, 0.95),
    columnspacing=0.4,
    handletextpad=0.2,
    loc='upper left',
    
)

fig.savefig('./Figures/Figure 4/figure_4a--IFNxR_KO.svg')

# %% [markdown]
# Figure 4B

# %% [code]
data = pd.read_csv(SOURCE_DATA_DIR / 'Imaging/Figure 4B/all_reps.csv')

data["('Stimulation', 'Duration')"] = pd.to_timedelta(data["('Stimulation', 'Duration')"])

del data["('Prestimulation', 'Agent')"]
del data["('Prestimulation', 'Amount')"]
del data["('Prestimulation', 'Unit')"]
del data["('Prestimulation', 'Duration')"]
del data["('Stimulation', 'Agent')"]
del data["('Stimulation', 'Unit')"]

data['FractionVirusPositive'] = data['VirusPositive'] / data['Total']

col = 'FractionVirusPositive'
data = data.groupby(['Experiment', 'CellLine'] + [c for c in data if 'Stimulation' in c])[[col]].mean()
data = data.reset_index('CellLine').pivot(columns='CellLine', values=col)

data['IFNLR-KO'] = data['IFNLR-KO'] / data['WT']
del data['WT']

data

# %% [code]
fig, ax = plot_cols(data)

ax.set_ylabel('Ratio of the fraction of IAV-positive cells\nbetween KO and WT')

fig.savefig('./Figures/Figure 4/figure_4b--IFNbeta_IFNLR_KO.svg')

# %% [markdown]
# Figure S6

# %% [code]
data = pd.read_csv(SOURCE_DATA_DIR / 'dPCR/Figure S6/all_reps.csv')

data["('Stimulation', 'Duration')"] = pd.to_timedelta(data["('Stimulation', 'Duration')"])

del data["('Stimulation', 'Agent')"]
del data["('Stimulation', 'Unit')"]

col = 'RNACountPerCell_Mean'
data = data.groupby(['Experiment', 'CellLine'] + [c for c in data if 'Stimulation' in c])[[col]].mean()
data = data.reset_index('CellLine').pivot(columns='CellLine', values=col)

data['IFNLR-KO'] = data['IFNLR-KO'] / data['WT']
del data['WT']

data

# %% [code]
fig, ax = plot_cols(data, right=1.3)

ax.set_ylabel('Ratio of IAV RNA molecule counts\nbetween IFNLR1 KO and WT')

ax.legend(
    [
        Line2D([0], [0], marker='^', ls='none', color=time_cmap_normed(1), alpha=0.5),
        Line2D([0], [0], marker='^', ls='none', color=time_cmap_normed(2), alpha=0.5),
        Line2D([0], [0], marker='^', ls='none', color=time_cmap_normed(3), alpha=0.5),
        Line2D([0], [0], marker='^', ls='none', color=time_cmap_normed(4), alpha=0.5),
        Line2D([0], [0], marker='none', ls='none', color='k', alpha=0.5),
        Line2D([0], [0], marker=moi_to_marker[0.01], ls='none', color='k', alpha=0.5),
        Line2D([0], [0], marker=moi_to_marker[0.1], ls='none', color='k', alpha=0.5),
    ],
    ['24 h p.i.', '48 h p.i.', '72 h p.i.', '96 h p.i.', '', 'MOI 0.01', 'MOI 0.1'],
    bbox_to_anchor=(1.05, 0.95),
    columnspacing=0.4,
    handletextpad=0.2,
    loc='upper left',
)

fig.savefig('./Figures/Figure S5/figure-s5.svg')

# %% [code]
```
