## Supplementary Data S2 for "Type III interferons may suppress viral infections by triggering cell death": Figure 5 and S8 plot SIR.ipynb

```
# %% [code]
import numpy as np
import scipy

import matplotlib as mpl
import matplotlib.pyplot as plt
import matplotlib.colors as mcolors
import matplotlib.lines as mlines

from subplots_from_axsize import subplots_from_axsize

import utils  # font settings

# %% [code]
!mkdir -p "Figures/Figure 5"
!mkdir -p "Figures/Figure S8"

# %% [code]
plt.rcParams['svg.fonttype'] = 'none'

# %% [code]
def sir_grad(y, t, beta, lmbda):
    ys, yi, yr = y
    
    return np.stack([
        -yi * ys * beta,
        yi * ys * beta - lmbda * yi,
        lmbda * yi,
    ])


def sir_solve(moi, ts, beta, lmbda):
    y0 = np.array([1-moi, moi, 0.0])
    result = scipy.integrate.odeint(
        sir_grad,
        y0,
        ts,
        args=(beta, lmbda),
    )
    return result

# %% [code]
def plot_by_beta(ax, ts, xs_by_betas, betas, ylabel, annotate_higher=False):
    assert xs_by_betas.shape[0] == len(betas)
    assert xs_by_betas.shape[1] == len(ts)

    cmap_beta = lambda c: mpl.colormaps['winter'](mcolors.LogNorm(vmin=betas.min(), vmax=betas.max())(c))

    for xs, beta in zip(xs_by_betas, betas):
        ax.plot(ts, xs, color=cmap_beta(beta))

    h = 1.1 if annotate_higher else 1.0
    ax.annotate('β', (0.39, h), xycoords='axes fraction', va='center', ha='right', color='darkblue', fontsize=12)
    ax.annotate('  varied', (0.53, h), xycoords='axes fraction', va='center', ha='center', fontsize=12)
    ax.set_ylabel(ylabel, fontsize=12)


def plot_by_lmbda(ax, ts, xs_by_lmbdas, lmbdas, ylabel, annotate_higher=False):
    assert xs_by_lmbdas.shape[0] == len(lmbdas)
    assert xs_by_lmbdas.shape[1] == len(ts)

    cmap_lmbda = lambda c: mpl.colormaps['autumn'](mcolors.LogNorm(vmin=lmbdas.min(), vmax=lmbdas.max()*1.3)(c))

    for xs, lmbda in zip(xs_by_lmbdas, lmbdas):
        ax.plot(ts, xs, color=cmap_lmbda(lmbda))

    h = 1.1 if annotate_higher else 1.0
    ax.annotate('λ', (0.39, h), xycoords='axes fraction', va='center', ha='right', color='darkred', fontsize=12)
    ax.annotate('  varied', (0.53, h), xycoords='axes fraction', va='center', ha='center', fontsize=12)
    ax.set_ylabel(ylabel, fontsize=12)


def make_sir_figure(moi, ts, beta_default, lmbda_default, legend=True):
    changes = 10**np.linspace(-0.5, 0.5, 5)
    lmbdas = lmbda_default * changes
    betas = beta_default * changes

    yss_by_lmbdas = np.stack([sir_solve(moi=moi, ts=ts, beta=beta_default, lmbda=lmbda) for lmbda in lmbdas])
    yss_by_betas = np.stack([sir_solve(moi=moi, ts=ts, beta=beta, lmbda=lmbda_default) for beta in betas])
    yis_by_lmbdas, yrs_by_lmbdas = yss_by_lmbdas[..., 1], yss_by_lmbdas[..., 2]
    yis_by_betas, yrs_by_betas = yss_by_betas[..., 1], yss_by_betas[..., 2]

    cmap_beta = lambda c: mpl.colormaps['winter'](mcolors.LogNorm(vmin=betas.min(), vmax=betas.max())(c))
    cmap_lmbda = lambda c: mpl.colormaps['autumn'](mcolors.LogNorm(vmin=lmbdas.min(), vmax=lmbdas.max()*1.3)(c))

    fig, axs = subplots_from_axsize(
        ncols=3, nrows=2,
        axsize=(2, 1.5),
        bottom=2.0 if legend else 0.4,
        hspace=0.85,
        wspace=0.90,
    )

    plot_by_lmbda(axs[0, 0], ts, yrs_by_lmbdas, lmbdas, ylabel='R')
    plot_by_lmbda(axs[0, 1], ts, yis_by_lmbdas, lmbdas, ylabel='I')
    plot_by_beta(axs[0, 2], ts, yis_by_betas, betas, ylabel='I')
    plot_by_lmbda(axs[1, 0], ts, yrs_by_lmbdas / yis_by_lmbdas, lmbdas, ylabel='R / I', annotate_higher=True)
    plot_by_beta(axs[1, 1], ts, yrs_by_betas / yis_by_betas, lmbdas, ylabel='R / I', annotate_higher=True)
    plot_by_lmbda(axs[1, 2], ts, yrs_by_lmbdas / (yrs_by_lmbdas + yis_by_lmbdas), lmbdas, ylabel='R / (R + I)', annotate_higher=True)
    
    for ax in axs.ravel():
        #ax.set_xlabel(f'time [initial doubling time for β={float(beta_default):.2f}]')
        ax.set_xlabel(f'Time')
        ax.set_ylim(0.0, 1.0)
        ax.spines['right'].set_visible(False)
        ax.spines['top'].set_visible(False)
        
    for ax in axs[1, :2]:
        ax.set_ylim(0.07, 10)
        ax.set_yscale('log')
        ax.set_yticks([0.1, 1, 10])
        ax.set_yticklabels([0.1, 1, 10])
        ax.axhline(1, color='grey', ls=':', lw=1)

    if legend:
        axs[1, 0].legend(
            [mlines.Line2D([0], [0], color=cmap_lmbda(lmbda)) for lmbda in lmbdas]
          + [mlines.Line2D([0], [0], color=cmap_beta(beta)) for beta in betas],
            [f'λ={float(lmbda):.2f}' for lmbda in lmbdas]
          + [f'β={float(beta):.2f}' for beta in betas],
            ncol=2,
            title='β, λ values (default bold)',
            bbox_to_anchor=(-0.04, -0.3),
            loc='upper left',
        )
    
    return fig, axs

# %% [markdown]
# plot

# %% [code]
beta_default = np.log(2)
lmbda_default = 0.3

# %% [markdown]
# Figure 5

# %% [code]
fig, axs = make_sir_figure(
    moi=0.1,
    ts=np.linspace(0.0, 12.0, 201),
    beta_default=beta_default,
    lmbda_default=lmbda_default,
)

fig.savefig('./Figures/Figure 5/figure_5x--SIR.svg', transparent=True)

# %% [markdown]
# Figure S8A

# %% [code]
fig, axs = make_sir_figure(
    moi=0.01,
    ts=np.linspace(0.0, 30.0, 201),
    beta_default=beta_default,
    lmbda_default=lmbda_default,
    legend=False,
)

fig.savefig('./Figures/Figure S8/figure_s8a--MOI-0,01.svg', transparent=True)

# %% [markdown]
# Figure S8B

# %% [code]
fig, axs = make_sir_figure(
    moi=0.3,
    ts=np.linspace(0.0, 30.0, 201),
    beta_default=beta_default,
    lmbda_default=lmbda_default,
)

fig.savefig('./Figures/Figure S8/figure_s8b--MOI-0,3.svg', transparent=True)

# %% [code]
```
