## Supplementary Data S2 for "Type III interferons may suppress viral infections by triggering cell death": Figure 6 plot WB+FACS, immunostainig+live exp.ipynb

```
# %% [code]
from pathlib import Path

import numpy as np
import pandas as pd
import scipy.stats as ss

import matplotlib.pyplot as plt
import matplotlib.colors as mcolors
from matplotlib.lines import Line2D

from subplots_from_axsize import subplots_from_axsize

from utils import load_rep, time_norm, time_cmap, time_cmap_normed, moi_to_marker, p_stars

# %% [code]
SOURCE_DATA_DIR = Path('./Quantifications')

# %% [code]
!mkdir -p "Figures/Figure 6"

# %% [markdown]
# plot utils

# %% [code]
def prepare_plot(
    ylabel,
    xticklabels,
    yticks=[1/3, 1, 3, 10],
    yticklabels=['⅓', '1', '3', '10'],
    **kwargs,
):
    fig, ax = subplots_from_axsize(
        **(dict(
            axsize=(2.5, 3),
            left=0.75,
            right=0.75,
            bottom=0.4,
        ) | kwargs),
    )
    
    log_yticks = np.log10(yticks)
    
    ax.spines[['bottom', 'top', 'right']].set_visible(False)
    
    ax.set_xlim(-0.5, 1.5)
    ax.set_xticks([0.0, 1.0])
    ax.set_xticklabels(xticklabels)
    
    ax.set_ylim(log_yticks.min(), log_yticks.max())
    
    ax.set_ylabel(ylabel)
    
    ax.set_yticks(np.log10(yticks))
    ax.set_yticks([], minor=True)
    ax.set_yticklabels(yticklabels)
    
    ax.axhline(0.0, color='gray', ls=':', zorder=-20)

    return fig, ax

# %% [markdown]
# Figure 6A

# %% [markdown]
## prepare FACS

# %% [code]
data_dir = SOURCE_DATA_DIR / 'FACS/Figure 6A'

for path_rep in data_dir.glob('rep*'):
    print(path_rep)

# %% [code]
data = pd.concat([
    load_rep(path_rep)
    for path_rep in data_dir.glob('rep*')
])

data = data[[
    'Experiment',
    'CellLine',
    ('Stimulation', 'Amount'),
    ('Stimulation', 'Duration'),
    'State',
    '% Gated',
]]

data = data[data[('Stimulation', 'Amount')].isin([0.1])]

data = data[data['State'].isin(['late apoptosis', 'dead'])]

data = (
    data
    .groupby([
        'Experiment',
        'CellLine',
        ('Stimulation', 'Amount'),
        ('Stimulation', 'Duration'),
    ])
    ['% Gated']
    .sum()
    .to_frame('FractionDying')
)

data = data.reset_index('CellLine').pivot(columns='CellLine', values='FractionDying')

data['IFNLR-KO / WT'] = data['IFNLR-KO'] / data['WT']

data_facs = data.copy().reset_index()
data_facs

# %% [markdown]
## prepare Western blots

# %% [code]
data_dir_1 = SOURCE_DATA_DIR / 'Western blot/Figure 6A'
data_dir_2 = SOURCE_DATA_DIR / 'Western blot/Figure 3B'

for data_dir in (data_dir_1, data_dir_2):
    for path_rep in data_dir.glob('rep*'):
        print(path_rep)

# %% [code]
data = pd.concat([
    load_rep(path_rep).assign(Figure=path_rep.parent.name)  # used to rename exp to avoid rep ID clash
    for data_dir in (data_dir_1, data_dir_2)
    for path_rep in data_dir.glob('rep*')
])

data['Experiment'] = data['Figure'].apply(lambda f: f.split()[-1]) + '-' + data['Experiment']
data = data[data['CellLine'].isin(['WT', 'IFNLR-KO'])]
data = data[data[('Stimulation', 'Agent')] == 'IAV']
data = data[data[('Stimulation', 'Duration')] > '0 hours']

data['IAV nucleoprotein'] = data[('Measurement', 'IAV nucleoprotein')]
data = data[[
        'Experiment',
        'CellLine',
        ('Stimulation', 'Amount'),
        ('Stimulation', 'Duration'),
        'IAV nucleoprotein',
]]

data = data.set_index([
    'Experiment',
    ('Stimulation', 'Amount'),
    ('Stimulation', 'Duration'),
])

data = data.pivot(columns='CellLine', values='IAV nucleoprotein')

data['IFNLR-KO / WT'] = data['IFNLR-KO'] / data['WT']

data_blot = data.copy().reset_index()
data_blot

# %% [code]
fig, ax = prepare_plot(
    'Ratio between IFNLR1 KO and WT',
    ['IAV NP level', 'Fraction of\ndead cells'],
    yticks=[1/3, 1, 3, 10, 30],
    yticklabels=['⅓', '1', '3', '10', '30'],
    right=1.4,
)

# scatter
for pos, data in enumerate([data_blot, data_facs]):
    for moi, data_plot in data.groupby(('Stimulation', 'Amount')):
        log_ratios = np.log10(data_plot['IFNLR-KO / WT'])
        days = data_plot[('Stimulation', 'Duration')] / pd.Timedelta(days=1)
        
        ax.scatter(
            pos + np.random.uniform(-0.1, 0.1, size=(len(log_ratios),)),
            log_ratios,
            marker=moi_to_marker[moi],
            s=50,
            c=days,
            edgecolors='none',
            norm=time_norm,
            cmap=time_cmap,
            alpha=0.7,
            clip_on=False,
        )

# errors
for pos, data in enumerate([data_blot, data_facs]):
    log_ratios = np.log10(data['IFNLR-KO / WT'])

    ax.errorbar(
        pos, log_ratios.mean(),
        yerr=(log_ratios.std() / len(log_ratios)**0.5),
        color='k',
        capsize=2,
        zorder=20,
    )

# tests
log_ratios_left = np.log10(data_blot['IFNLR-KO / WT'])
log_ratios_right = np.log10(data_facs['IFNLR-KO / WT'])

res = ss.ttest_ind(log_ratios_left, log_ratios_right, equal_var=False)
p = res.pvalue
s = p_stars(p)
ax.annotate(f'unpaired Student $t$-test $p$={p:.5f} ({s})', (0.55, 0.92), xycoords='axes fraction', ha='center', clip_on=False)

res = ss.mannwhitneyu(log_ratios_left, log_ratios_right)
p = res.pvalue
s = p_stars(p)
ax.annotate(f'Mann-Whitney $U$ test $p$={p:.4f} ({s})', (0.55, 0.87), xycoords='axes fraction', ha='center', clip_on=False)

# comparison bracket
h = np.log10(13.5)
ax.plot(
    [0, 0, 1, 1],
    [0.98 * h, h, h, 0.98 * h],
    color='k',
    clip_on=False,
)

# legend
ax.legend(
    [
        Line2D([0], [0], marker='^', ls='none', color=time_cmap_normed(1), alpha=0.5),
        Line2D([0], [0], marker='^', ls='none', color=time_cmap_normed(2), alpha=0.5),

    ],
    ['24 h p.i.', '48 h p.i.'],
    bbox_to_anchor=(1.15, 1.05),
    columnspacing=0.4,
    #ncols=2,
    handletextpad=0.2,
    #loc='center right', 
)

fig.savefig('./Figures/Figure 6/figure_6a--blot+FACS_WT_vs_IFNLR_KO.svg')

# %% [markdown]
# Figure 6B

# %% [code]
data = load_rep(SOURCE_DATA_DIR / 'Imaging/Figure 6B/all_reps.csv')

data = data[data['CellLine'].isin(['WT', 'IFNLR-KO'])]
data = data[data[('Stimulation', 'Agent')].isin(['IAV', 'RSV'])]
data = data[data[('Stimulation', 'Duration')] > '0 hours']

assert set(data['Stimulation', 'Amount'].unique()) == {0.1}

data = (
    data
    .groupby([
        'Experiment',
        'CellLine',
        ('Stimulation', 'Agent'),
        ('Stimulation', 'Duration'),
    ])
    [['FractionVirusPositive', 'CumulatedDeathsFraction']]
    .mean()
)

data = data.reset_index(level=2).pivot(columns=[('Stimulation', 'Agent')])
data = data.reset_index()

data['FractionVirusPositive RSV / IAV'] = data['FractionVirusPositive', 'RSV'] / data['FractionVirusPositive', 'IAV']
data['CumulatedDeathsFraction RSV / IAV'] = data['CumulatedDeathsFraction', 'RSV'] / data['CumulatedDeathsFraction', 'IAV']

#del data['FractionVirusPositive']
#del data['CumulatedDeathsFraction']

data = data.dropna()

data

# %% [code]
fig, ax = prepare_plot(
    'Ratio between RSV and IAV infection',
    ['Fraction of\nvirus-positive cells', 'Fraction of\ndead cells'],
    yticks=[0.1, 1, 10, 100, 1000],
    yticklabels=['0.1', '1', '10', '100', '1000'],
    right=1.6,
)

cell_line_to_marker = {
    'WT': 'o',
    'IFNLR-KO': '^',
}

# lines (with markers)
for _, data_row in data.iterrows():
    ratios = list(data_row[['FractionVirusPositive RSV / IAV', 'CumulatedDeathsFraction RSV / IAV']])
    log_ratios = np.log10(ratios)
    days = data_row['Stimulation', 'Duration'] / pd.Timedelta(days=1)
    
    ax.plot(
        [0, 1],
        log_ratios,
        marker=cell_line_to_marker[data_row['CellLine'].iloc[0]],
        markersize=50**0.5,
        c=time_cmap(time_norm(days)),
        markeredgecolor='none',
        alpha=0.7,
        clip_on=False,
    )

# errors
for pos, col in enumerate(['FractionVirusPositive RSV / IAV', 'CumulatedDeathsFraction RSV / IAV']):
    log_ratios = np.log10(data[col])

    ax.errorbar(
        pos, log_ratios.mean(),
        yerr=(log_ratios.std() / len(log_ratios)**0.5),
        color='k',
        capsize=2,
        zorder=20,
    )

# tests
log_ratios_left = np.log10(data['FractionVirusPositive RSV / IAV'])
log_ratios_right = np.log10(data['CumulatedDeathsFraction RSV / IAV'])

res = ss.ttest_rel(log_ratios_left, log_ratios_right)
p = res.pvalue
s = p_stars(p)
ax.annotate(f'Student $t$-test $p$={p:.5f} ({s})', (0.55, 0.92), xycoords='axes fraction', ha='center', clip_on=False)

res = ss.wilcoxon(log_ratios_left, log_ratios_right)
p = res.pvalue
s = p_stars(p)
ax.annotate(f'Wilcoxon signed-rank test $p$={p:.4f} ({s})', (0.55, 0.87), xycoords='axes fraction', ha='center', clip_on=False)

# comparison bracket
h = np.log10(200)
ax.plot(
    [0, 0, 1, 1],
    [0.98 * h, h, h, 0.98 * h],
    color='k',
    clip_on=False,
)

# legend
ax.legend(
    [
        Line2D([0], [0], marker=cell_line_to_marker['WT'], ls='none', color='k', alpha=0.5),
        Line2D([0], [0], marker=cell_line_to_marker['IFNLR-KO'], ls='none', color='k', alpha=0.5),
        Line2D([0], [0], marker='^', ls='none', color=time_cmap_normed(1), alpha=0.5),
        Line2D([0], [0], marker='^', ls='none', color=time_cmap_normed(2), alpha=0.5),

    ],
    ['WT', 'IFNLR1 KO', '24 h p.i.', '48 h p.i.'],
    bbox_to_anchor=(1.15, 1.05),
    columnspacing=0.4,
    handletextpad=0.2,
    loc='upper left',
)

fig.savefig('./Figures/Figure 6/figure_6b--immuno+live_IAV_vs_RSV.svg')

# %% [code]
```
