## Supplementary Data S2 for "Type III interferons may suppress viral infections by triggering cell death": Figure 7C and 7D histograms live exp.ipynb

```
# %% [code]
from pathlib import Path
import matplotlib.pyplot as plt
import pandas as pd

from scipy.spatial import cKDTree

# %% [code]
SOURCE_DATA_DIR = Path('./Quantifications')

# %% [code]
!mkdir -p "Figures/Figure 7"

# %% [code]
def compare_intensity_by_celltype(h2b_data, pi_data):
    
    h2b_coords = h2b_data[['center_x', 'center_y']].to_numpy()
    pi_coords = pi_data[['center_x', 'center_y']].to_numpy()

    h2b_tree = cKDTree(h2b_coords)
    pi_tree = cKDTree(pi_coords)

    threshold = 5.0  

    distances, _ = pi_tree.query(h2b_coords, k=1)

    h2b_data = h2b_data[distances > threshold]

    fig, (ax1, ax2) = plt.subplots(2, 1, figsize=(10, 12))
    
    pi_color = 'red'
    h2b_color = 'blue'
    alpha = 0.6
    
    unique_celltypes = pi_data['Cell Type'].unique()
    celltype1 = unique_celltypes[0]
    pi_data_type1 = pi_data[pi_data['Cell Type'] == celltype1]
    h2b_data_type1 = h2b_data[h2b_data['Cell Type'] == celltype1]
    
    if 'cellevent_intensity_mean' in pi_data_type1.columns:
        pi_intensities_1 = pi_data_type1['cellevent_intensity_mean'].dropna()
        ax1.hist(pi_intensities_1, bins=250, alpha=alpha, color=pi_color, 
                density=True, label='Dead cells (PI positive)', edgecolor='darkred', linewidth=0.5)
    
    if 'cellevent_intensity_mean' in h2b_data_type1.columns:
        h2b_intensities_1 = h2b_data_type1['cellevent_intensity_mean'].dropna()
        ax1.hist(h2b_intensities_1, bins=250, alpha=alpha, color=h2b_color, 
                density=True, label='Alive cells (PI negative)', edgecolor='darkblue', linewidth=0.5)
    
    ax1.set_title(f'{celltype1}', fontsize=26, fontweight='bold')
    ax1.set_xlabel('CellEvent Intensity Mean', fontsize=20)
    ax1.set_ylabel('Normalized Frequency', fontsize=20)
    ax1.tick_params(axis='both', labelsize=16) 
    ax1.set_xlim([-1, 30])
    ax1.legend()
    ax1.grid(True, alpha=0.3)
    
    # Plot for second cell type
    celltype2 = unique_celltypes[1]
    pi_data_type2 = pi_data[pi_data['Cell Type'] == celltype2]
    h2b_data_type2 = h2b_data[h2b_data['Cell Type'] == celltype2]
    
    if 'cellevent_intensity_mean' in pi_data_type2.columns:
        pi_intensities_2 = pi_data_type2['cellevent_intensity_mean'].dropna()
        ax2.hist(pi_intensities_2, bins=250, alpha=alpha, color=pi_color, 
                density=True, label='Dead cells (PI positive)', edgecolor='darkred', linewidth=0.5)
    
    if 'cellevent_intensity_mean' in h2b_data_type2.columns:
        h2b_intensities_2 = h2b_data_type2['cellevent_intensity_mean'].dropna()
        ax2.hist(h2b_intensities_2, bins=250, alpha=alpha, color=h2b_color, 
                density=True, label='Alive cells (PI negative)', edgecolor='darkblue', linewidth=0.5)
    
    ax2.set_title(f'{celltype2}', fontsize=26, fontweight='bold')
    ax2.set_xlabel('CellEvent Intensity Mean', fontsize=20)
    ax2.set_ylabel('Normalized Frequency', fontsize=20)
    ax2.tick_params(axis='both', labelsize=16) 
    ax2.set_xlim([-1, 30])
    ax2.legend()
    ax2.grid(True, alpha=0.3)
    ax1.legend(fontsize=20)
    ax2.legend(fontsize=20)

    plt.tight_layout()
    
    print(f"\nData Summary:")
    print(f"Cell Type 1 ({celltype1}):")
    if 'cellevent_intensity_mean' in pi_data_type1.columns:
        print(f"  PI - Count: {len(pi_intensities_1)}, Mean: {pi_intensities_1.mean():.2f}, Std: {pi_intensities_1.std():.2f}")
    if 'cellevent_intensity_mean' in h2b_data_type1.columns:
        print(f"  H2B - Count: {len(h2b_intensities_1)}, Mean: {h2b_intensities_1.mean():.2f}, Std: {h2b_intensities_1.std():.2f}")
    
    print(f"Cell Type 2 ({celltype2}):")
    if 'cellevent_intensity_mean' in pi_data_type2.columns:
        print(f"  PI - Count: {len(pi_intensities_2)}, Mean: {pi_intensities_2.mean():.2f}, Std: {pi_intensities_2.std():.2f}")
    if 'cellevent_intensity_mean' in h2b_data_type2.columns:
        print(f"  H2B - Count: {len(h2b_intensities_2)}, Mean: {h2b_intensities_2.mean():.2f}, Std: {h2b_intensities_2.std():.2f}")
    
    return fig, (ax1, ax2)

# %% [code]
for i in range (1,3):
    dead = pd.read_csv(SOURCE_DATA_DIR/f"Imaging/Figure 7C/rep{i}_dead.csv")
    alive = pd.read_csv(SOURCE_DATA_DIR/f"Imaging/Figure 7C/rep{i}_alive.csv")
    fig, axs = compare_intensity_by_celltype(alive, dead)
    fig.savefig(f"Figures/Figure 7/rep{i}.svg")
```
