## Supplementary Data S2 for "Type III interferons may suppress viral infections by triggering cell death": Figure S1C plot RT-PCR.ipynb

```
# %% [code]
from pathlib import Path

import numpy as np
import pandas as pd
import scipy.stats as ss

import matplotlib.pyplot as plt
import seaborn as sns

from subplots_from_axsize import subplots_from_axsize

from utils import p_stars
from r_stats import r_anova_and_tukey

# %% [code]
plt.rcParams["font.size"] = 7

# %% [code]
SOURCE_DATA_DIR = Path('./Quantifications')

# %% [code]
!mkdir -p "Figures/Figure S1"

# %% [markdown]
# Figure S1C

# %% [code]
def load_and_normalize(path, delta_to='GAPDH', drop_house_keeping=['GAPDH', '18S', 'HPRT1']):
    data = pd.read_csv(path)
    data['Detector'] = data['Detector'].apply(lambda n: n.split('-')[0])
    data = data.set_index(['Detector', 'Experiment'])
    data = data - data.loc['GAPDH']
    data = data[~data.index.get_level_values('Detector').isin(drop_house_keeping)]
    return data.reset_index()

# %% [code]
data_dir = SOURCE_DATA_DIR / 'RT-PCR/Figure S1C'

for path_rep in sorted(data_dir.glob('rep*')):
    print(path_rep)

# %% [code]
data = pd.concat([
    load_and_normalize(path_rep)
    for path_rep in sorted(data_dir.glob('rep*'))
])

# %% [code]
data_plot_bvk = pd.DataFrame()
data_plot_bvk['Detector'] = data['Detector']
data_plot_bvk['6h']  = data['ctrl'] - data['IFN-β 6h']
data_plot_bvk['10h'] = data['ctrl'] - data['IFN-β 10h']
data_plot_bvk['24h'] = data['ctrl'] - data['IFN-β 24h']

data_plot_lvk = pd.DataFrame()
data_plot_lvk['Detector'] = data['Detector']
data_plot_lvk['6h']  = data['ctrl'] - data['IFN-λ 6h']
data_plot_lvk['10h'] = data['ctrl'] - data['IFN-λ 10h']
data_plot_lvk['24h'] = data['ctrl'] - data['IFN-λ 24h']

data_plot_bvl = pd.DataFrame()
data_plot_bvl['Detector'] = data['Detector']
data_plot_bvl['6h']  = data['IFN-λ 6h']  - data['IFN-β 6h']
data_plot_bvl['10h'] = data['IFN-λ 10h'] - data['IFN-β 10h']
data_plot_bvl['24h'] = data['IFN-λ 24h'] - data['IFN-β 24h']

# order by β > ctrl
detector_order = [
    'IFIT2',
    'IFIT1',
    'IFIH1',
    'ISG15',
    'IRF7',
    'DDX58',
    'STAT2',
    'OAS1',
    'CCL5',
    'SOCS1',
    'STAT1',
    'EIF2AK2',
    'RNASEL',
    'NFKBIA',
    'IL6',
    'TNFAIP3',
]

# panel 1: β > ctrl, mean
data_panel_1 = data_plot_bvk.groupby('Detector').mean()
data_panel_1 = data_panel_1.loc[detector_order]

# panel 2: λ > ctrl, mean
data_panel_2 = data_plot_lvk.groupby('Detector').mean()
data_panel_2 = data_panel_2.loc[detector_order]

# %% [code]
# panel 3: β > λ, p-stars
data_test = data.melt(
    id_vars=['Detector', 'Experiment'],
    value_vars=['IFN-β 6h', 'IFN-β 10h', 'IFN-β 24h', 'IFN-λ 6h', 'IFN-λ 10h', 'IFN-λ 24h'],
    value_name='dCT',
    var_name='Condition',
)

data_test['Time'] = data_test['Condition'].apply(lambda c: c.split()[-1])
data_test['Stimulation'] = data_test['Condition'].apply(lambda c: c.split()[0])
data_test['Stimulation'] = data_test['Stimulation'].apply(lambda c: {'IFN-β': 'IFNbeta', 'IFN-λ': 'IFNlambda'}[c])

ps_anova = []
for detector in detector_order:
    result_anova, _result_tukey = r_anova_and_tukey(
        'dCT ~ Stimulation * Time + Experiment',
        ['Stimulation', 'Time', 'Experiment'],
        'Stimulation',
        data_test[data_test['Detector'] == detector],
    )
    
    p_anova = result_anova.loc['Stimulation']['Pr(>F)']
    ps_anova.append(p_anova)

ps_bh_anova = ss.false_discovery_control(ps_anova, method='bh')
data_panel_3 = pd.DataFrame({'β > λ': map(p_stars, ps_bh_anova)}, index=detector_order)

# %% [code]
fig, (ax_panel_1, ax_panel_2, ax_panel_3, ax_cbar) = subplots_from_axsize(
    nrows=1, ncols=4,
    axsize=([0.45, 0.45, 0.15, 0.15], 0.15 * len(detector_order)),
    wspace=[0.2, 0.25, 0.25],
    left=0.55, bottom=0.1, top=0.75, right=0.4,
)

vrange=dict(vmin=-2, vmax=+8, center=0)

sns.heatmap(
    data_panel_1,
    ax=ax_panel_1,
    cmap='coolwarm',
    **vrange,
    cbar_ax=ax_cbar,
)
ax_panel_1.annotate('IFN-β\nvs\nControl', (0.5, 1.22), xycoords='axes fraction', ha='center', va='center', fontsize=8)

sns.heatmap(
    data_panel_2,
    ax=ax_panel_2,
    cmap='coolwarm',
    **vrange,
    cbar=False,
    yticklabels=False,
)
ax_panel_2.annotate('IFN-λ1\nvs\nControl', (0.5, 1.22), xycoords='axes fraction', ha='center', va='center', fontsize=8)

sns.heatmap(
    data_panel_3.applymap(lambda _: 1e3),
    cmap='gray',
    annot=data_panel_3,
    fmt='s',
    vmin=0, vmax=1,
    ax=ax_panel_3,
    cbar=False,
    xticklabels=False,
    yticklabels=False,
)
ax_panel_3.annotate('IFN-β\nvs\nIFN-λ1', (0.5, 1.22), xycoords='axes fraction', ha='center', va='center', fontsize=8)

for ax in [ax_panel_1, ax_panel_2, ax_panel_3]:  
    ax.set_ylabel('')
    ax.tick_params(top=True, labeltop=True, bottom=False, labelbottom=False)
    ax.tick_params(axis='x', labelrotation=90) 
    for lbl in ax.get_yticklabels():
        lbl.set_style('italic')

ax_cbar.set_ylabel('log₂ fold-change')

# IL6 and TNFAIP3 are not ISGs
ax_panel_1.get_yticklabels()[-2].set_color("grey")
ax_panel_1.get_yticklabels()[-1].set_color("grey")
    
fig.savefig('./Figures/Figure S1/figure_s1c--RT-PCR.svg', transparent=True)

# %% [code]
```
