## Supplementary Data S2 for "Type III interferons may suppress viral infections by triggering cell death": Figure S7B Western blot.ipynb

```
# %% [code]
import sys
from pathlib import Path

# Add the 'functions' directory to sys.path
sys.path.append(str(Path.cwd().parent.parent / "functions"))

from pathlib import Path

import numpy as np
import pandas as pd

import scipy.stats as ss

import matplotlib as mpl
import matplotlib.pyplot as plt
import matplotlib.patches as mpatches
from matplotlib.lines import Line2D

from subplots_from_axsize import subplots_from_axsize

from utils import load_rep, p_stars
from r_stats import r_anova_and_tukey
from blot_utils import *

# %% [code]
plt.rcParams["font.size"] = 8
color_ns = 'grey'

# %% [code]
!mkdir -p "Figures/Figure S7"


# %% [markdown]
# blot manipulation utils

# %% [code]
def blot_renorm_max(data, cutoff=0.03):
    data_norm = data.copy().set_index(['Experiment', 'Lane'])

    for col in data_norm:
        if data_norm[col].dtype=='object':
            continue

        data_norm[col] = data_norm[col] / data_norm[col].groupby(level=0).max()
        data_norm[col] += cutoff
        data_norm[col] = data_norm[col] / data_norm[col].groupby(level=0).max()

    return data_norm.reset_index()

# %% [markdown]
# blot plotting utils

# %% [code]
def plot_groups(
    axs,
    start_ends=[],
    margin=0.25,
    **kwargs
):
    for ax in axs:
        ax.spines['bottom'].set_color('lightgray')
        kwargs = dict(color='black', lw=1, clip_on=False, zorder=10) | kwargs
        for start, end in start_ends:
            ax.hlines(0, start-margin, end+margin, **kwargs)

# %% [code]
mol_to_ylabel = {
    "ns1": "NS1",
    "np.": "IAV NP",
}

def significance_line(ax, x1, x2, y, h=0.2, text="", color='k', lw=1.5, fontsize=10):
    """Draw a small bracket between x1 and x2 with optional text."""
    ax.plot([x1, x1, x2, x2], [y, y+h, y+h, y], lw=lw, c=color, clip_on=False)
    if text:
        ax.text((x1+x2)/2, y+h, text, ha='center', va='bottom', color=color, fontsize=fontsize, clip_on=False)


def plot_blots(
    data,
    exps=None,
    mols=None,
    mol_to_ylim_max=None,
    scalex=1.0,
    scaley=1.0,
    scalebarwidth=0.2,
    bar_spacing=0.2,
    group_spacing=1.0,
    margin=0.2,
    exp_to_color=None,
    **subplots_kwargs,
):
    """
    Plots bars grouped by Cell Type (preserving data order).
    Returns: fig, axs, group_centers
      - group_centers: dict mapping mol -> {cell_type: center_x}
    """
    exps = exps or data["Experiment"].unique()
    mols = mols or ["ns1", "np."]
    mol_to_ylim_max = mol_to_ylim_max or {}
    exp_to_color = exp_to_color or {}

    fig, axs = plt.subplots(
        nrows=len(mols),
        ncols=1,
        figsize=(scalex * 2.5 * len(data["Cell Type"].unique()), scaley * 2.5 * len(mols)),
        **subplots_kwargs,
    )

    if len(mols) == 1:
        axs = [axs]

    group_centers = {} 

    for ax, mol in zip(axs, mols):
        ax.set_ylabel(mol_to_ylabel.get(mol, mol), fontsize=9)
        ylim_max = mol_to_ylim_max.get(mol, data[mol].max() * 1.3)
        ax.set_ylim(0, ylim_max)

        xpos = 0.0
        xticks = []
        group_labels = []
        group_positions = []
        cell_type_ranges = []

        group_max = {}

        for cell_type, df_cell in data.groupby("Cell Type", sort=False):
            group_start = xpos

            for exp in exps:
                df_exp = df_cell[df_cell["Experiment"] == exp]
                if df_exp.empty:
                    continue

                vals = df_exp[mol].values
                n = len(vals)
                positions = xpos + np.arange(n) * bar_spacing

                color = exp_to_color.get(exp, 'k')
                ax.bar(
                    positions,
                    vals,
                    color=color,
                    width=scalebarwidth,
                    zorder=2,
                )

                xticks.append(positions.mean())

                xpos = positions[-1] + 1.0


            center = (group_start + xpos - group_spacing) / 2
            group_labels.append(cell_type)
            group_positions.append(center)
            cell_type_ranges.append((group_start, xpos - 1.0))
            group_max[cell_type] = df_cell[mol].max()
            xpos += group_spacing


        ax.set_xticks(xticks, labels='', fontsize=6)
        ax.spines[["top", "right"]].set_visible(False)


        sec = ax.secondary_xaxis(location=0)
        sec.set_xticks(group_positions, labels=group_labels, fontsize=9, fontweight="bold")
        sec.tick_params(axis="x", length=0, pad=25)

        ax.spines['bottom'].set_color('none')
        for start, end in cell_type_ranges:
            ax.hlines(
                0,
                start - margin,
                end + margin,
                color='black',
                linewidth=1,
                zorder=10,
                clip_on=False,
            )

        group_centers[mol] = dict(zip(group_labels, group_positions))

        group_centers.setdefault('_max', {})[mol] = group_max


    return fig, axs, group_centers


def plot_p_value(
    ax,
    p,
    pos_x1,
    pos_x2,
    height,
    yscale=1.33,
    thickness=0.05,
    group_line=None,
    always_plot_frame=True,
):
    s = p_stars(p)  # assume p_stars is defined elsewhere

    if not always_plot_frame and not s:
        return

    h0 = (0 * thickness + height) * yscale
    h1 = (1 * thickness + height) * yscale
    h2 = (2 * thickness + height) * yscale
    color = 'k' if s else color_ns  # assume color_ns defined

    ax.plot(
        [pos_x1, pos_x1, pos_x2, pos_x2],
        [h0, h1, h1, h0],
        color=color,
        lw=0.75,
        clip_on=False,
    )

    ax.annotate(
        s,
        ((pos_x1 + pos_x2) / 2, h2),
        ha='center',
        va='bottom',
        fontsize=8,
        annotation_clip=False,
    )

    if not group_line:
        return

    for pos in [pos_x1, pos_x2]:
        ax.plot(
            [pos - group_line / 2, pos + group_line / 2],
            [h0, h0],
            color=color,
            lw=0.75,
            clip_on=False,
        )

def test_blot(data, mol):
    
    data_test = data.copy().rename({'Cell Type': 'Group'}, axis=1)
    
    
    data_test = data_test.groupby(['Group', 'Experiment'], sort=False).mean().reset_index()

    data_test['Measurement'] = data_test[mol]#)
    result_anova, result_tukey = r_anova_and_tukey("Measurement ~ Group+Experiment",
    ['Group', 'Experiment'],
    'Group',data_test)
    
    p_anova = result_anova.loc['Group']['Pr(>F)']
    ps_tukey = dict(result_tukey['p adj'])
    
    return p_anova, ps_tukey

def plot_tests(
    axs,
    mols,
    data,
    test_blot,
    group_centers=None,
    margin=0.1,
):
    """
    axs: list of axes returned by plot_blots (same order as mols)
    mols: list of molecule column names (order must match axs)
    data: full dataframe (will be filtered inside)
    test_blot: function(data, mol) -> (p_anova, ps_tukey_dict)
    group_centers: mapping returned by plot_blots (mol -> {group_label: center_x})
    """
    if isinstance(data, list):
        datas = data
    else:
        datas = len(mols) * [data]

    tests = [
        test_blot(d, mol)
        for d, mol in zip(datas, mols)
    ]

    ps_anova = [p for p, _ in tests]
    pss_tukey = [ps for _, ps in tests]
    ps_bh_anova = ss.false_discovery_control(ps_anova, method='bh')  # assume ss is available
    for ax, p_bh_anova, ps_tukey, mol in zip(axs, ps_bh_anova, pss_tukey, mols):
        if p_bh_anova >= 0.05:
            ax.annotate(
                'ANOVA not significant',
                (0.5, 1.2),
                xycoords='axes fraction',
                va='center',
                ha='center',
                color=color_ns,
                style='italic',
            )
            continue

        # group_centers must be provided
        if not group_centers or mol not in group_centers:
            raise ValueError("You must pass group_centers (returned by plot_blots) to plot_tests")

        centers = group_centers[mol]  # dict: group_label -> x center

        # compute per-group maxima (from the data we use for tests) to position brackets
        group_max = data.groupby('Cell Type')[mol].max().to_dict()
        heights=[]
        # iterate tukey pairwise results: keys expected like "A-B"
        for key, pval in ps_tukey.items():
            try:
                g1, g2 = key.split('-')
            except Exception:
                # if key format unexpected, skip
                continue

            if g1 not in centers or g2 not in centers:
                # skip comparisons that aren't in the plotted groups
                continue

            pos1 = centers[g1]
            pos2 = centers[g2]

            # compute base height: a bit above the taller bar from the two groups
            max_h = max(group_max.get(g1, 0), group_max.get(g2, 0))
            # add margin (so bracket is above bars and above bottom axis)
            if max_h + margin not in heights:
                height = max_h + margin
                heights.append(height)
            else:
                height = max_h + 3* margin
                heights.append(height)
            # draw bracket and stars/p-values
            plot_p_value(ax, pval, pos1, pos2, height, yscale=1)

    return None

# %% [markdown]
# Figure S7B: NS and NP IAV IFNLR IFNAR KO

# %% [code]
df = pd.read_csv("./Quantifications/Western blot/Figure S7")

# %% [code]
def test_blot_log(data, mol):
    
    data_test = data.copy().rename({'Cell Type': 'Group'}, axis=1)
    
    
    data_test = data_test.groupby(['Group', 'Experiment'], sort=False).mean().reset_index()

    data_test['Measurement'] = np.log(data_test[mol])
    result_anova, result_tukey = r_anova_and_tukey("Measurement ~ Group+Experiment",
    ['Group', 'Experiment'],
    'Group',data_test)
    
    p_anova = result_anova.loc['Group']['Pr(>F)']
    ps_tukey = dict(result_tukey['p adj'])
    
    return p_anova, ps_tukey
fig, axs, centers = plot_blots(blot_renorm_max(df), scalebarwidth=0.2, bar_spacing=0.3, group_spacing=0.5, exp_to_color={'rep2':'blue'})

# draw whatever group-level markers you have (plot_groups etc.)
plot_groups(axs)

# then plot tests using the same centers
plot_tests(axs, ["ns1", "np."], blot_renorm_max(df, cutoff=0.03), test_blot_log, group_centers=centers)
plt.savefig("Figures/Figure S7/S7B.svg")
```
