## Supplementary Data S2 for "Type III interferons may suppress viral infections by triggering cell death": Figure S9A and S9C histograms immunostaning, Wester blot.ipynb

```
# %% [code]
import numpy as np
import pandas as pd

# %% [code]
!mkdir -p "Figures/Figure S9"

# %% [markdown]
# A.

# %% [code]
df = pd.read_csv("./Quantifications/Western blot/Figure S9C/IAV_NP_levels.csv")

# %% [code]
import matplotlib.pyplot as plt
import seaborn as sns

# filter out nuc_id = 0
df = df[df['nuc_id'] != 0]

# treatment order for subplots
treatment_order = ["12h", "24h", "48h"]

# identify KO variants (exclude WT)
ko_variants = [ct for ct in df['CellType1'].unique() if ct != "A549 WT"]

# colors for WT and KO
palette = sns.color_palette("Set2", n_colors=2)  # WT + KO
color_map = {"A549 WT": palette[0]}

# loop through stocks and experiments
rep_i =1
for stock, stock_df in df.groupby(["stock", "experiment"]):
    for ko in ko_variants:
        if ko!='A549 IFNLR KO (old)':
            continue
        plot_df = stock_df[stock_df['CellType1'].isin(["A549 WT", ko])]

        fig, axes = plt.subplots(1, len(treatment_order), figsize=(16, 5), sharey=True, sharex=True)

        for ax, treatment in zip(axes, treatment_order):
            
            subdf = plot_df[plot_df["treatments"] == treatment]
            if subdf.empty:
                ax.set_visible(False)
                continue

            # assign KO color dynamically
            color_map[ko] = palette[1]

            for i, (ct, sdf) in enumerate(subdf.groupby("CellType1")):
                ct_label = ct.replace(' (old)', '').replace(' (new)', '').replace('A549', '')
                ax.hist(
                    sdf["nuc_IAV_mean"],
                    bins=150,
                    alpha=0.5,
                    label=ct_label,
                    density=True,
                    color=color_map[ct]
                )

                # percentage above threshold
                above = (sdf["nuc_IAV_mean"] > 5000).mean() * 100
                ax.text(
                    0.98,
                    0.6 - i*0.08,
                    f"{ct_label}: {above:.1f}%",
                    color=color_map[ct],
                    transform=ax.transAxes,
                    ha="right",
                    va="top",
                    fontsize=11,
                    fontweight="bold"
                )

            ax.set_title(f"{treatment}")
            ax.axvline(5000, linestyle="--", color="grey")
            ax.spines["top"].set_visible(False)
            ax.spines["right"].set_visible(False)
            ax.set_xlabel("Mean IAV in nucleus [a.u.]")
            ax.set_xlim([0, 20000])
            ax.set_ylim([0, 0.0002])
        plt.locator_params(axis='x', nbins=5)
        axes[0].set_ylabel("Density")


        #fig.suptitle(f"stock: {stock}, KO: {ko} vs WT", fontsize=16)
        axes[0].legend(title="Cell line", loc="upper right")
        plt.tight_layout()
        plt.savefig(f"Figures/Figure S9/Histogram_WT_vs_KO_time_rep_{rep_i}.svg", format="svg")
        rep_i+=1


# %% [markdown]
# C. 

# %% [code]
blots = pd.read_csv("./Quantifications/Western blot/Figure S9C/IAV_NP_levels.csv")

# %% [code]
blots["IAV NP level IFNLR-KO"] = "IAV NP level IFNLR-KO"/100+0.03
blots["IAV NP level WT"] = "IAV NP level WT"/100+0.03

# %% [code]
pivoted = blots.pivot(index="Experiment", 
                   columns="Time", 
                   values=["IAV NP level IFNLR-KO", "IAV NP level WT"])

pivoted["KO_ratio"] = pivoted["IAV NP level IFNLR-KO"]["48h"] / pivoted["IAV NP level IFNLR-KO"]["24h"]
pivoted["WT_ratio"] = pivoted["IAV NP level WT"]["48h"] / pivoted["IAV NP level WT"]["24h"]

pivoted["KO_vs_WT_ratio"] = pivoted["KO_ratio"] / pivoted["WT_ratio"]


from scipy import stats

ratios = pivoted["KO_vs_WT_ratio"].dropna()
tstat, pval = stats.ttest_1samp(np.log(ratios), popmean=0, alternative='greater')

print("t-stat:", tstat, "p-value:", pval)

# %% [code]
x = np.arange(len(ratios))

# Choose which bar index to color blue
blue_index = 13  

# Colors: all black except one blue
colors = ["black"] * len(ratios)
#colors[blue_index] = "blue"

plt.figure(figsize=(6,4))

# Barplot
plt.bar(x, np.log(ratios), color=colors, edgecolor="black")

# Dashed line at 1
plt.axhline(0, color="red", linestyle="--", linewidth=1)

# Style tweaks
plt.ylabel("Growth difference between IFNLR KO and WT \n in 24 h - 48 h p.i. interval", fontsize=12)
plt.xticks(x, [], rotation=45)
#plt.ylim(0, max(ratios) * 1.2)
plt.text(12, 0.9, "Δ>0 *", fontsize=15)

# Clean look (like your plots)
plt.box(False)
plt.tick_params(axis='x', length=0)  # hide x ticks

plt.tight_layout()
plt.savefig("Figures/Figure S9/blot_log_fold_diff.svg", format="svg")
```
