## Supplementary Data S2 for "Type III interferons may suppress viral infections by triggering cell death": r_stats.py

```
import subprocess
import tempfile

import pandas as pd


def _generate_TukeyHSD_R_script(
    file_input,
    file_output_anova,
    file_output_tukey,
    term,
    factors,
    tukey_factor,
):
    code = []
    code.append(f"data <- read.csv('{file_input}')")

    for factor in factors:
        code.append(f"data${factor} <- as.factor(data${factor})")
    
    code.append(f"anova_result <- aov({term}, data=data)")
    code.append(f"anova_summary <- summary(anova_result)[[1]]")
    code.append(f"write.csv(anova_summary, '{file_output_anova}')")

    code.append(f"tukey_hsd <- TukeyHSD(anova_result, '{tukey_factor}')")
    code.append(f"tukey_hds_frame <- as.data.frame(tukey_hsd${tukey_factor})")
    code.append(f"write.csv(tukey_hds_frame, '{file_output_tukey}')")

    code = '\n'.join(code)

    return code


def r_anova_and_tukey(
    term,
    factors,
    tukey_factor,
    data,
):
    with (
        tempfile.NamedTemporaryFile(mode='w') as file_r_script,
        tempfile.NamedTemporaryFile(mode='w') as file_input,
        tempfile.NamedTemporaryFile(mode='w') as file_output_anova,
        tempfile.NamedTemporaryFile(mode='w') as file_output_tukey,
    ):
        r_code = _generate_TukeyHSD_R_script(
            file_input.name,
            file_output_anova.name,
            file_output_tukey.name,
            term,
            factors,
            tukey_factor,
        )
        file_r_script.write(r_code)
        data.to_csv(file_input.name, index=False)
        file_r_script.flush()
        file_input.flush()
        subprocess.run(['Rscript', file_r_script.name])
        result_anova = pd.read_csv(file_output_anova.name, index_col=0)
        result_tukey = pd.read_csv(file_output_tukey.name, index_col=0)

    # R inserts random spaces in the index
    result_anova.index = [i.strip() for i in result_anova.index]
    return result_anova, result_tukey
```
