## Supplementary Data S2 for "Type III interferons may suppress viral infections by triggering cell death": readme.txt

Pythoon codes will generate figures and created new folders for them in the folder Figures.

To run the code "Figure 7C and 7D histograms live exp.ipynb" please first download and unpack data file 'quantifications-fig-7cd.zip' from Zenodo: 
https://doi.org/10.5281/zenodo.17122542
