## Supplementary Data S2 for "Type III interferons may suppress viral infections by triggering cell death": utils.py

```
import pandas as pd
import scipy.stats as ss

import matplotlib.pyplot as plt
import matplotlib.colors as mcolors


plt.rcParams["font.family"] = ['arial', 'sans-serif']
plt.rcParams["font.size"] = 10


# convert Duration columns to time, unpack tuples
def load_rep(path):
    data = pd.read_csv(path)
    
    for col in data:
        if 'Duration' in col:
            data[col] = pd.to_timedelta(data[col])
    
    data.rename(columns={
        col_str: tuple(col_str[1:-1].replace('\'', '').split(', '))
        for col_str in data
        if '(' in col_str 
    }, inplace=True)
    
    return data


# time cmap
time_norm = mcolors.Normalize(vmin=0, vmax=5.0)  #
time_cmap = plt.cm.turbo
time_cmap_normed = lambda c: time_cmap(time_norm(c))

moi_to_marker = {
    0.01: 's',
    0.1:  '^',
    1.0:  'o',
}

def p_stars(p):
    star = '∗'  # unicode asterisk operator U+2217
    if p < 0.01:
        return '$' + 2 * star + '$'
    elif p < 0.05:
        return '$' + 1 * star + '$'
    else:
        return '!'
```
